# Trial-level Representational Similarity Analysis

**DOI:** 10.1101/2025.03.27.645646

**Authors:** Shenyang Huang, Cortney M. Howard, Paul C. Bogdan, Ricardo Morales-Torres, Matthew Slayton, Alex Clarke, Roberto Cabeza, Simon W. Davis

## Abstract

Neural representation refers to the brain activity that stands in for one’s cognitive experience, and in cognitive neuroscience, a prominent method of studying neural representations is representational similarity analysis (RSA). While there are several recent advances in RSA, the classic RSA (cRSA) approach examines the structure of representations across numerous items by assessing the correspondence between two representational similarity matrices (RSMs): usually one based on a theoretical model of stimulus similarity and the other based on similarity in measured neural data. However, because cRSA cannot weigh the contributions of individual trials (RSM rows/columns), it is fundamentally limited in its ability to assess subject-, stimulus-, and trial-level variances that all influence representation. Here, we formally introduce trial-level RSA (tRSA), an analytical framework that estimates the strength of neural representation for singular experimental trials and evaluates hypotheses using multi-level models. First, we verified the correspondence between tRSA and cRSA in quantifying the overall representation strength across all trials. Second, we compared the statistical inferences drawn from both approaches using simulated data that reflected a wide range of scenarios. Compared to cRSA, the multi-level framework of tRSA was both more theoretically appropriate and significantly sensitive to true effects. Third, using real fMRI datasets, we further demonstrated several issues with cRSA, to which tRSA was more robust. Finally, we presented some novel findings of neural representations that could only be assessed with tRSA and not cRSA. In summary, tRSA proves to be a robust and versatile analytical approach for cognitive neuroscience and beyond.

## Introduction

Since the start of any systematic examination of the mind, the concept of *representation* has provided a link between the external world and the content of mental life (Brentano, 1874). Current perspectives in cognitive science have defined representations as brain activity patterns that convey some behaviorally relevant content, which could be sensory perception, memory, concept knowledge, or social relations (Kriegeskorte & Diedrichsen, 2019). One of the central approaches for evaluating information represented in the brain is representational similarity analysis (RSA), an analytical approach that queries the representational geometry of the brain in terms of its alignment with the representational geometry of some cognitive model (Kriegeskorte et al., 2008; Kriegeskorte & Kievit, 2013), or, in some cases, compares the representational geometry of two neural systems (e.g., Kriegeskorte et al., 2008) or two model systems (Sucholutsky et al., 2025). The RSA approach has demonstrated utility across many domains of cognitive neuroscience research, such as visual perception (Jozranjbar et al., 2023), episodic memory (Xue, 2018), concept knowledge (Bauer & Just, 2019), social information (Freeman et al., 2018), and cognitive control (Freund, Etzel, et al., 2021). Despite its proven success, classic RSA approach, henceforth cRSA, has several limitations in efficacy across various experimental scenarios, due to its inability to reflect the proper multi-level variance structure within the data. Recently, methodological advancements have addressed many known limitations in cRSA. For example, cross-validated distance measures have improved the reliability of representational dissimilarities in the presence of noise and trial imbalance (Diedrichsen et al., 2021; Nili et al., 2014; Walther et al., 2016). Bayesian approaches such as pattern component modeling (Diedrichsen et al., 2018) have extended representational approaches to accommodate continuous stimulus features or temporal variation. Further, model comparison RSA strategies (Diedrichsen et al., 2021) and generalization techniques across stimuli and participants (Schütt et al., 2023) have improved sensitivity and inference. Nevertheless, a common feature shared across most of improvements is that they require stimuli repetition to separate estimates of noise from the underlying neural activity patterns. This requirement limits their ability to probe brain-behavior questions at the level of individual events. In this paper, we present an advancement termed trial-level RSA, or tRSA, which addresses these limitations in cRSA and may be utilized in paradigms with or without repeated stimuli.

A typical implementation of cRSA involves four main steps (see **Figure 1**). First, brain activity responses to a series of *N* trials are compared against each other (typically using Pearson’s r) to form an *N×N* representational similarity matrix, or RSM_brain_. Second, a hypothesis of how this brain system ought to respond — a cognitive model — is created in the form of a model RSM, or RSM_model_. This RSM_model_ can be created based on objective features of the stimuli (e.g., image brightness, category membership), subject ratings or behaviors (e.g., pleasantness, memory success), or outputs from computational models (e.g., neuron activations in artificial neural networks, semantic embeddings from large language models). Third, values from the lower (or equivalently, upper) triangular parts of both RSM_brain_ and RSM_model_ are retrieved, vectorized, and compared. This RSM_brain_-RSM_model_ comparison focuses on the similarity between the two representational geometries, which is typically measured by Spearman’s rank correlation (Kriegeskorte et al., 2008) but alternatives have also been proposed (Bobadilla-Suarez et al., 2020; Diedrichsen et al., 2021; Walther et al., 2016). This similarity measure is often referred to as a first-level RSA score or *representational strength^1^*. Fourth, once the above procedures are completed for each subject and condition, the first-level measures of representation are submitted to a second-level hypothesis testing for statistical inferences — often using general linear models such as t-test and ANOVA. One notable feature of cRSA is that the representational geometries are compared to one another in their entireties in step 3, and this step produces as a single measure of representational strength collapsing across all experimental trials. In other words, cRSA estimates cannot be interpreted for specific experimental trials or stimuli; rather, they only reflect how strongly the studied neural system represents for the entire set of items, limiting the realm of possible research questions. Most critically, as trial-level information is collapsed in cRSA, multiple sources of variance become intractable, yet these ignored variances could render subsequent second-level analyses susceptible to erroneous inferences.

**Figure 1.**
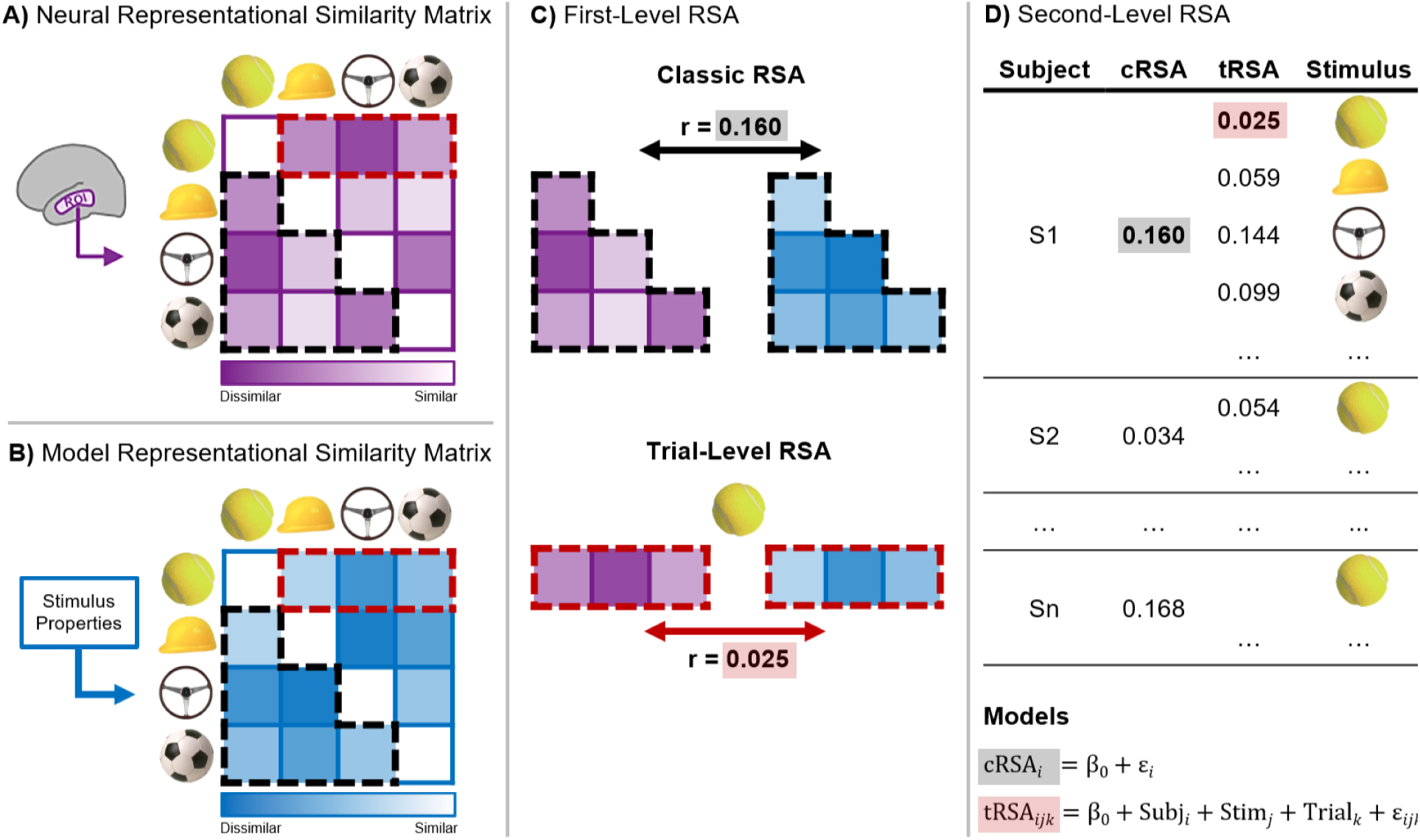
Representational Similarity Analysis (RSA). Steps for classic RSA (cRSA) and trial-level RSA (tRSA). **A)** Neural Representational Similarity Matrix (RSM_brain_) is generated by correlating multi-voxel activity patterns across all trials within a Region of Interest (ROI), reflecting similarity between neural responses. **B)** Model Representational Similarity Matrix (RSM_model_) is constructed by correlating of-interest stimulus properties across all trials. **C)** First-Level cRSA (top) and tRSA (bottom). For cRSA, the lower triangular parts (black outline) of RSM_brain_ and RSM_model_ are compared, producing a single summary statistic (e.g., Spearman’s rho) across all trials. For tRSA, representational similarity values from RSM_brain_ and RSM_model_ for the same trial (e.g., tennis ball; red dashed outline) are compared, producing a single representational strength estimate for that trial. **D)** Second-Level analysis for cRSA and tRSA. In cRSA, subject-level r values are submitted to a one-sample t-test to assess whether the values reliably exceed zero. In tRSA, a linear random effects model with random effects for subject and stimulus is fit, and hypothesis testing determines whether the estimated intercept is significantly greater than zero.

A behavioral or neural measure in a single experimental trial consists of meaningful variances from four distinct sources: *condition-level* (experimental task manipulations such as cognitive load and emotional valence)*, subject-level* (individual differences in perceptual acuity, prior knowledge, or other cognitive faculties)*, stimulus-level* (features of the stimuli with behavioral or neural relevance), and *trial-level* (physiological and nuisance variables affecting measurement). Notably, the distinction between condition-level and stimulus level is not always clear as researchers may manipulate stimulus-level features themselves. In these cases, what researchers ultimately consider condition-level and stimulus-level will depend on their specific research questions. For example, researchers intending to study generalized object representation may consider object category a stimulus-level feature, while researchers interested in if/how object representation varies by category may consider the same category variable condition-level.

To explain the variance in behavioral or neural measures, the appropriate statistical model should reflect said multi-level variance structure. However, the cRSA approach only does so imperfectly with three major limitations, which we detail below.

First, the heterogeneity of subject-and/or condition-level variances in real datasets adversely impacts the sensitivity and reliability of cRSA. One of the fundamental assumptions of general linear models (step 4 of cRSA; see **Figure 1D**) is *homoscedasticity* or homogeneity of variance — that is, all residuals should have equal variance. This assumption is often violated in real datasets. The variance of first-level cRSA scores depends largely on the number of observations (see **Figure 2**), but this trial count can be highly variable across subjects or conditions, resulting in *heteroscedasticity*. This issue is prevalent in studies where trials are selected and grouped based on subject behavior, such as in memory or attention tasks. In those cases, treating cRSA scores obtained from 100 trials and those obtained from 20 trials as having equal variance would violate the homoscedasticity assumption and lead to unreliable results. One suggested remedy of the issue is equating the number of trials via random subsampling conditions with more trials (Dimsdale-Zucker & Ranganath, 2018); however, this solution comes with the cost of not making full use of the rich information from those low-variance conditions and may not be ideal when the number of trials is highly unbalanced.

**Figure 2.**
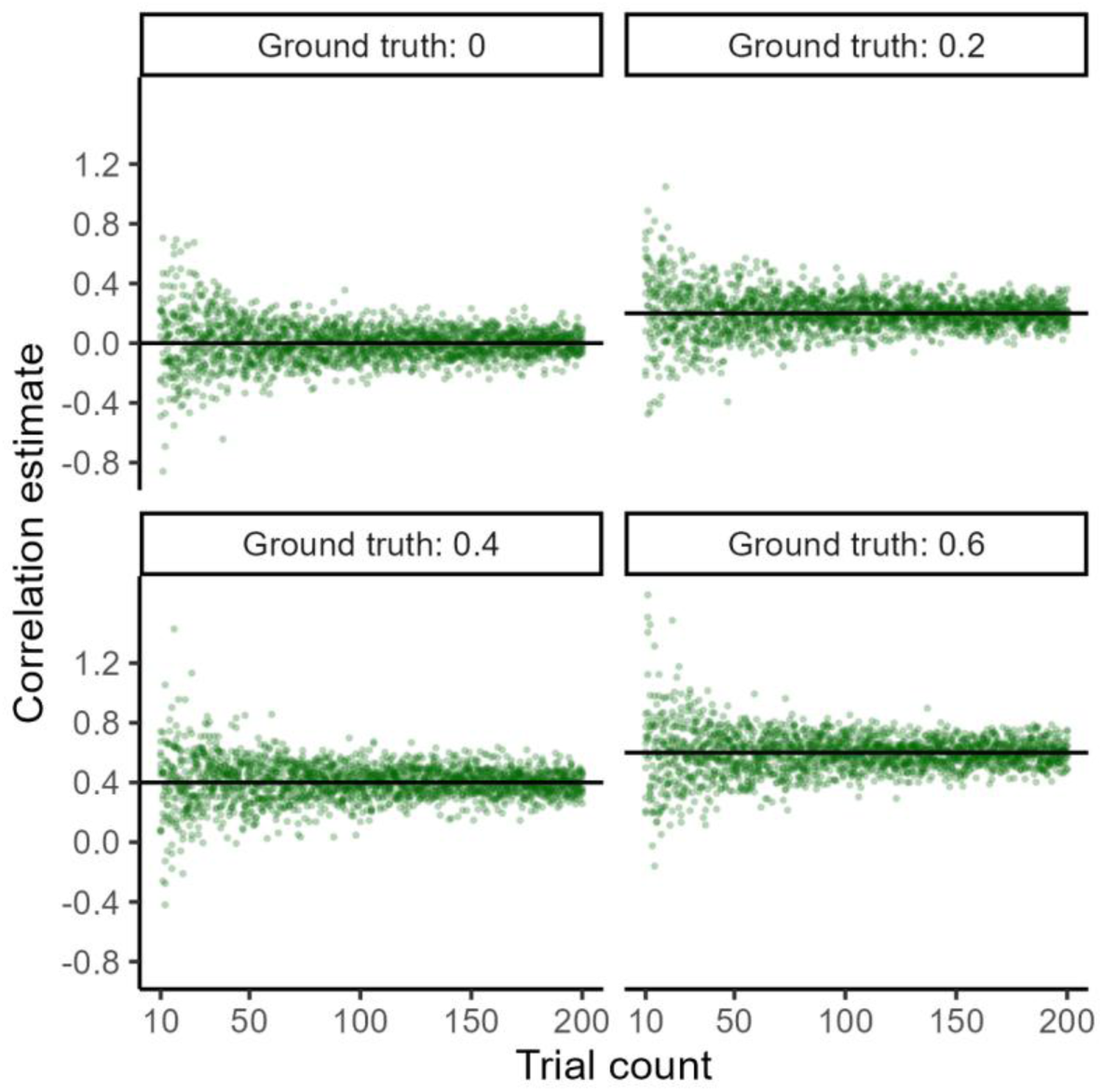
Trial count influences the reliability of correlations. The spread of correlation estimates (y-axis) varies nonlinearly with the number of observations, or trial count (x-axis). Trial count ranges from 10 to 200 with increments of 1. For each trial count, observations were drawn from a bivariate normal distribution with the given ground truth Pearson correlation, and their empirical Pearson correlation coefficient was computed. Ten random samples were drawn for each trial count and ground truth combination. Both ground truth and estimated values in the scatter plots are Fisher-transformed (z).

Second, cRSA is unable to model stimulus-level variance. Many studies present subjects with a fixed set of stimuli, which are supposedly samples representative of some broader category. In this case, using cRSA brings about two issues. For one, stimuli can vary in a wealth of properties. Object images, for example, vary in terms of image complexity, concept frequency, and memorability, all of which can affect both behavior and neural activity in important and systematic ways (Bainbridge et al., 2017; Hovhannisyan et al., 2021; Naspi et al., 2021). To mitigate this issue, one might explicitly manipulate the distribution of relevant stimulus properties or cross-validate cRSA results on subsets of stimuli (Freund, Bugg, et al., 2021). Nevertheless, neither solution could address the second issue, which is the fact that the same stimuli are presented to multiple subjects. To properly address this stimulus-level dependence in the data and generalize the results beyond the fixed stimulus set, one must model stimulus identity as random effects, for the very same reason subjects are specified as random effects (Chen et al., 2021; Yarkoni, 2022). However, this solution is not accessible for cRSA since its current implementation stipulates the collapsing of stimulus information across trials (step 3 of cRSA; see **Figure 1C**). While other approaches such as pattern component modeling(Diedrichsen et al., 2018) and encoding models (Naselaris et al., 2011) are well-suited to analyzing variables that vary continuously on a trial-by-trial or moment-by-moment basis, these frameworks address different inferential goals. Specifically, pattern component modeling and encoding models focus on estimating variance components or predicting activation from features, while cRSA is designed to evaluate representational geometry. Thus, cRSA as well as our proposed approach address a problem setting distinct from pattern component modeling and encoding models.

Third, cRSA is not well-suited for testing the influence of stimulus-level or trial-level properties on neural representations. For example, it may be desirable to explicitly test or control for how certain stimulus-level properties (e.g., memorability) or trial-level recordings (e.g., cardiac cycle, pupil dilation) affect neural representations (Critchley & Garfinkel, 2018; van der Wel & van Steenbergen, 2018). One strategy to analyze the effects of those stimulus-or trial-level continuous variables is to discretize them into categories (e.g., low vs. medium vs. high); however, discretization inevitably leads to a loss of information, reduced statistical power, and potentially misleading outcomes (Cohen, 1983; MacCallum et al., 2002). Alternatively, one could construct additional RSMs for the covariates and then compare all RSMs using a partial correlation or multiple regression framework, yet the statistical interpretations become much less straightforward after the conversion. For instance, even though a univariate random variable *v*, such as pleasantness ratings, can be conveniently converted to an RSM using pairwise distance metrics (Weaverdyck et al., 2020), the very same RSM would also be derived from the opposite random variable −*v*, leaving uncertain of the directionality (or if representation is strongest for pleasant or unpleasant items) of any findings with the RSM (see also Bainbridge & Rissman, 2018).

Here, we propose an original method for evaluating the neural representation of information at the level of individual experimental trials: *trial-level RSA* or *tRSA*. In this approach, instead of deriving a single measure of the similarity between RSM_model_ and RSM_brain_, we compute a series of similarity measures on a trial-by-trial (row-by-row or column-by-column) basis (Davis et al., 2021; King et al., 2019). For instance, the representational strength in the first trial is calculated as the similarity between the first row of RSM_model_ and the first row of RSM_brain_ (see **Figure 1C**). While several extensions of RSA have addressed key limitations in noise sensitivity, stimulus variance, and modeling (e.g., Diedrichsen et al., 2021; Schütt et al., 2023), our tRSA approach introduces a new methodological step by estimating representational strength at the trial level. This accounts for the multi-level variance structure in the data, affords generalizability beyond the fixed stimulus set, and allows one to test stimulus-or trial-level modulations of neural representations in a straightforward way. Critically, in single-presentation designs, a “trial” refers to one stimulus presentation, and corresponds to a row or column in the RSM. In studies with repeated stimuli, these rows are often called “conditions” and may reflect aggregated patterns across trials. tRSA is compatible with both cases: whether rows represent individual trials or averaged trials that create “conditions”, tRSA estimates are computed at the row level.

It is important to clarify that the two components of tRSA (row-level estimation and mixed-effects inference) address distinct methodological limitations of cRSA. First, the row-level computation yields trial-or stimulus-specific representational strength estimates that are otherwise collapsed when the lower triangular entries of the RSM are vectorized and summarized into a single correlation coefficient. By retaining one representational strength estimate per trial (or per row of the RSM), tRSA enables representational strength to be modeled as a function of stimulus-level or trial-level properties without requiring discretization or aggregation. Second, the use of linear mixed-effects models addresses a separate inferential concern. When the same fixed set of stimuli is presented to multiple participants, variability attributable to stimulus identity cannot be modeled if representational strength is reduced to a single subject-level summary statistic. Retaining row-level estimates makes it possible to include both subjects and stimuli as a random effect, thereby allowing generalization beyond the specific sample of participants and items.

To illustrate, consider a memory experiment in which 100 objects are presented once per participant. Some objects may be inherently more memorable than others, independent of participant-level variability. Under cRSA, one might compute representational strength separately for remembered and forgotten trials, yielding one value per subject per condition. However, it would not be possible to test whether item-level object memorability itself predicts representational strength, nor to model item-level variability explicitly. In tRSA, each object receives its own representational strength estimate, allowing memorability to be entered as a continuous predictor in a mixed-effects model with random intercepts for both subject and stimulus. This framework permits direct testing of item-level hypotheses and supports appropriate generalization beyond the specific stimulus set used in the experiment.

We assessed the efficacy of tRSA in comparison with cRSA using data from simulations representing a wide range of possible experimental scenarios (Experiment 1) and data from an extant real fMRI study (Experiment 2). In both experiments, our results demonstrated enhanced sensitivity, robustness, and flexibility with analyzing neural representations using the tRSA approach. Data is available upon request to the corresponding author and our simulations and example tRSA code is available at https://github.com/ElectricDinoLab/Trial-level-RSA.

## Methods

### Experiment 1: cRSA and tRSA comparisons in simulated data

Experiment 1 consisted of three main parts. First, we validated the similarity between tRSA and cRSA results in assessing the general representational strength across all trials. In other words, we established that tRSA can carry out the main purpose of cRSA. Second, we compared the statistical inferences generated by each approach in a set of synthetic within-subject studies with two conditions. Third, we demonstrated tRSA’s unique capability of assessing continuous modulators of representational strength in a set of synthetic scenarios. A host of parameters were varied to ensure the robustness of the comparisons in all three parts, such as the trial count, subject sample size, and effect size. All analyses were done in R 4.4.1 (R Core Team, 2024).

#### Estimating overall representational strength

We simulated *activity patterns*, from which RSMs were derived. Specifically, assuming an experiment with *n* trials and *q* measurement channels (e.g., voxels), we sampled a set of *n* ⋅ *q* values from a univariate standard normal distribution N(0,1), and another set of *n* ⋅ *q* values from a univariate normal distribution N(0, *σ*^2^). Both vectors were rearranged into matrices of *n* rows and *q* columns, yielding a *ground truth activity* pattern and a *noise* pattern. The summation of these two patterns yielded a *measured* pattern. Two RSMs were then generated from the ground truth pattern (RSM_model_) and the measured pattern (RSM_brain_) using Pearson’s correlation, with which cRSA and tRSA were conducted separately. This set of procedures was repeated for 10,000 iterations with different random samples. The following parameter values were used for reported results: *q* = 500, *n* ∈ {10, 15, 20, 25, 30, 40, 50, 100}, *σ*^2^ ranging from 0.8 to 2.0 with increments of 0.2.

We next assessed the correspondence between tRSA and cRSA in the presence of discrete conditions, which is often seen in real studies. Assuming a simple experimental design with two conditions, A and B, consisting of *n_A_* and *n_B_* trials, respectively, we generated a ground truth activity pattern with *n_A_* + *n_B_* rows and *m* columns in the same way as before. Then, we generated two sets of noise patterns, which were controlled by parameters *σ_A_* and *σ_B_*, respectively, one for each condition. The measured pattern was again computed as the summation of ground truth and noise. cRSA was conducted separately for trials in each condition. Critically, tRSA was conducted in an *across-condition* fashion: we generated RSMs with all *n_A_* + *n_B_* trials, obtained *n_A_* + *n_B_* tRSA values as previously described, and then split the values into two sets based on the condition each trial belonged to. Finally, as before, we computed the average of tRSA values separately for Conditions A and B to allow comparisons with cRSA values. This set of procedures was repeated for 10,000 iterations with different random samples. We assessed the influence of three factors: sample size, balance, and noise level, with the following parameter values: *q* = 500, *n_A_*, *n_B_* ∈ {20,80,320}, *σ_A_*, *σ_B_* ∈ {1,2}.

Of note, tRSA could conceivably also be conducted in a *within-condition* fashion, whereby one would generate two separate sets of RSMs according to condition, i.e., one set of *n_A_* × *n_A_* RSMs for Condition A and another set of *n_B_* × *n_B_* RSMs for Condition B, and then compute the two sets of *within-condition* tRSA values separately. We hereby advocate for the use of *across-condition* tRSA — which we have used in previous studies (Davis et al., 2021; Howard et al., 2024; S. Huang, Howard, et al., 2024; Naspi et al., 2023) — over within-condition tRSA. Comparisons of the statistical properties of within-condition and across-condition tRSA approaches are discussed in **Appendix 2**.

#### Statistical inferences from tRSA and cRSA

##### Modeling discrete conditions

The essential theoretical advantage of tRSA over cRSA is that representational strength can be estimated at the level of experimental trials, which would allow us to properly capture the multi-level variance structure in the data. To demonstrate this benefit, four sources of variances were hypothesized: condition, subject, stimulus, and trial. Condition-level variance denotes the effect of experimental conditions or manipulations on representation, with each condition receiving its own noise-level parameter *s_cond_*_,*k*_. This is the focal effect of interest for this simulation. Additionally, subject-level variance denotes individual differences in the quality of representation across *m* subjects, with each subject receiving one’s own noise-level parameter *s_subj_*_,*i*_. A within-subject design is common in real experiments and is thus assumed here, where each subject received all experimental conditions. Stimulus-level variance denotes the diversity of stimuli that may result in some being represented more strongly than others, with each stimulus receiving its own noise level parameter *s_stim_*_,*j*_. The subject-level and stimulus-level variances were fully crossed, i.e., all subjects were assumed to view all stimuli exactly once (condition counterbalanced) during the experiment. Finally, trial-level variance denotes random fluctuations in the signal across the experiment and is controlled by a single noise-level parameter *s_trial_*. Therefore, for a given event of subject *i* responding to stimulus *j* in condition *k*, we computed its noise level as:

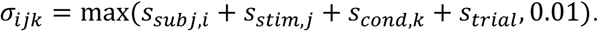

Notably, this multi-level variance structure replaced the fixed noise parameters *σ_A_*, *σ_B_*, while all other procedures remained the same as the previous simulation. The entire set of procedures was repeated for 10,000 iterations. The base simulation parameters were as follows:

- Experimental design: *q* = 500, *m* = 40, *n_A_* = *n_B_* = 100, and
- Multi-level variance: *s_cond_* ∈ {1,1.2}, *s_subj_*∼ N(0, 0.5^2^), *s_stim_* ∼ N(0, 0.2^2^), *s_trial_* = 2.

To assess the robustness of RSA methods in different situations that could occur in real data, we introduced a number of variations to the base design:

- Number of subjects: *m* ∈ {10, 20, 40, 80, 160, 320}.
- Trial count per subject: *n_trial_* ∈ {20, 30, 46, 70, 100, 250, 220, 330, 500}.
- Trial count ratio: *n_A_* ∈ {20, 40, 60, …, 180}, *n_B_* = 200 − *n_A_*.
- Variance in trial count ratio across subjects: *σ_n_* ∈ {0, 1, 2, 4, 8, 16, 32, 64, 128}. Actual trial counts were drawn from a truncated normal distribution,

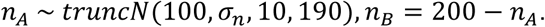
- Effect size: condition-level noise *s_cond_*_,*A*/*B*_ ∈ {1.00, 1.05, 1.10, …, 1.30}. Greater differences in the condition-level noise correspond to larger condition effects.

Following data generation, condition-level representational strength was estimated using cRSA and trial-level representational strength was estimated using tRSA. We focused on the statistical inferences of the effect of conditions, namely *b_B_*_−*A*_. To this end, cRSA estimates were submitted to a paired-sample t-test. Critically, tRSA estimates were submitted to a mixed-effects model which is statistically appropriate for modeling the hierarchical structure of the data, where observations are nested within both subjects and stimuli (Baayen et al., 2008; Chen et al., 2021). Specifically, a linear mixed-effects model with a fixed effect of condition (which estimates the average effect across the entire sample, capturing the overall effect of interest) and random effects of both subjects and stimuli (which model variations due to differences between individual subjects and items, allowing generalization beyond the sample) were fitted to tRSA estimates via the ‘lme4 1.1-35.3’ package in R (Bates et al., 2015), and p-values were estimated using Satterthwaites’s method via the ‘lmerTest 3.1-3’ package (Kuznetsova et al., 2017).

Given data generated with *s_cond_*_,*A*_ = *s_cond_*_,*B*_, the correct inference should be a failure to reject the null hypothesis of *b_B_*_−*A*_ = 0; any significant (*p* < 0.05) result in either direction was considered a false positive (spurious effect, or Type I error). Given data generated with *s_cond_*_,*A*_ >*s_cond_*_,*B*_, the inference was considered correct if it rejected the null hypothesis of *b_B_*_−*A*_ = 0 and yielded the expected sign of the estimated contrast (*b_B_*_−*A*_ > 0). A significant result with the reverse sign of the estimated contrast (*b_B_*_−*A*_ < 0) was considered a Type I error, and a nonsignificant (*p* ≥ 0.05) result was considered a false negative (failure to detect a true effect, or Type II error). Error rates were computed for each RSA method and were compared against the null hypothesis of equal proportions between methods.

##### Modeling continuous modulators

In addition to discrete experimental conditions, we also simulated data reflecting scenarios in which representational strength varies continuously with some modulator. For example, one may be interested in the effects of continuous variables such as image complexity and memorability (stimulus-level) or reaction time (trial-level). In this case, *s_trial_* was no longer a constant throughout the experiment but a trial-level variable. For a given event of subject *i* responding to stimulus *j* in trial *k*, we computed its noise level as:

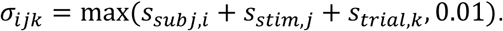

To capture the effect of this trial-level variable, we generated a random variable *v_measured_* that reflects a trial-level measurement with construct validity *r_val_* = *Cor*(*v_measured_*, *s_trial_*), which was restricted to be a nonpositive value. When *r_val_* = 0, *v_measured_* carries no information of the underlying *s_trial_* and should not predict trial-level representational strength. When *r_val_* < 0, *v_measured_* is inversely related to trial-level noise and should positively predict trial-level representational strength. The entire set of procedures was repeated for 10,000 iterations. The base simulation parameters were as follows:

- Experimental design: *q* = 500, *m* = 40, *n* = 200, and
- Variance structure: *r_val_* ∈ {0, −0.1}, *s_subj_*∼ N(0, 0.5^2^), *s_stim_* ∼ N(0, 0.2^2^), *s_trial_* = 2. As before, we introduced a number of variations, as follows:
- Number of subjects: *m* ∈ {10, 20, 40, 80, 160, 320}.
- Number of trials per subject: *n_trial_* ∈ {20, 30, 46, 70, 100, 250, 220, 330, 500}.
- Variance in the number of trials per subject: *σ_n_* ∈ {0, 20, 40, …, 160}. Actual trial counts were drawn from a truncated normal distribution, *n* ∼ *truncN*(*n*, *σ_n_*, 20, *n*).
- Effect size: *r_val_* ∈ {−0.10, −0.09, …, 0.00}. More negative values of *r_val_* should result in greater representational strength.

Following data generation, two conditions were created for each subject based on a median split (A=low *v_measured_*, B=high *v_measured_*), and we tested cRSA and tRSA models as described in the previous simulation, focusing on *b_B_*_−*A*_. Furthermore, we additionally modeled trial-level representational strength as a function of *v_measured_* directly, using a linear mixed-effects model of tRSA estimates with a fixed effect of *v_measured_* and random effects of both subjects and stimuli. Given data generated with *r_val_* = 0, the correct inference should be a failure to reject the null hypothesis of *b* = 0 in both discrete and continuous models; any significant (*p* < 0.05) result in either direction was considered a false positive or Type I error. Given data generated with *r_val_* < 0, the inference was considered correct if it rejected the null hypothesis of *b* = 0 and yielded the expected sign of the estimated contrast or slope *b* > 0. A significant result with the reverse sign of the estimated contrast or slope (*b* < 0) was considered a Type I error, and a nonsignificant (*p* ≥ 0.05) result was considered a false negative or Type II error. Error rates were computed for each RSA method and were compared against the null hypothesis of equal proportions between methods.

### Experiment 2: cRSA and tRSA comparisons in fMRI data

Experiment 2 proceeded in 3 steps. First, we assessed how the impact of subject-level variance in heterogeneity of trial ratios amongst two critical conditions (“Hits” and “Misses”) can lead to biased estimates of representational strength. Second, we examined the impact of modeling trial-level variance on both representational strength during Object Naming, as well as Mnemonic Strength during a memory retrieval task. Lastly, stimulus-level estimates of representational strength were related to object memorability — an analysis that is not directly accessible to cRSA.

#### Data acquisition

##### Participants

A pre-existing dataset was analyzed to evaluate tRSA. Main study findings have been reported elsewhere (S. Huang et al., 2025). A total of 38 adults, aged 18 to 30, participated in this study on a voluntary basis and received monetary compensation for their time. Eligibility criteria required participants to be native or fluent English speakers, with no history of significant neurological or psychiatric conditions, and not taking medications that could affect cognitive function or cerebral blood flow (except for antihypertensive agents). All participants provided written informed consent before the start of the study. Six participants did not complete the study and were excluded from the analysis. Additionally, two participants were excluded from the Memory Retrieval analyses due to poor memory performance. The final sample included 32 participants (21 women and 11 men) for the Object Perception task and 30 participants (19 women and 11 men) for the Memory Retrieval task.

##### Experimental design

An expansive outline of the study design for the tasks described below can be found in (S. Huang et al., 2025); here we briefly review relevant details pertinent to our application of trial-level RSA for a subset of that data. Data from two experimental datasets were used, including an Object Perception dataset and a Memory dataset (**Figures 7A and 8A**). In the Object Perception task, participants were shown images of 114 unique everyday objects on a white background. The corresponding label presented underneath each object and participants were asked to rate how well the label described the object on a four-point scale. The main purposes of this rating task was to make sure that participants assess the meaning of the objects and to verify that these objects are indeed familiar to them (mean rating = 3.59). The objects were presented for 4 seconds, followed by a jittered fixation cross with an average duration of 4 seconds. In the memory task, a Memory Encoding session took place at least 7 days after Object Perception. During Encoding, participants viewed images of 114 unique real-world scenes along with the 114 objects they had seen in session 1. In each trial, participants were shown a scene image for 3 seconds, followed by a jittered empty box indicating the continuation of the trial, which lasted for an average of 3 seconds, and finally an object image for 4 seconds. During the object presentation, participants rated “how likely it is to find the object in the scene” on a 4-point scale (1 = “very unlikely,” 2 = “somewhat unlikely,” 3 = “somewhat likely,” 4 = “very likely”). Each trial was separated by a jittered fixation cross with an average duration of 4 seconds. Memory Retrieval consisted of three scanning runs, each with 38 trials, lasting approximately 9 minutes and 12 seconds. Memory Retrieval consisted of three scanning runs, each with 38 trials, lasting approximately 9 minutes and 12 seconds (within-run comparisons were later excluded from RSA analyses). Memory Retrieval took place one day after Memory Encoding and involved testing participants’ memory of the objects seen in the Encoding phase. Neural data during the Encoding phase has been reported elsewhere. In the main Memory Retrieval task, participants were presented with 144 labels of real-world objects, of which 114 were labels for previously seen objects and 30 were unrelated novel distractors. Participants performed old/new judgements, as well as their confidence in those judgements on a four-point scale (1 = Definitely New, 2 = Probably New, 3 = Probably Old, 4 = Definitely Old). In a subsequent Perceptual Memory Retrieval task participants were shown 126 images of real-world objects (96 old images, 18 were different exemplar images of old objects, and 12 images of unrelated novel objects). Participants were asked to determine whether the image was old, similar, or new. All analyses in the current study pertained to fMRI data from Object Perception and the main Memory Retrieval tasks, as well as behavioral data from the Perceptual Memory Retrieval task.

##### MRI data acquisition

MRI data were collected using a 3T GE MR750 Scanner equipped with an 8-channel head coil at the Brain Imaging and Analysis Center (BIAC) at Duke University. Each MRI session began with a localizer scan, during which 3-plane (straight axial/coronal/sagittal) faster spin echo images were obtained. A high-resolution T1-weighted (T1w) structural scan was then acquired, consisting of 96 axial slices parallel to the AC-PC plane, with voxel dimensions of 0.9 × 0.9 × 1.9 mm³. This was followed by blood-oxygenation-level-dependent (BOLD) functional scans using a whole-brain gradient-echo echo planar imaging sequence (repetition time = 2000 ms, echo time = 30 ms, field of view = 192 mm, 36 oblique slices with voxel dimensions of 3 x 3 x 3 mm³). Task instructions and stimuli were delivered using the PsychToolbox program (Kleiner et al., 2007) and projected onto a mirror at the back of the scanner bore. Participants responded using a four-button fiber-optic response box. To reduce scanner noise, participants wore earplugs, and MRI-compatible lenses were provided when necessary to correct vision. Foam padding was placed inside the head coil to minimize head movement.

#### Data analysis

##### MRI data preprocessing

fMRIPrep 23.0.1 (Esteban et al., 2019, 2023) was used to preprocess structural and functional MRI data, as well as generating text descriptions of preprocessing details, which were condensed below. T1w structural images collected across all MRI sessions for the same participant were corrected for intensity non-uniformity with ‘N4BiasFieldCorrection’ (Tustison et al., 2010) from ANTs 2.3.3 (Avants et al., 2011). The T1w-reference was skull-stripped with a Nipype implementation of the ‘antsBrainExtraction.sh’ workflow from ANTs, using OASIS30ANTs as target template. Brain tissue segmentation of cerebrospinal fluid (CSF), white-matter (WM), and gray-matter (GM) was performed on the brain-extracted T1w using ‘fast’ from FSL (Zhang et al., 2001). An anatomical T1w-reference map was computed after registration of T1w images using ‘mri_robust_templatè from FreeSurfer 7.3.2 (Reuter et al., 2010). Brain surfaces were reconstructed using ‘recon-all’ from FreeSurfer 7.3.2 (Dale et al., 1999), and the brain mask estimated previously was refined with a custom variation of the method to reconcile ANTs-derived and FreeSurfer-derived segmentations of the cortical gray-matter of Mindboggle (Klein et al., 2017). Volume-based spatial normalization to the ICBM 152 Nonlinear Asymmetrical template version 2009c standard space was performed through nonlinear registration with ‘antsRegistration’ from ANTs 2.3.3, using brain-extracted versions of both T1w reference and the T1w template.

BOLD functional data across all sessions and runs were preprocessed collectively. A reference volume and its skull-stripped version were generated using a custom methodology of fMRIPrep. Head-motion parameters with respect to the BOLD reference were estimated, followed by spatiotemporal filtering using ‘mcflirt’ from FSL (Jenkinson et al., 2002). BOLD runs were slice-time corrected to 0.972s (0.5 of slice acquisition range 0s-1.94s) using ‘3dTshift’ from AFNI (Cox & Hyde, 1997). The BOLD time-series were resampled onto their original, native space by applying the transforms to correct for head-motion. The BOLD reference was then co-registered to the T1w reference using boundary-based registration via ‘bbregister’ from FreeSurfer (Greve & Fischl, 2009). Co-registration was configured with six degrees of freedom. Confounding time-series calculated based on the preprocessed BOLD included: root mean square displacement (RMSD) between frames (Jenkinson et al., 2002), absolute sum of relative framewise displacement (FD) (Power et al., 2014), and the derivative of root mean square variance over voxels (DVARS) (Power et al., 2014), as well as global signals extracted within CSF, WM, and the whole-brain mask. Additionally, a set of physiological regressors were extracted to allow for component-based noise correction (CompCor) (Behzadi et al., 2007). Principal components were estimated after high-pass filtering the preprocessed BOLD time-series (using a discrete cosine filter with 128s cut-off) for the two CompCor variants: temporal (tCompCor) and anatomical (aCompCor). tCompCor components were calculated from the top 2% variable voxels within the brain mask. For aCompCor, three probabilistic masks (CSF, WM and combined CSF+WM) were generated in anatomical space. For each CompCor decomposition, the k components with the largest singular values that cumulatively explained at least 50% of variance across the nuisance mask (CSF, WM, combined, or temporal) were retained. The BOLD time-series were resampled into standard space with a spatial resolution of 2 × 2 × 2 mm^3^ or 97 × 115 × 97 voxels. First, a reference volume and its skull-stripped version were generated using a custom methodology of fMRIPrep. All resamplings can be performed with a single interpolation step by composing all the pertinent transformations (i.e., head-motion transform matrices, susceptibility distortion correction when available, and co-registrations to anatomical and output spaces). Gridded (volumetric) resamplings were performed using ‘antsApplyTransforms’ (ANTs), configured with Lanczos interpolation to minimize the smoothing effects of other kernels (Lanczos, 1964). Non-gridded (surface) resamplings were performed using ‘mri_vol2surf’ (FreeSurfer).

##### Single-trial modeling

Neural activity in gray matter (GM) voxels for each object presentation event was estimated using first-level least squares separate general linear models (Mumford et al., 2012) constructed with SPM12 (Friston et al., 2006) and custom MATLAB scripts. Subject-specific GM masks were generated by binarizing the fMRIPrep-derived probabilistic masks with a threshold set to exclude voxels with 80% probability of CSF or WM. Each model included a regressor for the event of interest along with a regressor for all other objects. Both regressors were convolved with the canonical double-Gamma hemodynamic response function, including their temporal and dispersion derivatives to account for variations in the timing of the peak response.

Covariates of no interest included global signal, WM signal, CSF signal, FD, DVARS, RMSD, and six motion parameters related to translation and rotation. A high-pass temporal filter with a cutoff of 128 seconds was applied, and the AR(1) model was used to correct for autocorrelation. These first-level models yielded regression coefficients (betas) that estimated voxel-level neural activity corresponding to a single trial. Trials with FD greater than 1mm either before or during the trial were excluded (range: 0-28). Additionally, runs with more than 25% motion trials were excluded. In total, 1 Object Perception run, and 3 Memory Retrieval runs were excluded.

##### Behavioral Analyses

Trials for which participants gave no response in the Object Perception (mean 5) or the main Memory Retrieval (mean 4) were excluded from all analyses involving fMRI data at Perception and Memory Retrieval tasks respectively. As our primary behavioral measure, we assessed the memory performance for the object labels. Participants demonstrated variability in their ability to distinguish between old and new objects, as well as in their decision criteria, making direct comparisons of raw responses challenging. To address these biases, we performed a receiver-operating characteristics (ROC) analysis using ‘yardstick 1.3.1’ (Kuhn et al., 2024) in R. This analysis allowed us to categorize Hit and Miss trials by determining whether counting only “4” responses or both “3” and “4” responses as “old” yielded the best decision outcome (i.e., closest to 100% true positive and 0% false positive). The outcome of this analysis was then used to compute the sensitivity index (d’) of recognition. Adjusted hit rates were used to determine adherence to the task and sufficient fMRI data during Memory Retrieval.

Behavioral analyses categorized each trial during Memory Retrieval as either a “Hit” or “Miss” for the purpose of categorizing the conditions in the fMRI data. This analysis determined whether counting only “4” responses or both “3” and “4” responses as “old” produced the most accurate decision outcome (i.e., closest to 100% true positive and 0% false positive). In total, data from 8 participants were adjusted to consider only “4” responses as “hit” trials. The results were then used to calculate the sensitivity index (d’) of recognition within each subjective congruency condition and the corrected hit rate. Participants with post-adjustment hit rates below 40% were considered to have low adherence to the task and were excluded from all subsequent analyses. The number of included trials per participant for perception (mean = 110), ROC values (mean = 0.78), the count of hit trials (mean =76) as well as the corrected hit rate (mean = 0.70), are detailed in **Appendix 3 Table 4**.

##### Neural Representational Similarity Matrices

Models for neural pattern similarity (RSM_brain_) were constructed using both the Object Perception study and the main Memory Retrieval task. Twenty six regions from the Human Brainnetome Atlas (Fan et al., 2016), focusing on eight areas within the lateral occipital cortex (LOC), 14 areas within the inferior temporal cortex (ITC), and four areas in the inferior parietal cortex (IPL, combining IPL sub regions into lateral posterior and anterior regions), as these regions are critically involved in visual representations during object perception and memory retrieval (Davis et al., 2021; Favila et al., 2020; Howard et al., 2024; Long & Kuhl, 2021). For each of these 26 regions, we constructed similarity matrices of voxel activation patterns across stimuli with custom scripts in MATLAB and SPM12 (Friston et al., 2006). This was done by vectorizing the voxel-level activation values within each region and calculating their correlations using Pearson’s r, excluding all within-run comparisons. While each cell in a RSM_model_ reflects the similarity in stimulus properties, each cell in the 114×114 RSM_brain_ represents the similarity in activation patterns across different stimuli (**Figure 1A**).

##### Model Representational Similarity Matrix

The C2 layer of the Hierarchical Model of object recognition (HMAX) was used to capture visual similarity between the 114 objects. The PsyTorch implementation of the HMAX model was used and contains four sequential stages, each with their own output: S1, C1, S2, and C2. The S1 layer applies Gabor filters to the input image across multiple scales and orientations. The Gabor filters capture edge-like features such as bars and gratings. The output consists of feature maps that highlight these basic visual components. Following the S1 layer, the C1 layer performs local max pooling over the S1 feature maps. This operation introduces some degree of invariance to position and scale. By selecting the maximum response within localized regions, the C1 layer reduces the spatial resolution while retaining the most salient features. The stages S2 and C2 build upon this foundation using similar pooling mechanisms. Specifically, S2 units pool information from the C1 stage using linear filters and function as radial basis functions, responding most strongly to specific prototype input patterns. These prototypes are derived from random fragments extracted from a set of natural images, independent of the stimuli used in this study. The C2 layer then pools outputs from S2 units using a MAX operation, which provides a degree of position and scale tolerance, allowing for robust representation of visual objects. This global pooling results in a feature vector that is highly invariant to position and scale, capturing the presence of complex features regardless of their location in the input image. The C2 layer’s output serves as a compact representation of the visual input (Kriegeskorte et al., 2008; Riesenhuber & Poggio, 1999, 2002; Serre et al., 2005, 2007; Sufikarimi & Mohammadi, 2020).

Features from the C2 layer of the HMAX model were used to create the model RSM. Pairwise similarity between images was computed using Pearson’s correlation coefficient (r), measuring the relationship between their C2 feature vectors. These values were organized into a 114×114 RSM, where each cell represented the similarity between two images. The resulting RSM_model_ provided a structured representation of visual similarity, with higher correlations indicating shared complex features (**Figure 1B**).

##### Classic Representational Similarity Analysis (cRSA)

We conducted cRSA using data from both the Object Perception dataset and the Memory Retrieval dataset. For Object Perception, we estimated participants’ classic representational strength in each of the 26 regions of interest by calculating Spearman’s rho between the RSM_brain_ and the RSM_model_, while excluding within-run comparisons and the diagonal cells of the matrices (see **Figure 1C, Top**). We then assessed the strength and reliability of visual representations across participants using one-sample t-tests (testing Fisher-transformed rho > 0) for each of the 26 regions, applying a false discovery rate (FDR) correction with a threshold of q = 0.05.

To evaluate cRSA’s performance in detecting differences in representational strength between conditions, we analyzed data from the main Memory Retrieval task, focusing on old items. An RSM_brain_ for each of the 26 regions of interest was parsed into two separate RSMs: one containing data from “Hit” trials and the other from “Miss” trials. Similarly, the model RSMs were split based on participant responses. We then calculated Spearman’s rhos for the Hit and Miss conditions by correlating RSM_brain_ and RSM_model_, again excluding within-run comparisons and the diagonal cells of the matrices. These subject-specific representational strength measures were subsequently Fisher-transformed and analyzed using paired-sample t-tests. Regions showing significantly higher representational strength for Hit trials compared to Miss trials, after FDR correction with q < 0.05, were identified as *mnemonic representation regions*.

To assess the impact of unbalanced trial counts on mnemonic representational strength, we calculated a Contrast Variance Factor (CVF) for each participant in regions showing evidence of mnemonic representation (p < 0.05, uncorrected). The participant contrast score was determined by subtracting the Miss representational strength (*R_i_*_,*M*_) from the Hit representational strength (*R_i_*_,*H*_).

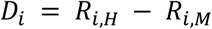

Next, we computed the mean difference *D^-^* across all participants:

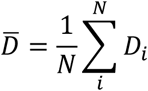

To quantify how much each participant’s mnemonic representational strength in each ROI deviated from the mean (thus inflating the contrast variance), we calculated the normalized absolute difference between the participant contrast score (*D_i_*) and the average contrast score (*D^-^*):

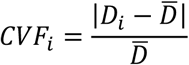

Finally, to evaluate the influence of unbalanced trial counts on the variance factor, we correlated the participant CVF averaged across regions with the ratio of Hit trial count (*N_i_*_,*H*_) to Miss trial count (*N_i_*_,*M*_).

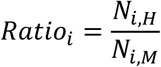

##### Trial-Level Representational Similarity Analysis (tRSA)

The tRSA approach used the same model and neural RSMs from the Object Perception dataset and the main Memory Retrieval dataset as described above. However, instead of correlating the two matrices in their entirety, correlations were computed on a row-by-row basis, excluding within-run similarities and the diagonal cells of the matrices (see **Figure 1C, Bottom**). The trial-level estimates from both phases were then fitted to a series of mixed-effects models using the ‘lme4’ package in R (R Core Team, 2024) with restricted maximum likelihood methods. These fitted models were then evaluated using the Akaike Information Criterion (AIC) to determine the optimal random effect structure (Meteyard & Davies, 2020; Park et al., 2020). To further validate model selection (Matuschek et al., 2017), models were refit using maximum likelihood estimation and subjected to model selection via log-likelihood ratio tests (LRTs). The average AICs and the results of the LRTs are reported in **Appendix 3 Table 5**. Restricted maximum likelihood models with the selected random effects structure were further analyzed using the ‘lmerTest’ package. Denominator degrees of freedom were estimated using Satterthwaite’s method (Satterthwaite, 1946), and fixed effects were tested using t-tests with an alpha level of 0.05, corrected for false discovery rate (FDR) to account for multiple comparisons.

To estimate tRSA representational strength, trial-level estimates from the Object Perception dataset were fit to a mixed-effects model with random intercepts for Participant and Stimulus. Estimated intercepts (b > 0) were used to identify representational regions. For detecting tRSA mnemonic representations in the main Memory Retrieval task, instead of constructing separate Hit and Miss RSMs, row-wise correlations were performed across all trials in the matrices (i.e., across-condition tRSA). This approach allows for trial-level estimates of representational strength that can be subsequently categorized as “Hit” or “Miss” and analyzed using a series of mixed-effects models. The model for mnemonic representation data included the fixed effect of Memory Success (Hit vs. Miss trials) and random intercepts for Participant and Stimulus.

##### Examining continuous modulators of representation

The capacity of tRSA in examining stimulus-level variance modulating representation was assessed. For each item, a memorability score was calculated as the average response across participants (on a scale of 1-4 for the main Memory Retrieval and 1-3 for Perceptual Retrieval), normalized by the maximum possible value (4 for main retrieval and 3 for perceptual retrieval). This approach produced two stimulus-specific continuous variables indicating the overall confidence with which each item would be recalled. We refer to these measures as Conceptual Memorability and Item Memorability (perceptual). The Item Memorability measure was used as an of-interest fixed effect for trial-level representational strength estimates from the Object Perception dataset. Additional fixed nuisance variables included Conceptual Memorability, fMRI run, and trial-level reaction time. Random intercepts for Participant and Stimulus were also included.

## Results

### Experiment 1: cRSA and tRSA comparisons in simulated data

#### Basic statistical properties of tRSA estimates

We simulated activity patterns from which RSMs were then derived and used to estimate representational strength. Notably, while it is difficult to fix the ground truth representational strength, we manipulated the statistical dependence between the activity patterns — and by extension, between RSM_brain_ and RSM_model_ — by injecting varying levels of measurement noise (*σ*^2^). Simulations indicated that increasing measurement noise indeed reduced estimates of representational strength by both cRSA and tRSA (see **Figure 3A**). Also as expected, greater trial counts led to more stable estimates for both methods. Most importantly, a strong positive correlation between cRSA and tRSA estimates can be observed even in the “noisiest” simulation (i.e., *n* = 10, *σ*^2^ = 2; Intercept b_0_ = 0.00, SE = 0.01, t = 0.33, p = 0.74; Slope b_cRSA_ = 1.00, SE = 0.02, t = 64.95, p < 0.001). These results demonstrated the close numerical correspondence between tRSA and cRSA in estimating the overall representational strength of a given system across all trials.

**Figure 3.**
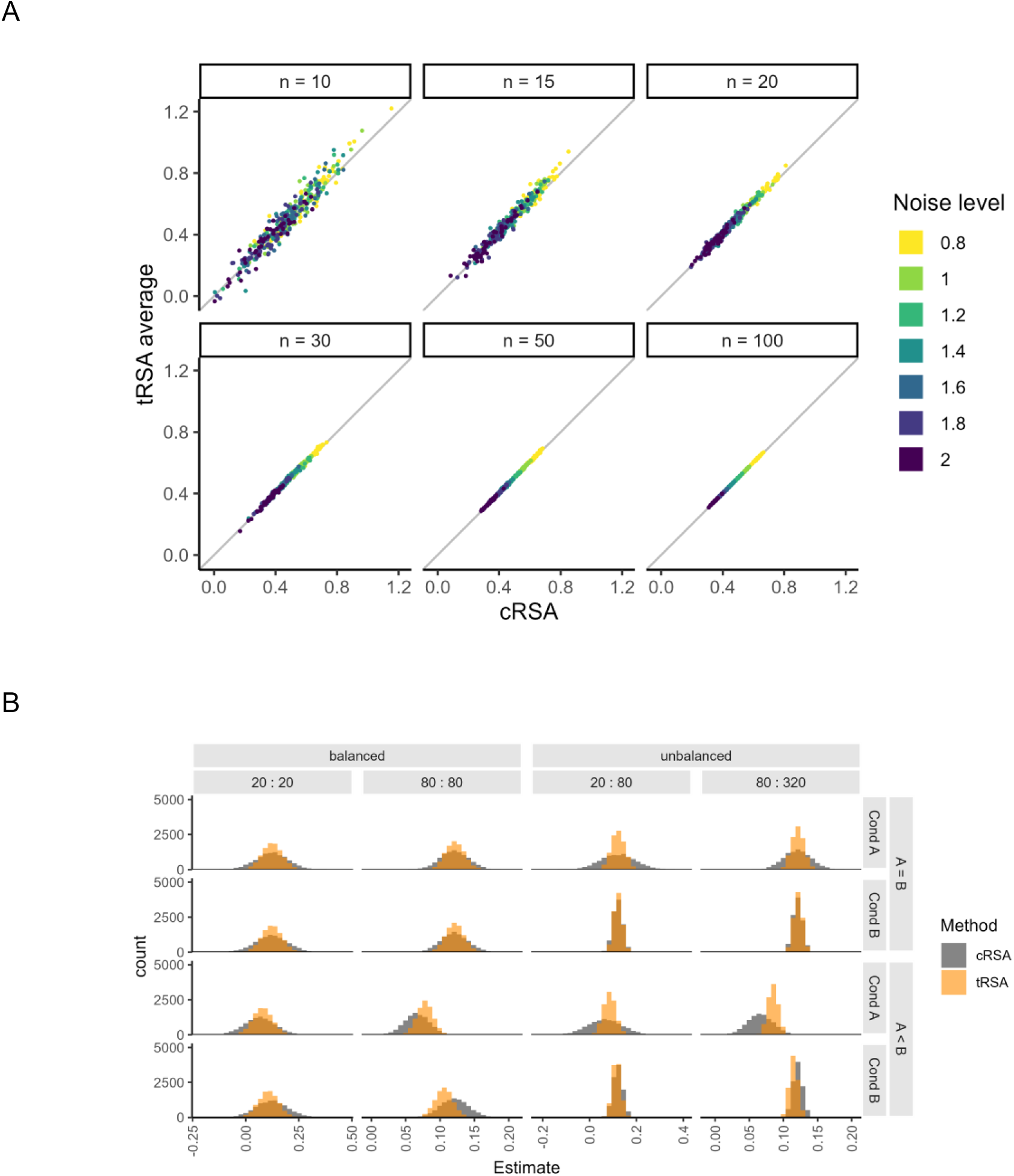
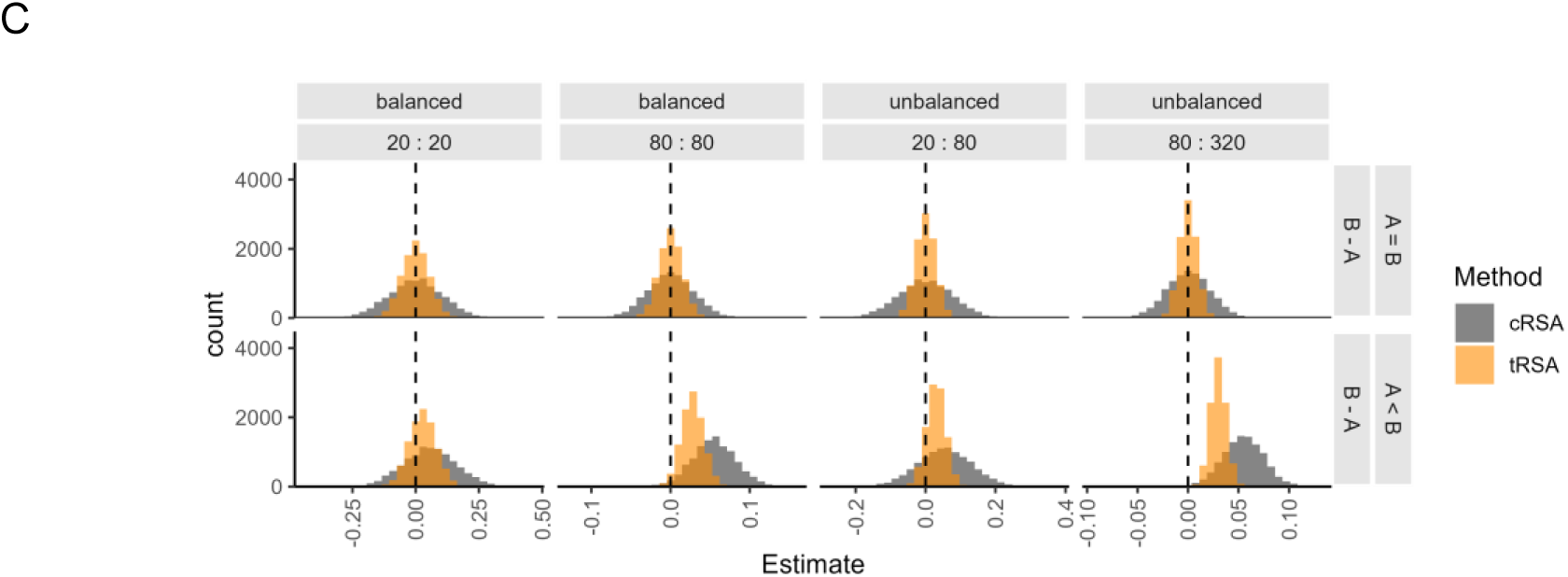
Correspondence between classic and trial-level RSA. **A)** Scatter plots of overall representational strength estimates, based on simulated activity patterns, produced by cRSA (x-axis) and tRSA (y-axis). Trial count (n) and measurement noise level (σ^2^) were both varied. 50 iterations were performed for each parameter combination. Solid slopes indicate where tRSA equaled cRSA (y = x). **B)** Histograms of overall representational strength estimates, based on simulated activity patterns for two separate conditions, produced by cRSA (gray) and across-condition tRSA (orange). Each histogram depicted 10,000 iterations. Trial counts were varied; for example, “20: 80” means that Condition A contained 20 trials and Condition B contained 80 trials, which was an unbalanced scenario. Effect sizes were also varied, such that the ground truth representational strength would be equal between conditions (“A = B”) or stronger in Condition B (“A < B”). **C)** Histograms of condition differences in representational strength values in **B**.

#### Discrete conditions

Oftentimes, researchers are interested in not only representational strength *per se*, but also how representation *changes* with other factors like experimental manipulations (e.g., “easy” vs. “hard”), behavioral performance (e.g., “remembered” vs. “forgotten”), or some combination. Analyzing these differences with cRSA entails partitioning the data to generate separate condition-specific RSMs. Subsequently, an intuitive version of *within-condition* tRSA can be computed using those RSMs, and its estimates would closely track cRSA estimates, as we have demonstrated in the previous section. Instead, we advocate for *across-condition* tRSA: full RSMs are used for computing trial-level representational strength, and the estimates are then split by conditions as necessary. For a direct comparison of *within-*and *across-condition* tRSA, see **Appendix 2**.

Here, we examined the correspondence of *across-condition* tRSA with cRSA in a wide range of scenarios where data can be meaningfully split into two subsets. Specifically, we assessed the influence of three important factors that may vary widely across studies: *raw trial counts*, *balance of trial counts*, and *effect sizes*. Because cRSA was performed separately for each condition, changes in the trial count or effect size of Condition B had no influence on cRSA estimates for Condition A. However, they indeed affected estimates obtained from *across-condition* tRSA, where all experimental trials were used for representational strength estimation. Specifically, all else being equal, increasing the trial count in Condition B from 20 to 80 or from 80 to 320 improved the reliability of tRSA estimates in Condition A (see **Figure 3B, Rows 1 and 3**). Additionally, increasing the noise level for Condition A to be higher than that for Condition B led to reduced tRSA estimates in Condition B as well. These effects were also reflected in the condition difference in tRSA estimates (see **Figure 3C**). Moreover, it is critical to note that tRSA appeared to be more reliable than cRSA in almost all simulated scenarios, especially in terms of the estimated difference between conditions. These findings suggest that tRSA may be more sensitive to true effects and at lower risk of false positives, i.e., reduced Type II and Type I error rates in formal statistical tests.

#### Contrasting conditions

The next set of simulations compared across-condition tRSA to cRSA in terms of statistical inferences, focusing on the difference in representational strength between two conditions as a typical effect of interest in many empirical studies. We implemented the condition differences by altering two condition-level noise parameters, where more noise would result in lower representational strength. The statistical significance of the condition difference *b_B_*_−*A*_ was estimated by a paired-sample t-test for cRSA and by a linear mixed-effects model for tRSA, using a threshold of α = 0.05. Depending on whether the two conditions ought to have the same quality of representation, outcomes were classified as true positives, true negatives, false positives (Type I error), or false negatives (Type II error). The performance of cRSA and tRSA were quantified with their specificity (better avoids false positives, 1 - Type I error rate) and sensitivity (better avoids false negatives 1 - Type II error rate).

We first examined how cRSA and tRSA performed with effects of different sizes, under a base within-subject experimental design consisting of 40 subjects and 200 unique stimuli randomly and evenly split between two conditions. Across 10,000 simulations, cRSA and tRSA were comparable in terms of specificity, with both Type I error rates neighboring around the nominal 5% (see **Figure 4**, diagonal cells in red). As the true condition difference increased, the sensitivity of both methods also increased accordingly. Importantly, tRSA were significantly more sensitive to true effects than cRSA in a range of simulated scenarios, consistent with our hypothesis (see **Figure 4**, off-diagonal cells in blue; asterisks indicate where tRSA was statistically more sensitive then cRSA).

**Figure 4.**
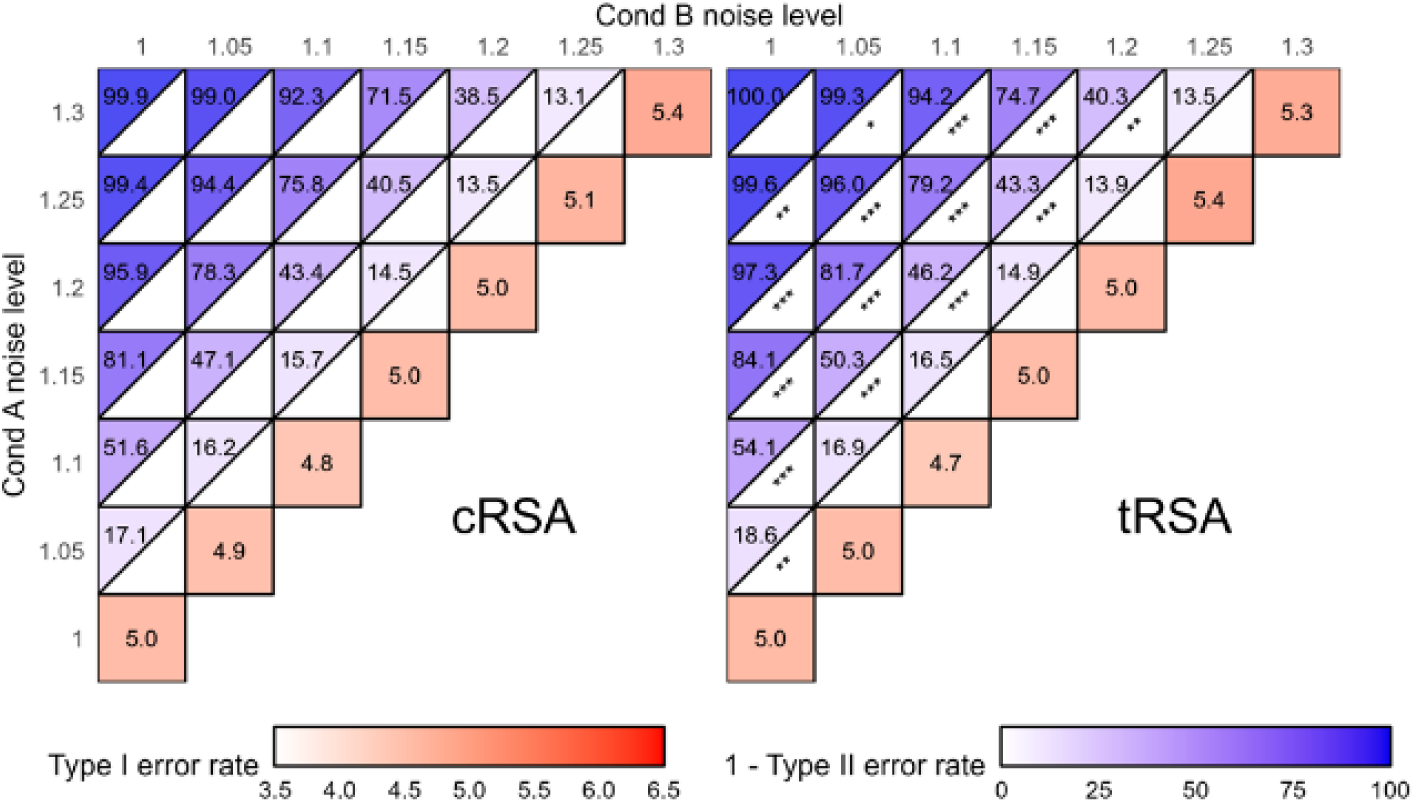
Testing condition differences in representational strength with varying effect sizes. The effect size of the condition difference in representation was manipulated by changing the noise level in each condition in small increments, with higher noise levels corresponding to lower ground truth representational strengths. When the noise level was the same for both conditions, Type I error rates (red) were computed as the proportion of significant contrasts across 10,000 iterations, regardless of sign of the estimate. Otherwise, proportions were computed separately for effects of the correct sign (+, or B>A; blue) and of the incorrect sign (-, or B<A). Asterisks indicate the significance level of the test of equal proportions between simulation results from cRSA (left) and tRSA (right). Significance annotation: *** p < 0.001, ** p < 0.01, * p < 0.05.

We further probed the robustness of cRSA and tRSA across datasets that varied in several important experimental design factors, namely the number of participants, trial counts per participant, and the balance of trial counts. In terms of specificity, our simulations did not suggest an effect of those manipulated factors, as the proportions of significant contrasts yielded by both cRSA and tRSA did not significantly deviate from the nominal alpha level of 5%. Tests of significant contrast proportions reported by cRSA and by tRSA also failed to reject the null hypothesis of equal proportions (*p* > 0.05), suggesting similarly acceptable Type I error rates of both approaches (see **Figure 5, Left**).

**Figure 5.**
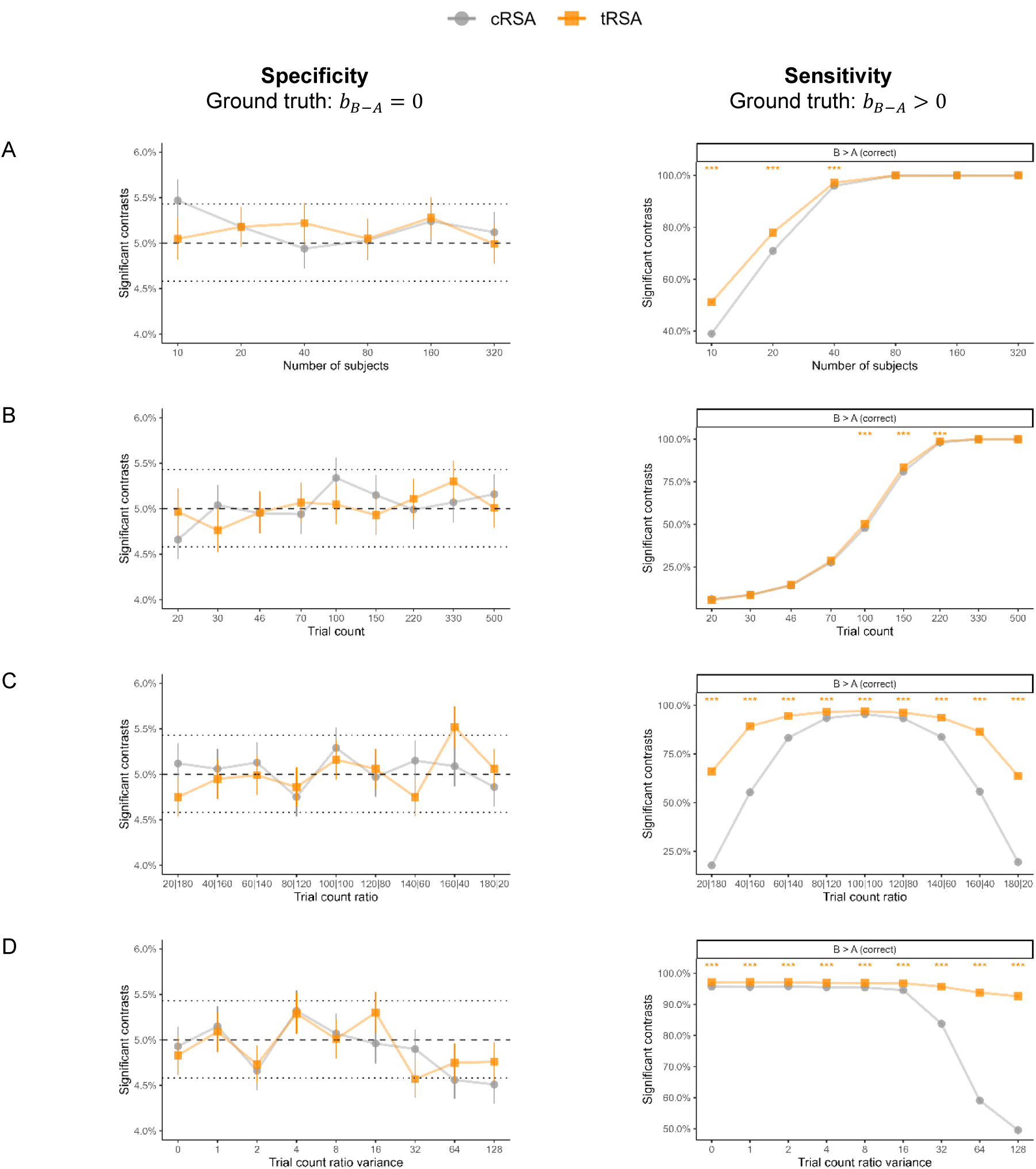
Testing condition differences in representational strength with varying designs. The robustness of cRSA and tRSA were assessed in several variations of a base experimental design consisting of 40 subjects and 200 unique stimuli (randomly and evenly split into two sets). Simulations varied in **A)** number of subjects, **B)** trial count per subject, **C)** the ratio of trial counts between two conditions (with a fixed total), and **D)** the variance of trial count ratios across participants (even split overall). In each simulation, the statistical significance of the condition difference in representational strength was determined by a paired-sample t-test for cRSA and a linear mixed-effects model for tRSA, with α = 0.05. The proportion of significant contrasts was computed across 10,000 iterations. Type I error rates were computed regardless of the direction of the contrast. Type II error rates were computed only for the correct contrast (i.e., B > A). Error bars indicate standard errors. In the left column, dashed horizontal lines mark the nominal α level of 0.05, and dotted horizontal lines mark the critical values beyond which the estimated proportion would be significantly different from α. Asterisks indicate the significance of the deviation in error rates between cRSA and tRSA results, *** p < 0.001, ** p < 0.01, * p < 0.05.

In terms of sensitivity, our simulations revealed large impacts of all manipulated parameters. Increasing the number of subjects or the trial count per subject improved the sensitivity of true effects, benefiting both cRSA and tRSA substantially. Tests comparing the sensitivity of tRSA and cRSA suggested an advantage of tRSA at small-to-medium sample sizes of up to at least 40 subjects, and at medium trial counts of between 100 and 220 trials per subject (see **Figure 5A-B, Right**). We additionally tested variabilities in trial counts that may occur in real experiments. In one set of simulations, the total trial count was fixed at 200 but the ratio of trial counts between conditions varied consistently across subjects (i.e., systematic imbalance; **Figure 5C**). In another set of simulations, while trial counts were kept approximately equal between conditions *on average*, subject-specific ratios varied (**Figure 5D**). In both scenarios, increasing heterogeneity in trial counts resulted in substantially reduced sensitivity for cRSA. This result was expected: the reliability of cRSA estimates depends largely on trial counts (**Figure 2**), which violated the homoscedasticity assumption of the subsequent paired-sample t-test. In the meantime, with trial-level estimates of representational strength from tRSA, the subsequent mixed-effects models were able to properly handle this variance in trial count. Indeed, our simulations validated that tRSA was more robust to extreme scenarios and always significantly outperformed cRSA in sensitivity (see **Figure 5C-D, Right**).

#### Continuously varying effects

A crucial advantage of tRSA over cRSA is that this approach offers a straightforward way to model representational strength as a function of stimulus-or trial-level measures, such as image complexity and subjective familiarity. While these measures could be discretized into a much smaller number of bins (e.g., “low” and “high”), they are better treated as continuous variables to preserve meaningful variance. To simulate these scenarios, we simulated a trial-level measurement variable *v_measured_* that hypothetically tracks trial-level variability *s_trial_* with construct validity *r_val_* = *Cor*(*v_measured_*, *s_trial_*). With higher construct validity, *v_measured_* becomes more predictive of trial-level variability in noisiness and, therefore, representational strength. To perform cRSA, trials were post-hoc split into two conditions by performing a median split on *v_measured_* for each participant. Statistical inferences were again drawn for the difference in representational strength between the two conditions, using both cRSA and tRSA_discrete_. In addition, we also assessed the performance of tRSA_continuous_, where representational strength was directly predicted by the measured continuous variable *v_measured_* without discretization.

Specificity and sensitivity were first examined for datasets with various effect sizes. Across 10,000 iterations, cRSA had a Type I error rate (i.e., when *r_val_* = 0) close to the nominal rate of 5%, while both tRSA_discrete_ and tRSA_continuous_ showed significantly improved specificity (see **Figure 6A**). As construct validity and effect size of the trial-level measurement increased (i.e., more negative *r_val_*), sensitivity expectedly improved for all methods. Relative to cRSA, tRSA_continuous_ appeared to show slightly lower sensitivity for small effects (e.g., *r_val_* = −0.01), though it was also more conservative than cRSA with a lower chance of declaring significance in the incorrect direction (B<A). Moreover, tRSA_continuous_ was the most sensitive method for moderate effects (e.g., *r_val_* = −0.04). We further probed the robustness of cRSA and tRSA across datasets that varied widely in the number of subjects and trial counts. Overall, we observed that the Type I error rates were stable around the nominal 5% for cRSA, as expected. However, both tRSA_discrete_ and tRSA_continuous_ demonstrated significantly enhanced specificity across all manipulations (see **Figure 6B-D, Left**). Sensitivity expectedly improved for all three approaches with increasing subject numbers and trial counts and hit the ceiling with enough samples, though tRSA_continuous_ significantly outperformed other methods at smaller sample sizes, such as 10 participants with 200 trials each or 40 participants with 30 trials each (see **Figure 6B-C, Right**). Additionally, we simulated scenarios in which subject-level trial counts were variable (e.g., missingness, exclusions), resulting in not only a reduction of total sample size but also additional variance in any subject-level estimates. The sensitivity of cRSA dropped substantially with this change because cRSA does not account for such variance, while tRSA-based inferences were minimally impacted (see **Figure 6D, Right**).

**Figure 6.**
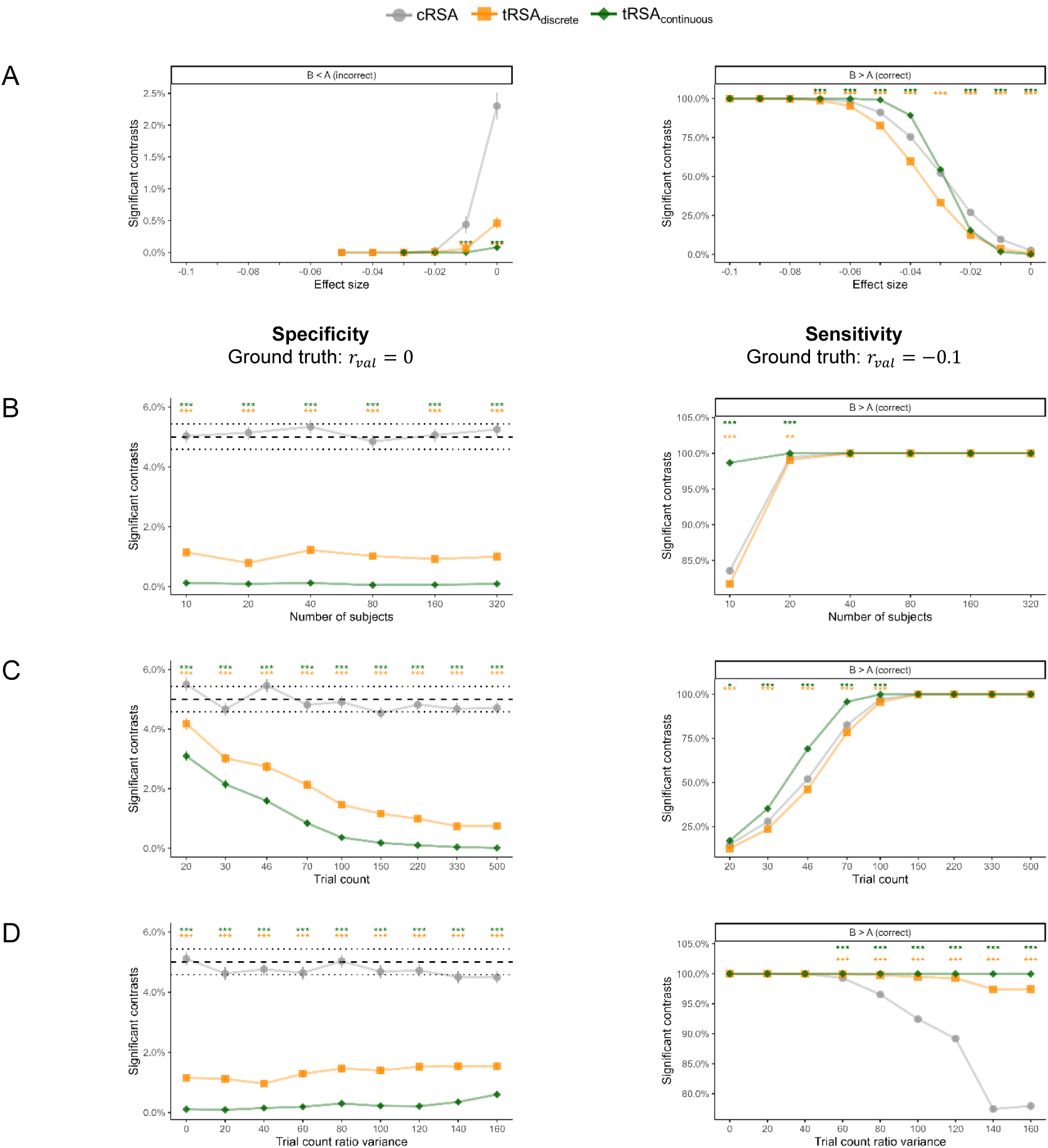
Testing continuous modulations of representational strength with varying designs. The robustness of cRSA and tRSA were assessed in several variations of a base experimental design consisting of 40 subjects and 200 unique stimuli presented once for each subject. Simulations varied in **A)** the true effect size of representational strength, **B)** number of subjects, **C)** trial count per subject, and **D)** the variance of trial count across subjects (assuming a fixed total). Three statistical tests were performed in each simulation: a paired-sample t-test on the difference between “high” and “low” conditions (median-split of v_measured_) for cRSA, a linear mixed-effects model on the same condition difference for tRSA_discrete_, and a linear mixed-effects model on the effect of continuous variable v_measured_ for tRSA_continuous_. Statistical significance was determined with α = 0.05. The proportion of significant results was computed across 10,000 iterations. Type I error rates were computed regardless of the sign of the estimate. Type II error rates were computed only for the correct sign (+). Error bars indicate standard errors. In the left column for **B** through **D**, dashed horizontal lines mark the nominal α level of 0.05, and dotted horizontal lines mark the critical values beyond which the estimated proportion would be significantly different from α. Asterisks indicate the significance level of the test of equal proportions between each tRSA approach and cRSA, *** p < 0.001, ** p < 0.01, * p < 0.05.

#### Summary of Experiment 1

Collectively, our simulations offered three key insights into the commonalities and distinctions between our novel tRSA and the commonly implemented cRSA approaches. First, when the goal is to quantify the overall representational strength of the entire set of trials, tRSA produces estimates highly similar to those generated by cRSA. In other words, little information is lost in performing the trial-level computations (**Figure 3**). Second, tRSA is robust to variances that cRSA is agnostic to, such as imbalances in trial count between conditions and variability in trial count ratios across subjects. In almost all cases tRSA significantly outperformed cRSA in sensitivity (**Figure 5**). This outcome was expected, as the collapsing of trials in cRSA inevitably leads to the loss of important information regarding meaningful variances from other levels; tRSA is capable of properly modelling the multi-level variance structure. Third, tRSA is uniquely advantageous when neural representations are to be linked to continuous variables, rather than discrete conditions. The voided need for post-hoc discretization prevents additional loss of meaningful variance from the modulators, and tRSA_continuous_ demonstrated significantly enhanced specificity and sensitivity over cRSA in a wide range of scenarios (**Figure 6**).

### Experiment 2: cRSA and tRSA comparison in fMRI data

#### Trial-level RSA estimates for Perceptual and Mnemonic Processing

The first goal of Experiment 2 was to compare the results from tRSA and cRSA approaches obtained from real fMRI data. We used fMRI data collected when participants implicitly named 114 colored images of everyday objects in a white background (Object Perception; see **Figure 7A**). Estimates of regional representational strength in the tRSA models were compared with mean cRSA values across participants, for each region of interest. Unsurprisingly, the tRSA model estimates and the cRSA mean correlation values demonstrated similar distributions (**Figure 7B**) and were highly correlated (**Figure 7C**). Critically, however, the two approaches yielded different results in terms of statistically significant regions. Specifically, the cRSA approach identified four significant regions in the LOC during Object Perception, whereas the tRSA approach identified two additional regions within the LOC (**Table 1**, **Figure 7D**, and **Figure 7E**). This discrepancy indicated the ability of tRSA to more accurately and robustly identify representational regions.

**Figure 7.**
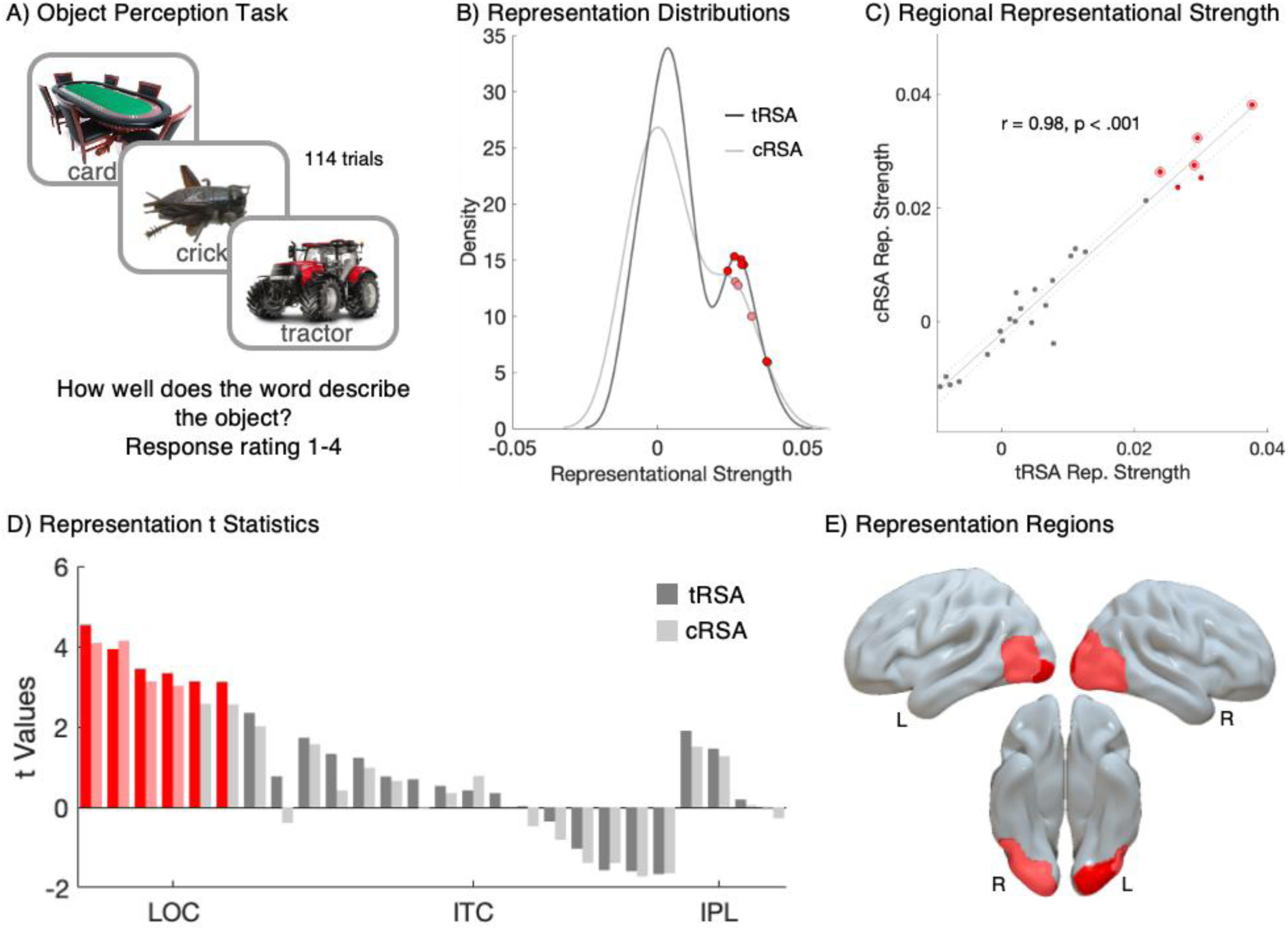
tRSA outperforms cRSA in identifying representational regions during object perception. **A)** Visualization of the Object Perception Task. The fMRI task required participants to view 114 labeled objects and rate how well the labels describe the objects on a scale of 1 to 4 (mean 3.59). **B)** Density plots (y-axis) of regional representational strengths (x-axis) for tRSA (model estimates, dark) and cRSA (mean correlation values, light).Regions with significant representational strength greater than zero (q < 0.05) are highlighted in red. **C)** Scatter plot showing the correlation between regional representational strengths from tRSA (model estimates, y-axis) and cRSA (mean correlation values, x-axis). Regions with significant tRSA representation are highlighted in red, and those significant in cRSA are outlined in red. The representational strength measures are highly correlated across regions (r = 0.98, p < 0.001). **D)** Bar graph of t-statistics for each of the 26 regions of interest. Representation t-statistics (y-axis) from cRSA (light) and tRSA (dark) across 26 regions of interest (x-axis). Regions significant after FDR correction (q < 0.05) are highlighted in red. **E)** Brain regions with significant representation. Regions in light red were identified by the tRSA and cRSA. Regions in dark red were identified by the tRSA method only.

**Table 1.**
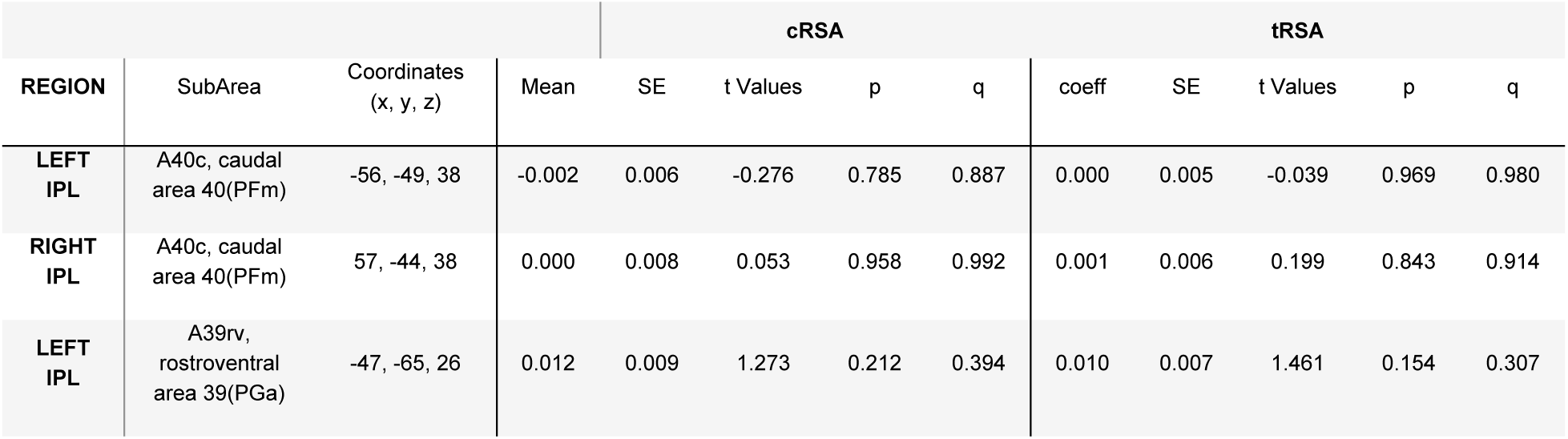
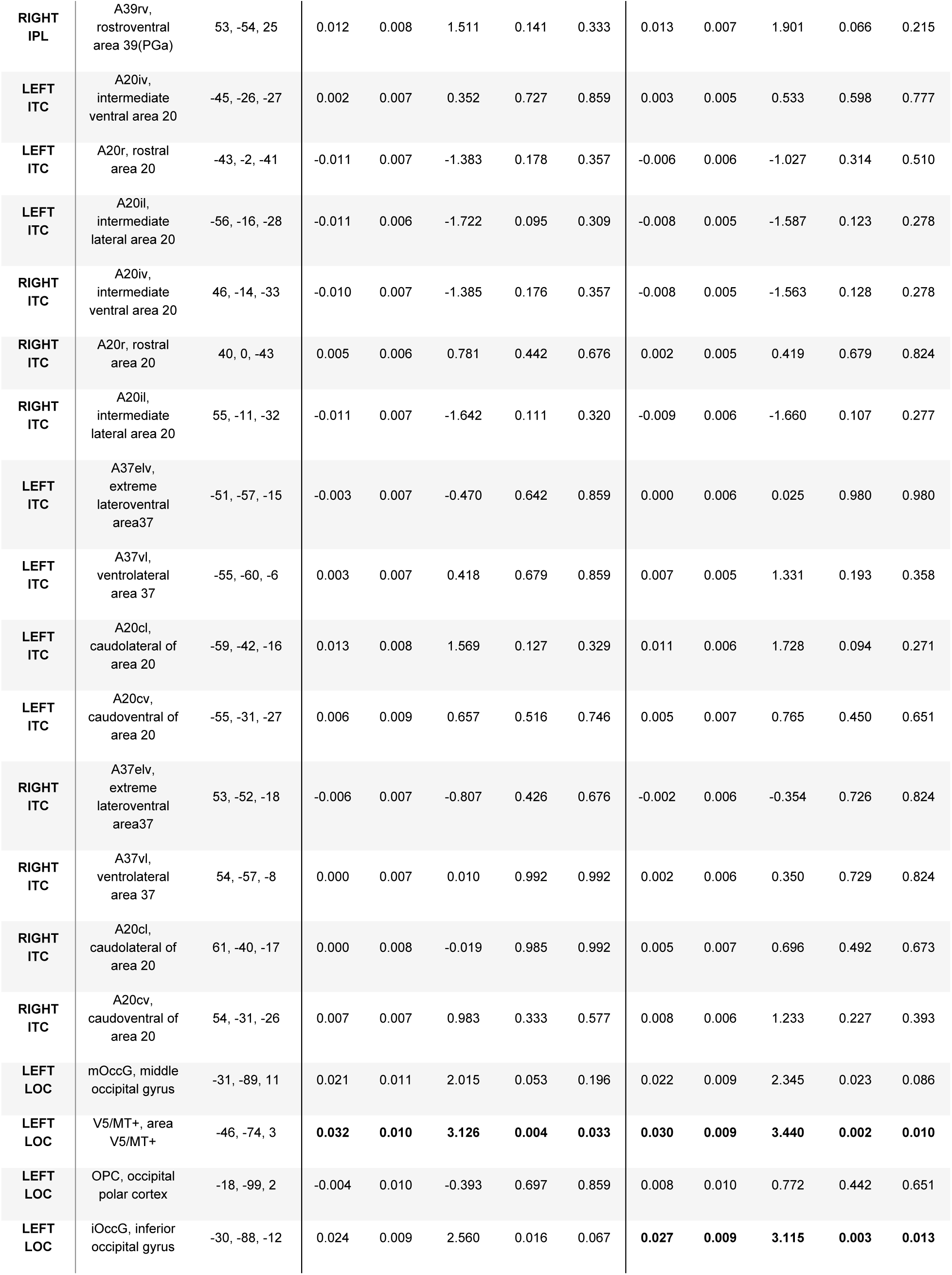
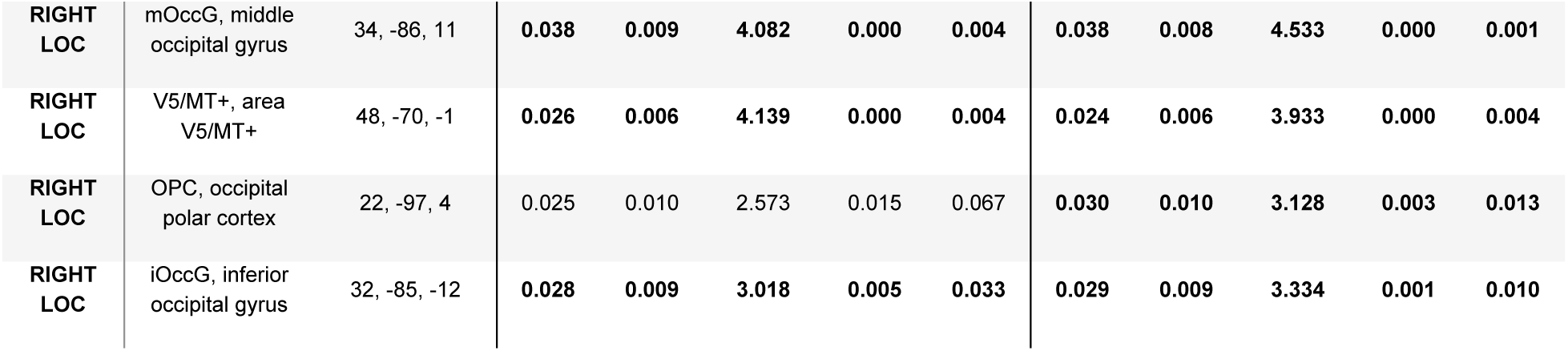
Representation regions identified by cRSA and tRSA.

#### Subject-level variance not accounted for by cRSA

The second goal of Experiment 2 was to demonstrate the utility of tRSA in accounting for the heterogeneity in subject-level variance. In the main Memory Retrieval task, experimental trials were categorized as “Hit” or “Miss” based on participants’ memory success, and the exact number of trials in each condition varied widely across participants (**Figure 8A, Bottom**). Importantly, the difference in the number of trials had a strong impact on the estimation of representational strength (see **Figure 2**). Participants with fewer trials in a given condition exhibited a noticeably wider distribution of representational strength values across ROIs in that condition, indicating that trial count was a substantial source of across-subject variance indeed (see **Figure 8B**). Moreover, trial count was also negatively associated with the standard deviation of representational strengths across ROIs, for both Hit trials (rho = −0.37, *p* = 0.043) and Miss trials (rho = −0.71, *p* < 0.001). However, this subject-level variance is often not accounted for in subsequent group-level t-test or ANOVA, which assumes that each participant contributes equally reliable estimates (i.e., homoscedasticity). Indeed, CVFs were much higher for participants with more extreme Hit to Miss trial counts (rho = 0.45, p = 0.016; see **Figure 8C**).

**Figure 8.**
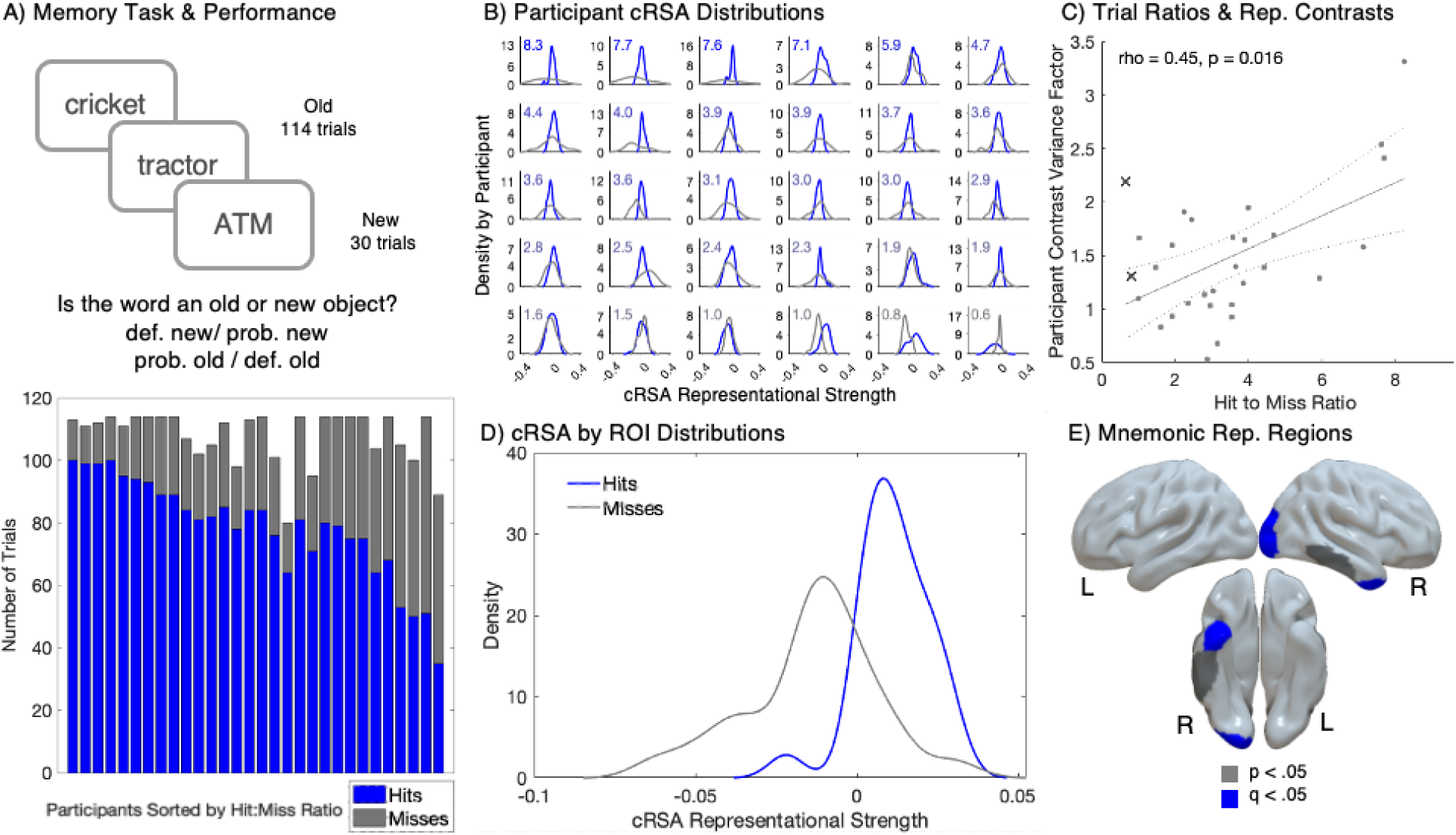
Unmodeled within-and between-participant variance in cRSA for mnemonic representations. **A) Top:** Visualization of the main memory task. Participant recognition and judgement confidence was tested for 144 concepts (114 “old”, 30 “new”). **Bottom:** Stacked bar plots with the number of “Hit” (blue) and “Miss” (gray) trials (y-axes) for each participant (x-axis). Participants were sorted by their hit to miss ratio in descending order. **B)** Density plots of cRSA representational strength (y-axes) across the 26 ROIs (x-axes) for Hits (blue) and Misses (gray) for each participant, sorted by descending Hit-to-Miss count ratio. The Hit-to-Miss ratios were displayed in the top-left corner of each subplot, with the font color transitioning from blue (indicating more Hits) to gray (indicating more Misses). **C)** Scatter plot illustrating the influence of the Hit-to-Miss ratio (x-axis) on the participant CVFs (y-axis). The Spearman’s correlation was significant, with a coefficient of 0.45 and a p-value of 0.016. Two participants are marked X and were excluded from this correlation as their Hit-to Miss ratio is less than 1. **D)** Density plots (y-axis) of cRSA representational strength across participants for Hits (blue) and Misses (gray). **E)** Regions exhibiting significant cRSA mnemonic representation (Hits > Misses). Regions highlighted in blue were significant after FDR correction with q < 0.05. Regions in gray were significant at p < 0.05 but did not survive FDR correction.

This issue of neglected heteroscedasticity remains when conditional contrasts (e.g., Hit minus Miss) are computed. Given the nonlinear relationship between trial counts and the reliability of correlations (see **Figure 2**), a more extreme trial imbalance (e.g., 100 Hits and 10 Misses, versus 70 Hits and 40 Misses) would also lead to a less reliable estimate of the difference. To demonstrate this issue in real fMRI data, we computed the CVF for each participant, measuring how much their Hit-Miss contrast in representational strength estimates deviated from the group mean. Indeed, CVFs were much higher for participants with more extreme Hit to Miss trial counts (rho = 0.45, p = 0.016; see **Figure 8C**). Critically, the two participants with Hit to Miss ratios < 1 were excluded from this correlation (since in those cases we would expect an increased CVF), but their data points are included in the corresponding figure for transparency.

Additionally, the distributions of representational strengths for Hits and Misses across participants were unequal. Since most participants had more Hit trials than Miss trials (Hit-to-Miss Ratio mean = 3.7, median = 3.1), the distribution of representational strength values was broader for misses compared to hits (**Figure 8D**). Ultimately, results from cRSA revealed significant mnemonic representation in one right LOC region and one right anterior ITC region, with two additional ITC regions showing trend-level significance (**Table 2**, **Figure 8E**); these outcomes were revisited in comparison to the tRSA approach below. Together, these results demonstrate that cRSA failed to model the across-subject variance in real fMRI data, which stemmed from the inevitable variability in trial count ratios affecting the reliability of each participant’s cRSA contrast estimates. The violation of statistical assumptions challenged the interpretability of standard t-tests in detecting representational regions.

**Table 2.**
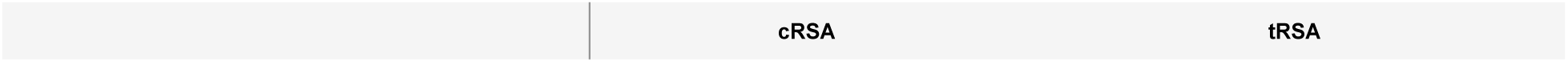
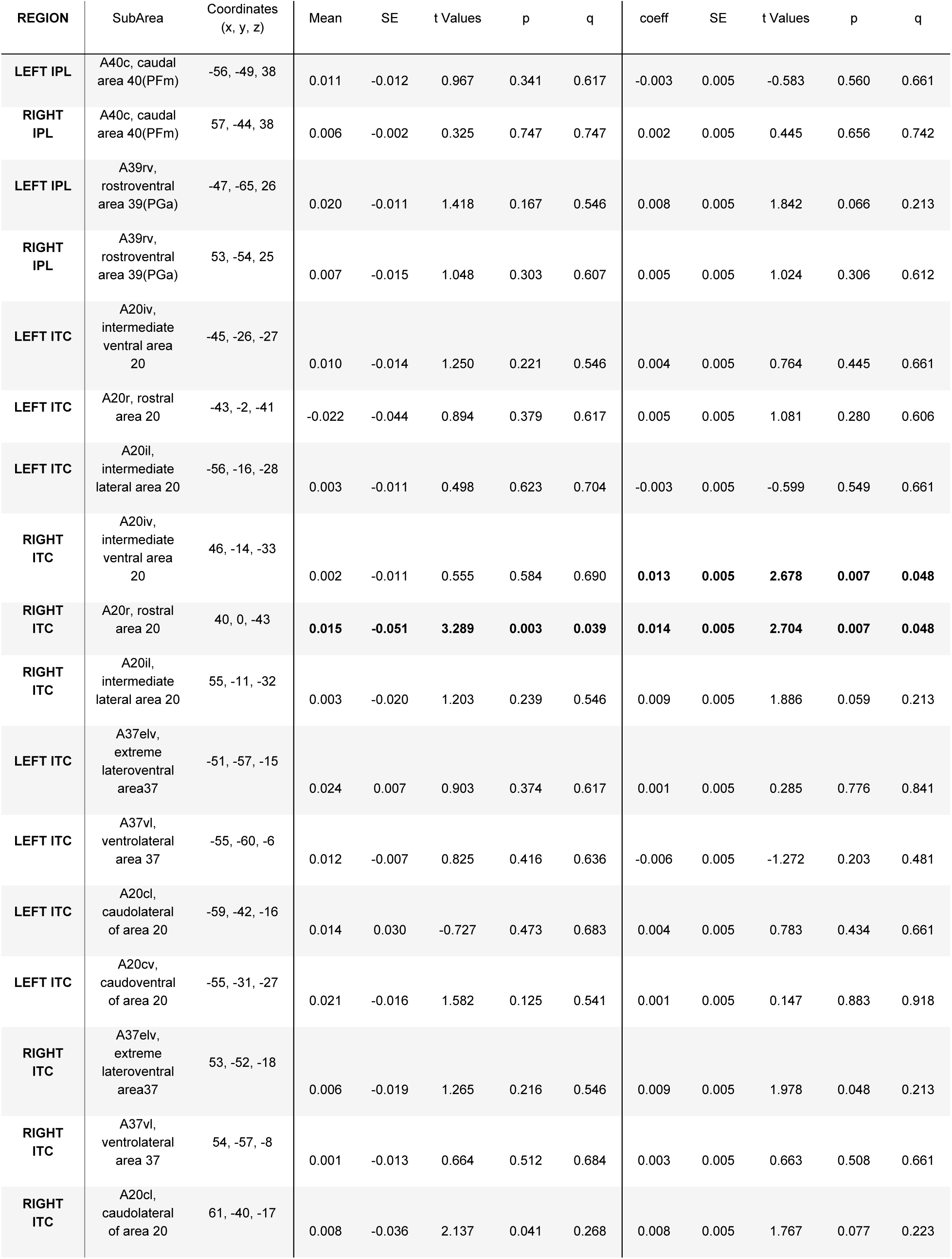
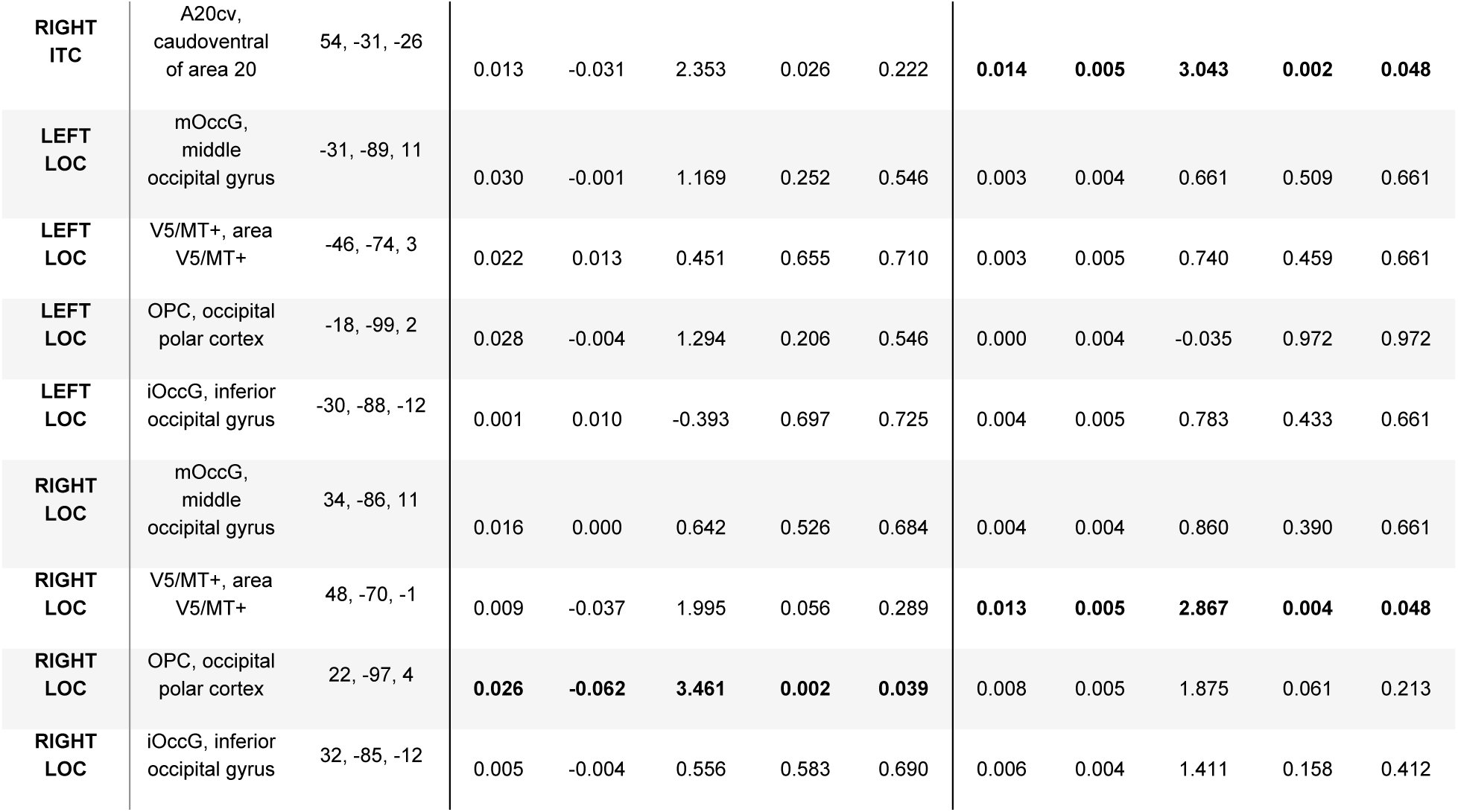
Mnemonic representation regions identified by cRSA and tRSA.

#### Subject-level variance accounted for by tRSA

We also sought to assess how the tRSA approach would improve the estimates of representational strength for memory success. The mixed-effects models used to test tRSA mnemonic representations supporting retrieval identified four significant mnemonic representation regions: one in the right LOC and three in the right ITC (**Table 2**, **Figure 9A**, and **Figure 9B**). While the t-statistics for tRSA and cRSA showed a correlation across ROIs (**Figure 9C**), the tRSA approach, which produced trial-level estimates of representational strength, was able to detect mnemonic representation regions that the cRSA approach did not. This result demonstrated the enhanced sensitivity of tRSA in identifying subtle differences in representational strength. Moreover, a region in the LOC, initially considered a mnemonic representation region by the cRSA method — largely due to data from participants with low miss trial counts (**Appendix 3 Table 4**) — was correctly excluded from being classified as such by tRSA (**Figure 9A**, **Figure 9B**). This result highlighted tRSA’s improved specificity in the identification of true mnemonic representation regions by reducing potential biases introduced by variances in trial counts that went neglected by cRSA.

**Figure 9.**
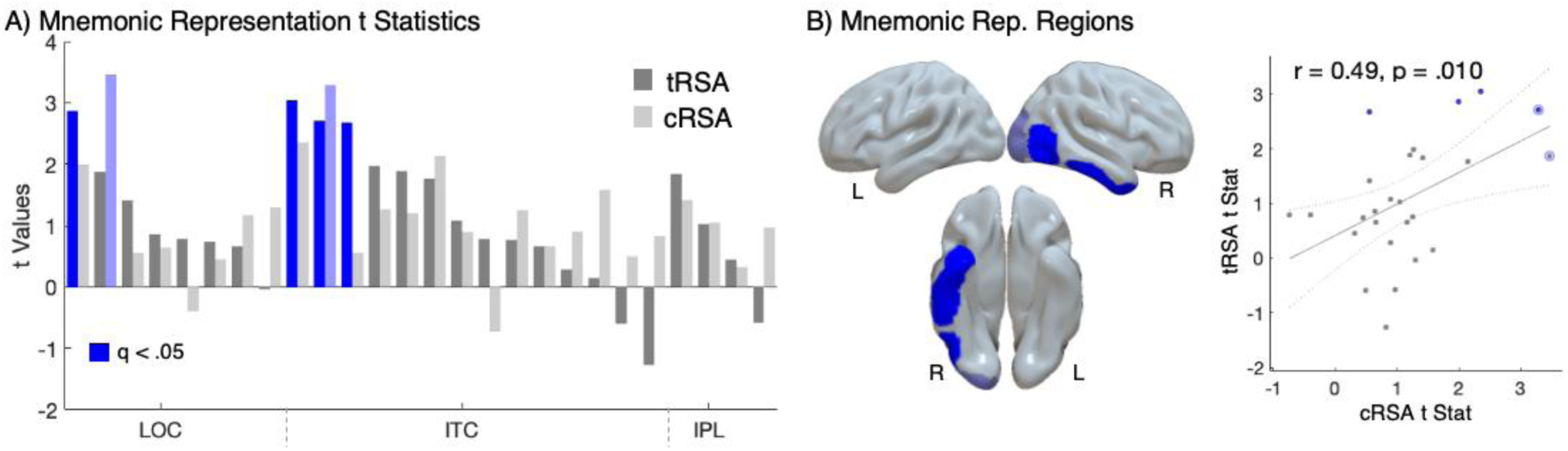
tRSA outperforms cRSA in detecting mnemonic representations. **A)** Bar graph of t Statistics for each of the 26 regions of interest. Mnemonic representation t-statistics (y-axis) from cRSA (light) and tRSA (dark) across 26 regions of interest (x-axis). Regions that are significant after FDR correction (q < 0.05) are highlighted in blue. **B)** Brain regions with significant mnemonic representation identified by tRSA are shown in blue. The region that was significant in cRSA but not in tRSA is depicted in light blue. **C)** Scatter plot of t-statistics calculated from tRSA (y-axis) and cRSA (x-axis), showing a significant Pearson correlation (r = 0.49, p = 0.010). Significant tRSA regions are highlighted in blue. Significant cRSA regions are outlined in light blue.

#### Estimations of continuous modulators

Our final goal was to assess the utility of tRSA in testing hypotheses regarding continuous modulators of representational strength. Here, we focused on the effect of item memorability, a stimulus-level variable (see **Figure 10A**). The tRSA approach successfully detected continuous, stimulus-level modulation of representational strength during Object Perception, when participants were engaged in basic-level naming of color objects displayed on a white background; notably, no memory task was engaged before or during this phase of the experiment. Four regions within the LOC showed significant modulation by Perceptual Memorability, a measure based on average responses in the Perceptual Retrieval task (see **Table 3 and Figure 10B**). While not the explicit focus of this analysis, the analyses also identified significant effects of the nuisance variables, Conceptual Memorability, and reaction time in five regions of interest (**Appendix 3, Table 6**) within the LOC. These regions are often associated with processing the constituent categories or visual features of an object (Tyler et al., 2013), and consistent with the view that the memorability of images can be predicted by statistical properties of semantic features (Bainbridge, 2022; Hovhannisyan et al., 2021; Kramer et al., 2023). Critically, memorability is a robust stimulus property that is consistent across participants and paradigms(Bainbridge, 2022). Moreover, object memorability effects have been replicated using a variety of methods aside from tRSA, including univariate analyses and representational analyses of neural activity patterns where trial-level neural activity pattern estimates are correlated directly with object memorability (Slayton et al., 2025).

**Figure 10.**
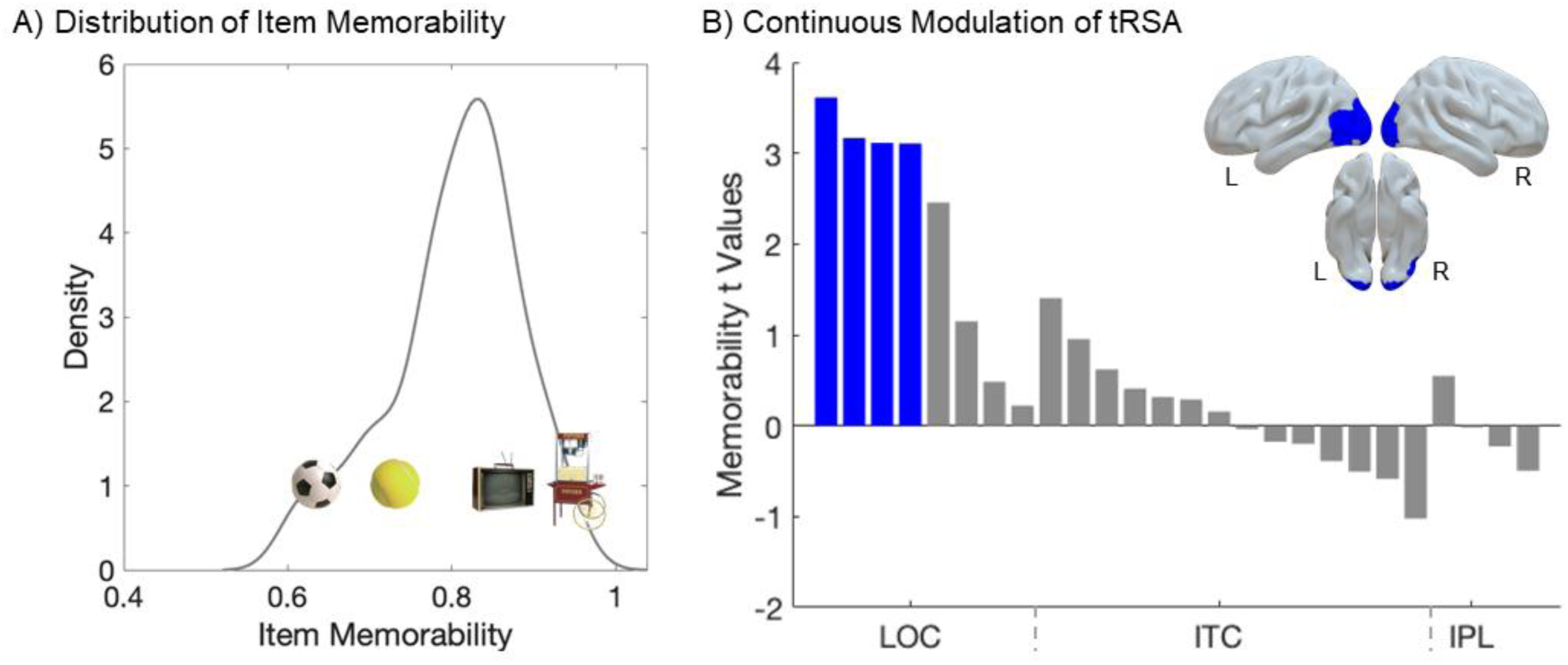
tRSA captures stimulus-level modulations on representation. **A)** Distribution of Item Memorability. Density plot of memorability values demonstrates a wide distribution of memorability scores (i.e., the average confidence of remembering a particular item). This stimulus-level variance is an underexplored target of representational analyses. **B)** Cortical representational effects of Memorability. Relationship between item-level Memorability and representational strength, as assessed with tRSA, for each of the 26 regions of interest. Trial-level representational strength estimates from Object Perception were significantly modulated by a continuous, stimulus-level measure of Item Memorability in four regions (blue). The t-statistics for this modulation were plotted (y-axis) across regions of interest (x-axis).

**Table 3.** Continuous modulation in tRSA.

| REGION | SubArea | Coordinates<br>(x, y, z) | coeff | SE | t Values | p | q |
| --- | --- | --- | --- | --- | --- | --- | --- |
| LEFT IPL | A40c, caudal<br>area 40(PFm) | -56, -49, 38 | -0.036 | 0.072 | -0.496 | 0.620 | 0.949 |
| RIGHT IPL | A40c, caudal<br>area 40(PFm) | 57, -44, 38 | 0.038 | 0.069 | 0.547 | 0.584 | 0.949 |
| LEFT IPL | A39rv,<br>rostroventral<br>area 39(PGa) | -47, -65, 26 | -0.015 | 0.068 | -0.227 | 0.820 | 0.949 |
| RIGHT IPL | A39rv,<br>rostroventral<br>area 39(PGa) | 53, -54, 25 | -0.001 | 0.071 | -0.018 | 0.986 | 0.986 |
| LEFT ITC | A20iv,<br>intermediate<br>ventral area 20 | -45, -26, -27 | 0.044 | 0.071 | 0.621 | 0.534 | 0.949 |
| LEFT ITC | A20r, rostral<br>area 20 | -43, -2, -41 | 0.025 | 0.079 | 0.315 | 0.752 | 0.949 |
| LEFT ITC | A20il,<br>intermediate<br>lateral area 20 | -56, -16, -28 | 0.068 | 0.072 | 0.955 | 0.340 | 0.949 |
| RIGHT ITC | A20iv,<br>intermediate<br>ventral area 20 | 46, -14, -33 | -0.013 | 0.074 | -0.177 | 0.860 | 0.949 |
| RIGHT ITC | A20r, rostral<br>area 20 | 40, 0, -43 | -0.046 | 0.078 | -0.587 | 0.558 | 0.949 |
| RIGHT ITC | A20il,<br>intermediate<br>lateral area 20 | 55, -11, -32 | 0.021 | 0.071 | 0.290 | 0.771 | 0.949 |
| LEFT ITC | A37elv,<br>extreme<br>lateroventral<br>area37 | -51, -57, -15 | 0.029 | 0.071 | 0.409 | 0.682 | 0.949 |
| LEFT ITC | A37vl,<br>ventrolateral<br>area 37 | -55, -60, -6 | 0.105 | 0.075 | 1.405 | 0.160 | 0.694 |
| LEFT ITC | A20cl,<br>caudolateral of<br>area 20 | -59, -42, -16 | -0.014 | 0.068 | -0.201 | 0.841 | 0.949 |
| LEFT ITC | A20cv,<br>caudoventral of<br>area 20 | -55, -31, -27 | -0.027 | 0.069 | -0.389 | 0.698 | 0.949 |
| RIGHT ITC | A37elv,<br>extreme<br>lateroventral<br>area37 | 53, -52, -18 | -0.073 | 0.071 | -1.022 | 0.307 | 0.949 |
| RIGHT ITC | A37vl,<br>ventrolateral<br>area 37 | 54, -57, -8 | 0.011 | 0.070 | 0.156 | 0.876 | 0.949 |
| RIGHT ITC | A20cl,<br>caudolateral of<br>area 20 | 61, -40, -17 | -0.003 | 0.072 | -0.038 | 0.970 | 0.986 |
| RIGHT ITC | A20cv,<br>caudoventral of<br>area 20 | 54, -31, -26 | -0.036 | 0.072 | -0.506 | 0.613 | 0.949 |
| LEFT LOC | mOccG, middle<br>occipital gyrus | -31, -89, 11 | **0.255** | **0.080** | **3.166** | **0.002** | **0.012** |
| LEFT LOC | V5/MT+, area<br>V5/MT+ | -46, -74, 3 | **0.258** | **0.071** | **3.614** | **0.000** | **0.008** |
| LEFT LOC | OPC, occipital<br>polar cortex | -18, -99, 2 | **0.276** | **0.089** | **3.113** | **0.002** | **0.012** |
| LEFT LOC | iOccG, inferior<br>occipital gyrus | -30, -88, -12 | 0.019 | 0.083 | 0.223 | 0.823 | 0.949 |
| RIGHT LOC | mOccG, middle<br>occipital gyrus | 34, -86, 11 | 0.201 | 0.082 | 2.456 | 0.014 | 0.073 |
| RIGHT LOC | V5/MT+, area<br>V5/MT+ | 48, -70, -1 | 0.089 | 0.077 | 1.149 | 0.251 | 0.931 |
| RIGHT LOC | OPC, occipital<br>polar cortex | 22, -97, 4 | **0.277** | **0.089** | **3.107** | **0.002** | **0.012** |
| RIGHT LOC | iOccG, inferior<br>occipital gyrus | 32, -85, -12 | 0.040 | 0.083 | 0.485 | 0.628 | 0.949 |

#### Summary of Experiment 2

Three key takeaways from Experiment 2 mirrored those from Experiment 1. First, while access to ground truths was not possible with empirical fMRI datasets, results from tRSA and cRSA approaches showed a relatively high degree of correspondence as before (**Figure 7C**). Nonetheless, modeling trial-level estimates from tRSA showed improved sensitivity to neural representations in the Object Perception dataset, as the tRSA model was able to weigh study subjects with different numbers of trials appropriately (**Figure 7D**). Second, in the Memory Retrieval dataset, participants had unbalanced and heterogenous distributions of trial counts in two conditions (“Hits” and “Misses”), as expected with any psychological task focusing on subject performance (**Figure 8A**). tRSA’s improved characterization can be seen in multiple empirical outcomes: tRSA was able to detect mnemonic representation regions that the cRSA approach missed (improved; sensitivity **Figure 9B**), and afforded more appropriate weighting of participant’s contrast estimates while maintaining similar outcome statistics as cRSA (**Figure 9C**). Third, tRSA supported the investigation of research questions that cannot be readily addressed with cRSA, namely the trial-or stimulus-level modulations of representational strength. We demonstrated this point using Item Memorability — a stable stimulus property that is thought to contribute to memory independently of task context, experimental context, or individual differences. Representational strength in several regions in the ventral stream varied continuously with stimulus-level memorability (**Figure 10B**). Moreover, model-fits became the strongest when the models also incorporated trial-level variables such as fMRI run and reaction time (**Appendix 3 Table 6**). These findings showed that the representational strength estimates produced by tRSA indeed captured the multi-level variance structure in the data and that tRSA can be implemented to study item-specific or trial-level modulators of representational strength effectively.

## Discussion

This paper formally presents tRSA — a novel technique for evaluating the strength of neural representation at the level of individual experimental trials. The performance of tRSA was evaluated and compared to that of cRSA using both simulations (Experiment 1) and empirical fMRI datasets (Experiment 2). Three principal insights can be drawn from these analyses. First, tRSA produces highly similar estimates of *overall* representational strength as cRSA such this new approach comes with little cost in efficacy. Second, tRSA is robust to subject-level variances that would be neglected by cRSA, namely the trial count differences across subjects. With both simulations and empirical datasets, tRSA demonstrated enhanced sensitivity over cRSA in detecting true effects. Third, tRSA provides an opportunity to examine continuous modulators of representational strength with decent specificity and sensitivity. With the empirical fMRI dataset, we demonstrated that tRSA was able to detect significant associations between representational strength and item-level memorability, as well as trial-level nuisance variables. Below we discuss the methodological and conceptual implications of the tRSA approach for the field of cognitive neuroscience and beyond.

### Similarities between cRSA and tRSA

Since the advent of cRSA, the method has been widely implemented in studies of neural representations with great success (Dimsdale-Zucker & Ranganath, 2018; Kriegeskorte et al., 2008). Our first and foremost objective was to ensure that our novel tRSA technique produces estimates of representational strength that are comparable to cRSA. This objective was achieved in both Experiment 1 (simulations) and Experiment 2 (real fMRI data). Simulations suggested that tRSA and cRSA estimates reacted in very similar ways to manipulations of the number of subjects, trial counts, and noise level. Although tRSA and cRSA both aim to quantify representational strength, they differ in how they operationalize this concept. cRSA summarizes the correspondence between RSMs as a single measure, such as the matrix-wise Spearman correlation. In contrast, tRSA computes such correspondence for each trial, enabling estimates at the level of individual observations. This flexibility allows trial-level variability to be modeled directly, but also introduces subtle differences in what is being measured. Nonetheless, our simulations showed that, although numerical differences occasionally emerged—particularly when comparing between-condition tRSA estimates to within-condition cRSA estimates—the magnitude of divergence was small and did not affect the outcome of downstream statistical tests. Analyses with real fMRI datasets corroborated their correspondence, showing highly similar distributions of estimated representational strength across the brain. These results confirm that our novel tRSA approach performs at least no worse than cRSA. In other words, researchers can confidently replace their existing cRSA analyses with the tRSA framework, not having to be concerned about the loss of information or inferior statistical performance. To further evaluate demonstrate their correspondence between trial-level RSA (tRSA) relative and classical RSA (cRSA), we conducted an additional representational similarity analysis using a simple categorical semantic model applied to an existing object-naming fMRI dataset originally reported by Clarke & Tyler, 2014 and further analyzed in Devereux et al, 2018 and Seth et al., 2024. The original study examined representational gradients along the ventral visual stream using a combined deep visual-semantic neural network model across early visual cortex, posterior ventral temporal cortex (pVTC), and perirhinal cortex (PRC). Methods and Results are reported in the Appendix.

### Methodological significance

Statistically, tRSA is designed to properly capture the multi-level variance structure in the data, yielding improved specificity and sensitivity. An important motivation of ours for devising this novel tRSA technique accords with a steady emergence in many scientific fields that advocates for proper treatments of the multi-level variance structure in the data (Dedrick et al., 2009; Winter & Grice, 2021; Yarkoni, 2022). As laid out in the Introduction, we argue that (at least) four independent sources of variance contribute to how strongly a given neural system represents a single event (e.g., looking at an image of “basketball”): condition-level, subject-level, stimulus-level, and trial-level.

*Subject-level variance* was a major focus of analysis in both Experiments. Research subjects are samples (ideally, random and representative) drawn from some population of interest. Barring quantifiable individual differences that may be relevant to the cognitive functions being studied (e.g., age, education, personality traits), subjects are considered to vary randomly around the population-level mean, which is often the focus of analysis. In cRSA, subject-level estimates of representational strength or condition differences are typically entered into a general linear model where homoscedasticity is assumed (e.g., t-test). However, this assumption is easily violated due to heterogeneity in subject-level trial counts, resulting in unreliable model outputs. Indeed, artificially induced variability in trial counts led to catastrophic reductions in the sensitivity of cRSA (Experiment 1). Empirical studies that select trials based on subjective performance or expect to have extensive missingness or exclusions must be wary of this problem, as we demonstrated the same issue of cRSA using real fMRI data from an episodic memory study (Experiment 2). Critically, tRSA proved to be more robust to this subject-level heteroscedasticity issue in both Experiments, for trial count information was available to the subsequent models with subjects entered as random factors (Yarkoni, 2022).

*Stimulus-level variance* was another focus of our investigations. A major advantage of RSA over multivariate pattern classification techniques is that it allows the use of “condition-rich” experimental designs with large and diverse collections of stimuli, so long as meaningful hypotheses can be constructed in the form of an RSM_model_ (Kriegeskorte et al., 2008). For instance, using an RSM_model_ based on human similarity judgments, past work has implemented RSA on neuroimaging data collected with nearly two thousand unique natural object images (THINGS database; Hebart et al., 2023). Various stimulus properties have also been the target of neural representation research, such as the real-world size of objects (T. Huang et al., 2022), the geographical location of landmarks (Morton et al., 2021), or more abstractly, visual and semantic features of naturalistic images (Devereux et al., 2018; Jozwik et al., 2023; Naspi et al., 2021). Despite such prevalent appreciation for the neurocognitive relevance of stimulus properties, cRSA often does not account for the fact that the same stimulus (e.g., “basketball”) is seen by multiple subjects and produces statistically dependent data an issue addressed by Schütt et al. (2023), who developed cross validation and bootstrap methods that explicitly model dependence across both subjects and stimulus conditions. With tRSA, this issue is at least partially addressed by modeling stimulus identity as random effects, resulting in generalizability and reliability (Chen et al., 2021; Yarkoni, 2022). Admittedly, a complete resolution of this stimulus-as-fixed-effect fallacy would require stimulus information to be incorporated in any preceding procedures, including first-level models of fMRI data (Westfall et al., 2017); while beyond the scope of the current paper, these relevant analytical pipelines may be integrated for a full solution of the fallacy. Furthermore, as demonstrated in Experiment 2, tRSA supports the straightforward investigation of how representational strength varies with stimulus-level properties such as memorability. With the flexibility of tRSA, future studies may freely explore other interesting stimulus-level research questions.

Of note, in the current paper, we only examined experimental designs where each unique stimulus is presented once, i.e., no repetitions. Repetitions are sometimes included as targets to which participants should respond, in order to promote engagement level (Allen et al., 2022; Chang et al., 2019; Lahner et al., 2024). Additionally, some preprocessing or modeling procedures use repetitions to improve the accuracy of their estimates of neural activity responses and representations (Charest et al., 2018; Prince et al., 2022), though the effectiveness of this approach may depend on various other factors (Ritchie et al., 2021). Importantly, repetitions have been found to robustly induce neural activity adaptation or *repetition suppression* in task-relevant brain regions (Barron et al., 2016; Grill-Spector et al., 2006), and this phenomenon has profound impacts on multi-voxel activity patterns as well (Mazurchuk et al., 2023). While beyond the scope of the current paper, we briefly mention two possible treatments of stimulus repetitions in implementing tRSA. For one, repetitions can be collapsed in first-level models of brain recordings for more reliable estimations of stimulus-specific neural activations; in this case, any subsequent analyses including tRSA can simply proceed as if individual stimuli are only presented once. Alternatively, repetitions can also be kept as separate events for which individual tRSA estimates are assigned; this option allows one to probe the stability of neural representations across time and repetitions. In either case, our tRSA approach can flexibly suit the various goals of research.

*Trial-level variance* refers to the effect on the estimations of neural representations by factors that vary from trial to trial. For example, attention level fluctuates during the task and changes the fidelity of neural representations (Aly & Turk-Browne, 2016; Rothlein et al., 2018). Like stimulus-level variance, trial-level variance is not accounted for by cRSA as the comparison of RSMs collapses across trials. In our simulations, we demonstrated that tRSA can indeed capture trial-level variance and reliably estimate its effect on representational strength (Experiment 1). In a real fMRI dataset, we found that reaction time, a trial-level nuisance variable, also had statistically significant effects on representational strength (Experiment 2). In addition, past research has found that pure physiological changes in pupillary size, cardiac rhythm, and respiration may also influence both neural and cognitive processes (Critchley & Garfinkel, 2018; van der Wel & van Steenbergen, 2018). While a portion of those effects could be due to the incomplete removal of artifacts during signal preprocessing, some physiological changes may also have intriguing neurocognitive relevance that is worth exploring in future studies.

### Conceptual significance

Fundamentally, approaching RSA at the level of individual trials is an advancement in terms of not only *methodology* (discussed above) but also *conceptualization*. The notion of *neural representation*, albeit used more broadly than the phrase is intended to, emphasizes the link between the representee (contents being represented, such as stimuli or memories) and its representative (alternative format of the content, such as neural activity) (Baker et al., 2022; Favela & Machery, 2023; Vilarroya, 2017). Collapsing trial-level information in its formulation, cRSA could only indicate the quality of a *kind* of representation or the *overall* representational strength across events. Instead, tRSA provides a measure that directly corresponds to the conceptualization of representation, with each trial or event receiving its own estimated representational strength. This level of analysis is consistent with previous studies that focused on quantifying the similarity of multi-voxel activity patterns across different trials (S. Huang, Faul, et al., 2024; Wing et al., 2020; Yu et al., 2024) and at different stages of memory (Ritchey et al., 2013; Shao et al., 2023). As such, tRSA provides a versatile tool for inquiring about many different brain-behavior relationships at an appropriate level of analysis and with straightforward interpretations (Becker et al., 2024; Howard et al., 2024; Morales-Torres et al., 2024; Naspi et al., 2023; Pacheco-Estefan et al., 2024). Notably, tRSA is not limited to neuroimaging data; past research has also implemented this approach to perform item-specific analyses on behavioral data (Walsh & Rissman, 2023).

At a broader level, the tRSA approach also engenders a host of other innovative techniques. While a number of studies have addressed the validity of measuring representational geometry using designs with multiple repetitions, a conceptual benefit of the tRSA approach is the reliance on a regression framework that allows competing conceptual models of stimulus representation to be evaluated while simultaneously modeling trial-and stimulus-level predictors within a hierarchical framework (e.g., taxonomic vs. encyclopedic semantic features, as in Davis et al., 2021). Unlike approaches that rely on repeated stimulus presentations to improve estimates of the underlying neural response, tRSA does not seek to recover a noise-free representation for each individual trial. Single-trial activation estimates inevitably contain measurement noise, and this limitation is shared by any analysis based on single-trial fMRI estimates. Instead, tRSA treats these estimates as noisy observations of representational structure and leverages mixed-effects modeling to estimate the relationship between neural and model representational similarity across participants and stimuli. Consequently, the method does not eliminate measurement noise but rather accommodates it within the statistical model, allowing population-level inferences without requiring repeated stimulus presentations. The emerging research on representational connectivity focuses on the intersection of representational information and functional connectivity analyses, and seeks to uncover new multivariate views on how concepts are distributed across the brain (Anzellotti & Coutanche, 2018). Using the tRSA approach, trial-level representational strength values from any pair of brain regions may be correlated to yield their model-based representational connectivity (S. Huang, De Brigard, et al., 2024; Karimi-Rouzbahani et al., 2022). Trial-level representational strength values may also be combined with other trial-level measures of neural activity such as univariate activation to probe diverse inter-regional interactions that are theoretically motivated (S. Huang, Howard, et al., 2024).

### Methodological considerations

While linear mixed-effects modeling offers a powerful framework for analyzing representational similarity measures, it is critical that researchers carefully construct and validate their models. In our simulations, we designed our model RSM to be the “true” RSM for demonstration purposes. However, researchers should always consider if their models match the goals of their analysis, including 1) constructing the random effects structure that will converge in their dataset and 2) testing their model fits against alternative structures (Meteyard & Davies, 2020; Park et al., 2020) and 3) considering which effects should be considered random or fixed depending on their research question. An effect is fixed when the levels represent the specific conditions of theoretical interest (e.g., *task condition*) and the goal is to estimate and interpret those differences directly. In contrast, an effect is random when the levels are sampled from a broader population (e.g., *subjects*) and the goal is to account for their variability while generalizing beyond the sample tested. Note that the same variable (e.g., *stimuli*) may be considered fixed or random depending on the research questions.

One limitation of the proposed tRSA framework is that row-wise correlations are derived from a common representational similarity matrix and therefore are not strictly independent. This dependency is inherent to analyses based on representational similarity matrices and is not unique to tRSA. Although our linear mixed-effects model accounts for major sources of dependence through random effects for participants and stimuli, the shared-information structure among row-wise observations may vary across datasets and experimental designs. In the present study, simulations, empirical analyses, and additional permutation-based tests demonstrated that this dependency did not inflate false-positive rates or meaningfully alter statistical conclusions. We recommend that researchers applying tRSA evaluate the covariance structure of their own data and, where appropriate, complement mixed-effects analyses with permutation-based inference to verify the robustness of statistical conclusions. To ensure that the model assumptions were satisfied, we conducted a series of diagnostic checks on an exemplar ROI (right LOC; middle occipital gyrus) in the Object Perception dataset, including visual inspection of residual distributions and autocorrelation (Appendix 3, Figure 13). These diagnostics supported the assumptions of normality, homoscedasticity, and conditional independence of residuals. In addition, we conducted permutation-based inference, similar to prior improvements to cRSA (Nili et al., 2014), using a nested model comparison to test whether the mean similarity in this ROI was significantly greater than zero. The observed likelihood ratio test statistic fell in the extreme tail of the null distribution (Appendix 3, Figure 14), providing strong nonparametric evidence for the reliability of the observed effect. We emphasize that this type of model checking and permutation testing is not merely confirmatory but can help validate key assumptions in RSA modeling, especially when applying mixed-effects models to neural similarity data.

Nevertheless, because mixed-effects models may not fully account for all forms of dependence or model misspecification, this issue remains a consideration in principle. In settings where assumptions about the error structure are uncertain or difficult to verify, permutation-based inference provides a valuable complementary approach because it does not rely on parametric assumptions and can offer more robust significance testing.

Additionally, we note that while our proposed tRSA framework provides a flexible and statistically principled approach for modeling trial-level representational strength, we acknowledge that there are alternative methods for addressing trial-level variability in RSA. In particular, the use of cross-validated distance metrics (e.g., crossnobis distance) has become increasingly popular for controlling differences in measurement noise variance and accounting for possible covariance structures across trials (Walther et al., 2016). These metrics offer several advantages, including unbiased estimation of representational dissimilarities under Gaussian noise assumptions and improved generalization to unseen data. However, cross-validated distances are conceptually distinct from the approach taken here: whereas cross-validation aims to correct for noise-related biases in representational dissimilarity matrices, our trial-level RSA method focuses on estimating and modeling the variability in representation strength across individual trials using mixed-effects modeling. Rather than proposing a replacement for cross-validated RSA, tRSA adds a complementary tool to the methodological toolkit—one that supports hypothesis-driven inference about condition effects and trial-level covariates, while leveraging the full structure of the data.

## Conclusion

We present a comprehensive overview and diagnostics of a trial-level approach to representational similarity analysis, termed tRSA. With both simulations and real fMRI datasets, we have demonstrated that tRSA properly captures the multi-level variance structure in the data, shows improved and robust specificity and sensitivity, and offers flexible modeling options. We believe that tRSA is an important advancement of the generic RSA implementation, both methodologically and conceptually, and we hope that this innovation provides a versatile and useful tool for digging deeper into the fascinating complexity of cognition.

## Acknowledgement

This study was supported by the National Institutes of Health grant numbers: R01-AG066901, R21-AG058161, and R01-AG075417.

## Appendices

### Appendix 1. Row-wise correlations

We explored the statistical properties of computing row-wise correlations, which is a critical step in tRSA. Assuming an experimental design with *n* = 200 trials, we sampled *n*(*n* − 1)/2 = 19900 pairs of values from a bivariate normal distribution, with their Pearson correlation fixed at some given value via the ‘MASS 7.3-60.2’ package (Venables et al., 2002) in R. The two vectors were rearranged as the lower triangular part of two *n* × *n* matrices, and the upper triangular parts were filled out accordingly to yield two symmetric matrices *M* and *S*. With each pair of matrices, the Fisher-transformed Pearson correlation coefficient of the lower triangular parts of the two matrices is equal to the ground truth parameter *z*. We computed row-wise Pearson correlations for row vectors *M_i_*_·_ and *S_i_*_·_, barring entries on the diagonal (see **Figure 1C, Bottom**). Fisher transformation of correlation coefficients was applied such that the distribution of values is more approximately normal (Silver & Dunlap, 1987). This set of procedures was repeated for 10,000 iterations, for each ground truth parameter *z* ranging from −0.2 to 0.6 with increments of 0.1.Our simulations indicated that the average of row-wise correlations numerically approximates the overall correlation. Specifically, the estimates centered around zero when it was the ground truth; as the magnitude of the ground truth increased, there appeared to be a bias toward slightly larger magnitudes. Despite this, the spread of estimates derived from the row-wise method remained small, with a constrained range of [0.599, 0.604] around the ground truth of 0.600 (see **Appendix 11 Figure 11**).

**Appendix 1 Figure 11.**
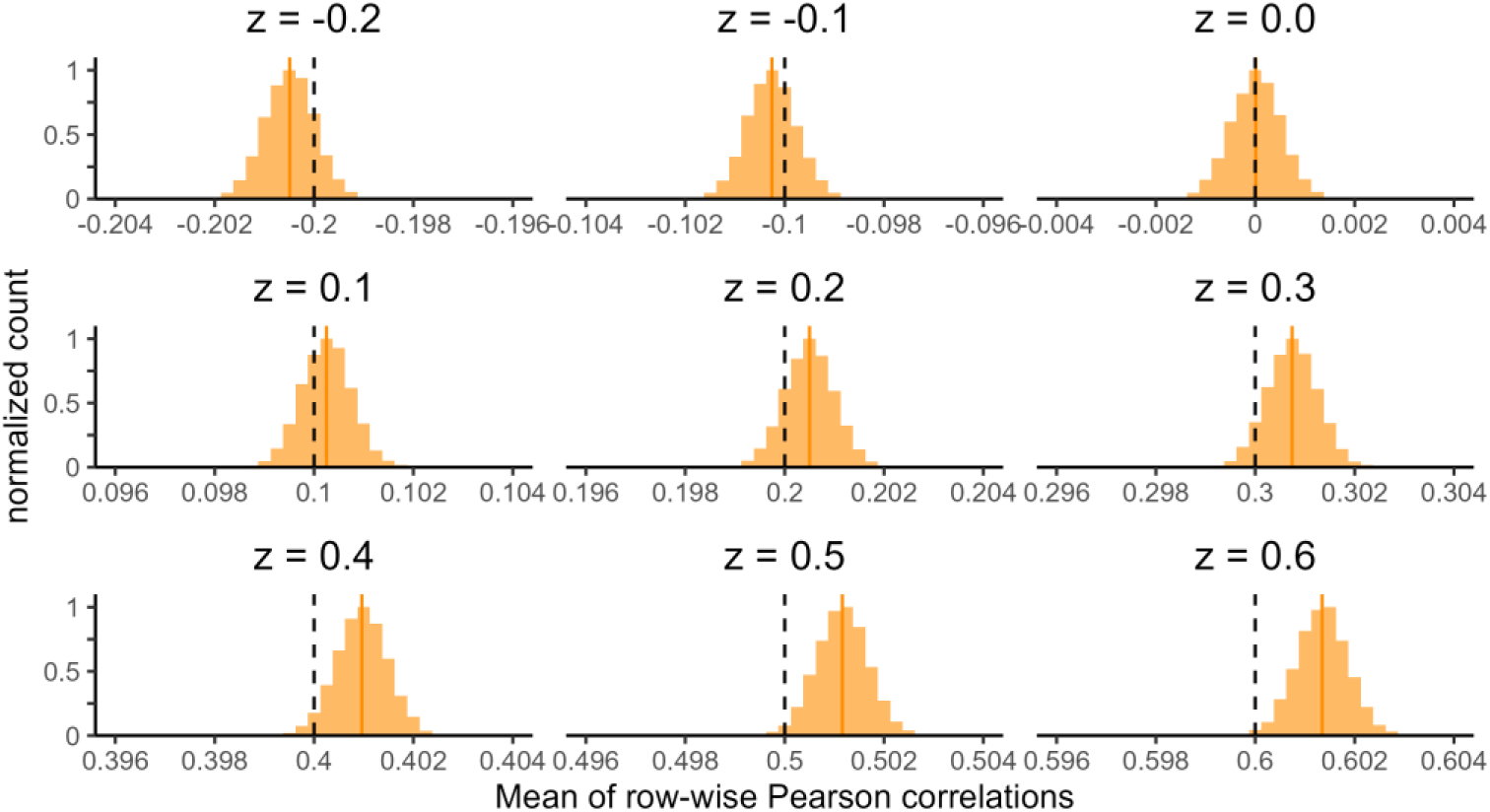
Row-wise correlations. Histograms of the mean of row-wise correlation values. Each histogram depicted 10,000 iterations. In each iteration, two symmetrical matrices with a fixed ground truth Pearson correlation coefficient were randomly generated, and the mean of row-wise correlation coefficients were computed as the estimate. Dashed vertical lines indicate the ground truth parameter, and solid vertical lines indicate the mean of estimates.

Note that a limitation of this simulation is that the symmetric matrices were randomly generated without being guaranteed to be mathematically valid similarity matrices (i.e., positive semidefinite). For example, [[1, 1, 1], [1, 1, −1], [1, −1, 1]] is symmetric but is not a valid correlation RSM, since it is impossible to satisfy *a* = *b* = *c* (row 1) and *a* = *b* = −*c* (row 2) unless *a* = *b* = *c* = 0, in which case the Pearson correlation is undefined. Similarly, RSMs using distance metrics such as Euclidean distance must satisfy the triangle inequality, i.e., *d_xy_* + *d_xz_* ≥ *d_yz_*. Moreover, it is difficult to implement the multi-level variance structure directly on RSMs; rather, those variances are theorized to operate directly on the underlying *activity patterns*.

### Appendix 2. Comparing across-condition and within-condition tRSA

Given discrete experimental conditions, tRSA could be formulated either within each condition or across all conditions. In *within-condition* tRSA, one would generate condition-specific RSMs, e.g., one set of *n_A_* × *n_A_* RSMs for Condition A and another set of *n_B_* × *n_B_* RSMs for Condition B, and compute the two sets of *within-condition* tRSA values separately. These tRSA estimates are then subjected to subsequent analysis. In *across-condition* tRSA, RSMs are generated across all trials in all conditions, and tRSA estimates are computed as if there were no multiple conditions. Then, tRSA estimates are categorized based on the condition they belong to in any subsequent analysis.

We advocate for the use of across-condition tRSA for its enhanced statistical reliability over within-condition tRSA. One major limitation of within-condition tRSA is its vulnerability to small and variable trial counts. Suppose that a condition contains *n* trials, cRSA would use *n*(*n* − 1)/2 observations to compute its estimate, whereas within-condition tRSA could only use *n* − 1 observations. This effective sample size could turn out to be fairly small and variable in practice (see **Figure 8A, Bottom**), resulting in unreliable estimates (see **Figure 2**). On the contrary, across-condition tRSA suffers much less from this issue, since most experiments should have been designed to have an adequate total trial count.

Another theoretical advantage of across-condition tRSA is that it better corresponds to tRSA_continuous_, i.e., analyses in which trial-level representational strength estimates are hypothesized to be modulated by some continuous variable such as memorability or reaction time. In both across-condition tRSA and tRSA_continuous_, full RSMs are used to estimate representational strength before any discrete condition effect or continuous modulation effect is assessed.

**Appendix 2 Figure 12** below summarizes the differences between within-condition and across-condition tRSA outcomes in a series of simulations. Notably, while within-condition tRSA estimates showed better numerical correspondence with cRSA estimates, they are less reliable than across-condition tRSA, as evidenced by the wider spread of estimates.

**Appendix 2 Figure 12.**
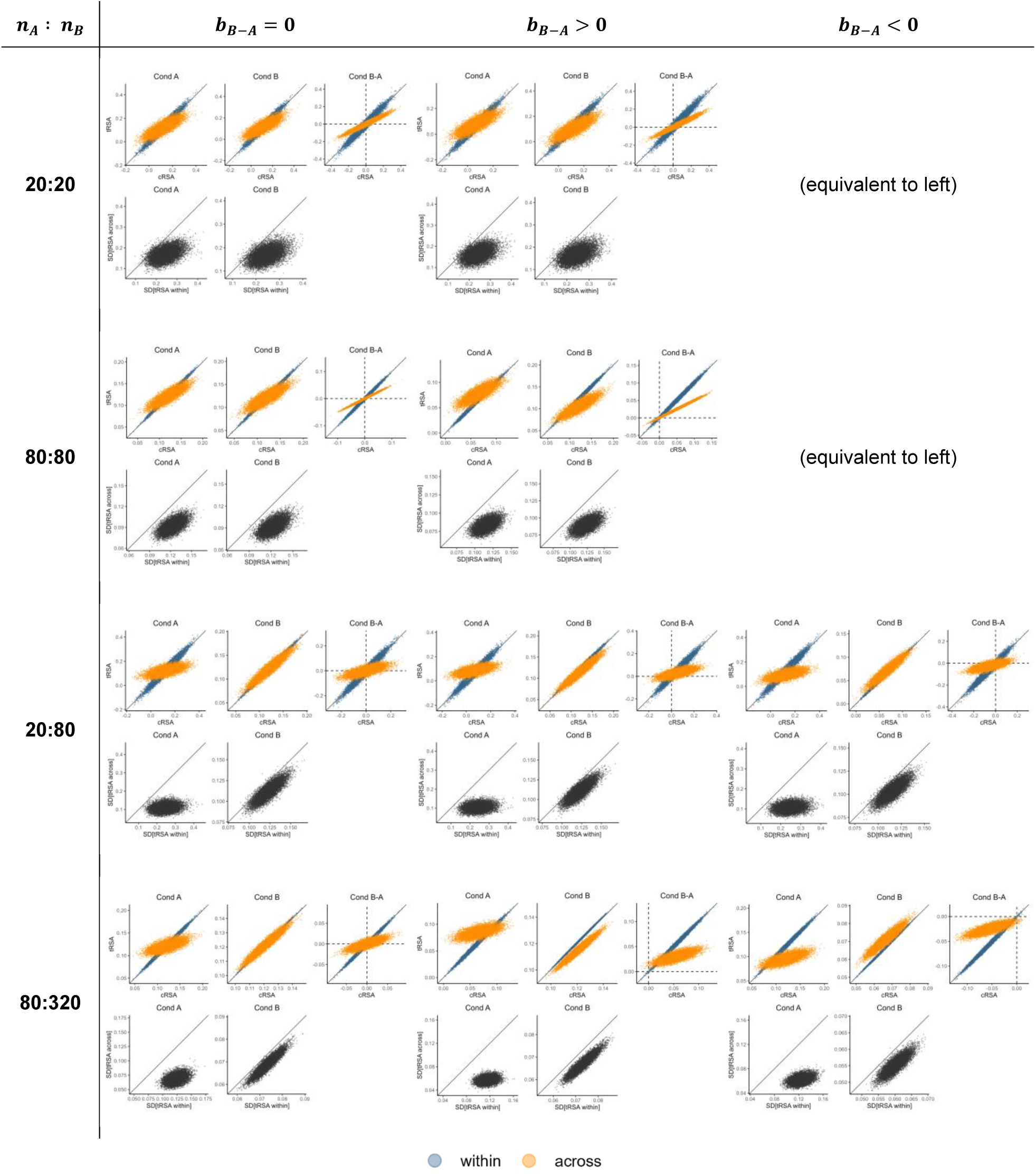
Comparing across-condition and within-condition tRSA. Each cell depicts descriptive statistics of different RSA methods from 10,000 simulations. Cells differ in raw trial counts (20, 80, or 320 in a given condition), balance of trial counts between conditions (balanced or unbalanced), and effect (A = B, A > B, or A < B). In each cell, the top scatter plots depict condition-level tRSA estimates (y-axis; blue, within-condition; orange, across-condition) against cRSA estimates (x-axis), whereas the bottom scatter plots depict the standard deviations of trial-level representational strength estimates from across-condition tRSA (y-axis) and within-condition tRSA (x-axis).

### Appendix 3. Supporting data and results for Experiment 2

#### Exemplar Permutation Testing

To test whether the mean representational strength in the ROI right LOC (middle occipital gyrus) was significantly greater than zero, we used a permutation-based likelihood ratio test implemented via the ‘permlmer’ function. This test compares two nested linear mixed-effects models fit using the ‘lmer’ function from the lme4 package, both including random intercepts for Participant and Stimulus ID to account for between-subject and between-item variability.

The null model excluded a fixed intercept term, effectively constraining the mean similarity to zero after accounting for random effects:

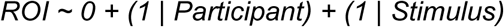

The full model included the same random effects structure but allowed the intercept to be freely estimated:

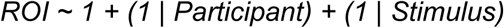

By comparing the fit of these two models, we directly tested whether the average similarity in this ROI was significantly different from zero. Permutation testing (1,000 permutations) was used to generate a nonparametric p-value, providing inference without relying on normality assumptions.

The full model, which estimated a nonzero mean similarity in the right LOC (middle occipital gyrus), showed a significantly better fit to the data than the null model that fixed the mean at zero (χ²(1) = 17.60, p = 2.72 × 10⁻⁵). The permutation-based p-value obtained from permlmer confirmed this effect as statistically significant (p = 0.0099), indicating that the mean similarity in this ROI was reliably greater than zero. These results support the conclusion that the right LOC contains representational structure consistent with the HMAXc2 RSM. A density plot of the permuted likelihood ratio tests is plotted along with the observed likelihood ratio test in Appendix 3 Figure 14.

**Figure 13.**
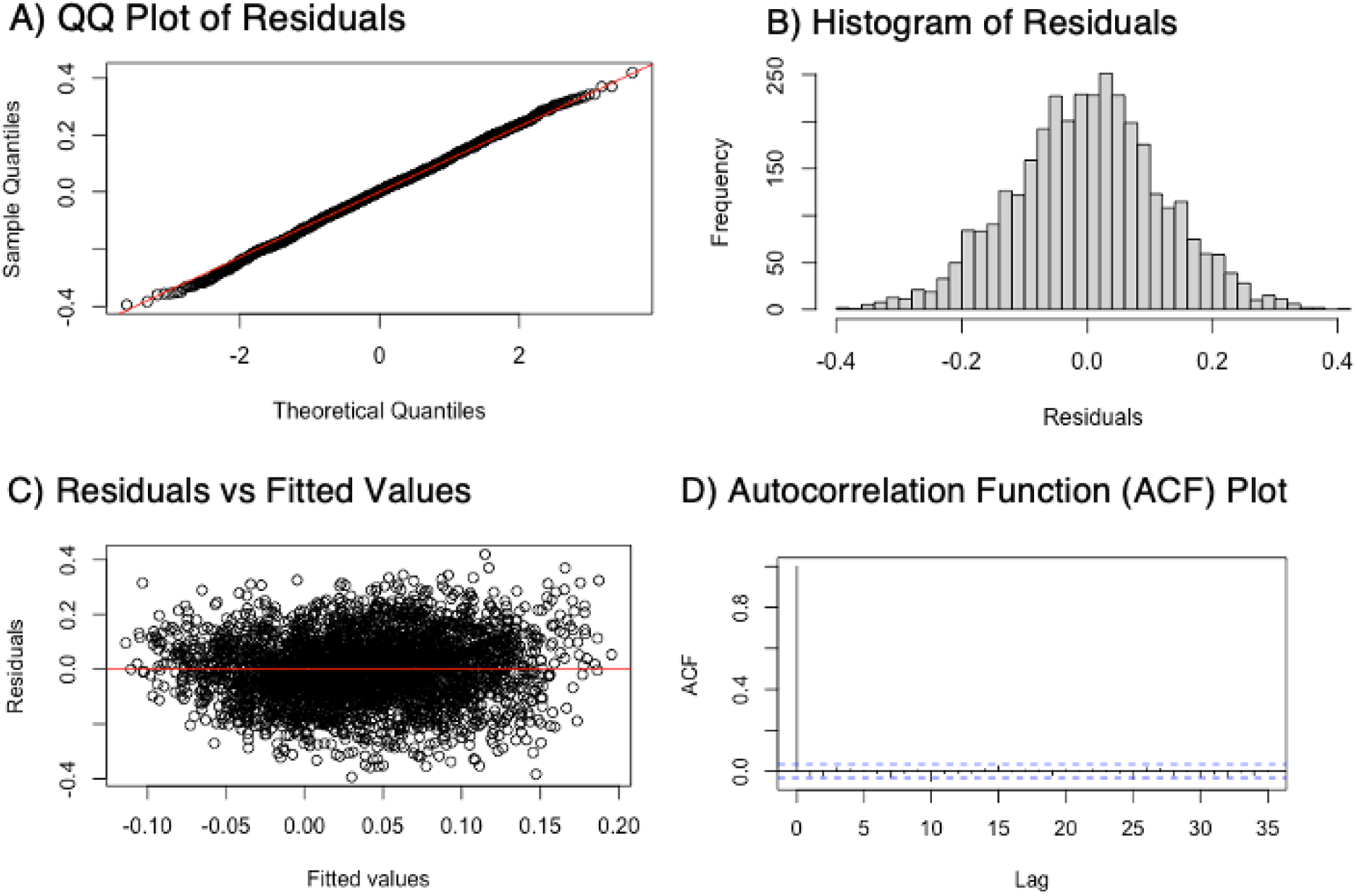
Exemplar model checks Experiment 2 Object Perception in right LOC (middle occipital gyrus). A) QQ Plot of Residuals. B) Histogram of Residuals. C) Residuals vs Fitted Values. D) Autocorrelation Function (ACF) Plot. These diagnostic plots were generated to evaluate whether the linear mixed-effects model satisfied key statistical assumptions. The model was fit to an exemplar region of interest (ROI) with subject and stimulus modeled as random intercepts. Residuals were extracted from the fitted model and plotted using base R functions. The QQ plot (A) and histogram (B) indicate that residuals are approximately normally distributed. The residuals vs. fitted values plot (C) shows no clear patterns or funneling, suggesting homoscedasticity and no evidence of model misspecification. Finally, the ACF plot (D) displays no significant autocorrelation, supporting the assumption that residuals are independent after accounting for the specified random effects structure.

**Appendix 3 Figure 14.**
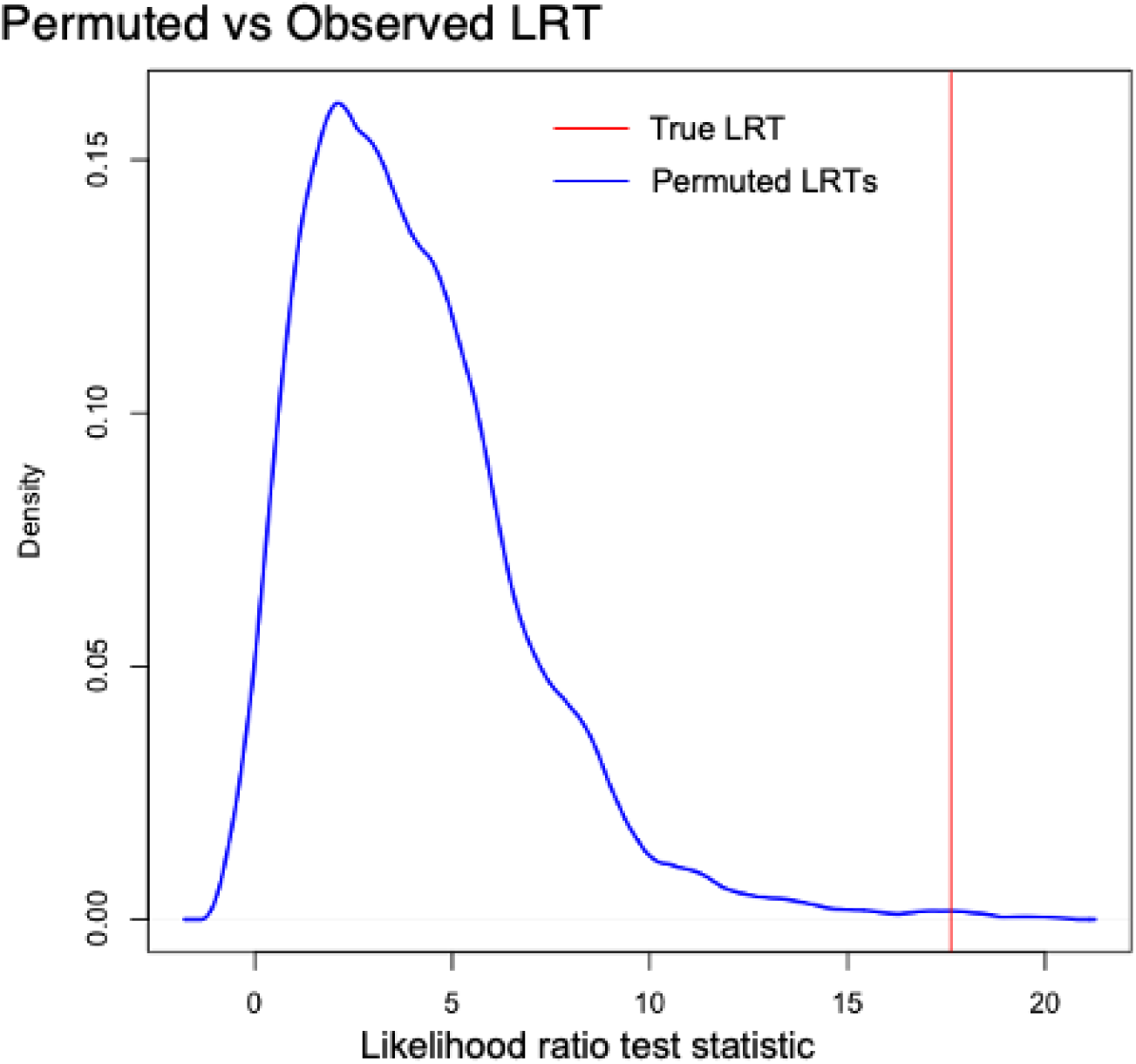
Permutation-based likelihood ratio test (LRT) comparing models with and without a fixed intercept in the right LOC (middle occipital gyrus). The x-axis shows the distribution of LRT statistics obtained from 1,000 permutations under the null hypothesis that the mean similarity is zero. The blue line represents the null distribution of permuted LRT values, and the red vertical line indicates the observed LRT statistic from the original model comparison (χ² = 17.60). The observed value falls in the extreme tail of the null distribution, yielding a permutation-based p-value of 0.0099, indicating that the mean similarity in this ROI is significantly greater than zero.”

**Appendix 3 Table 4.** Behavioral results. Bold indicates participants excluded from Conceptual Retrieval Analyses.

| Participant | Gender | Response Trials at Perception | ROC | Count of Hits | Hit Rate |
| --- | --- | --- | --- | --- | --- |
| 1 | M | 114 | 0.84 | 84 | 0.74 |
| 2 | W | 114 | 0.73 | 82 | 0.78 |
| **3** | **W** | 105 | **0.70** | **32** | **0.36** |
| 4 | W | 112 | 0.93 | 94 | 0.82 |
| 5 | W | 114 | 0.95 | 85 | 0.76 |
| 6 | W | 96 | 0.74 | 84 | 0.79 |
| 7 | W | 108 | 0.93 | 99 | 0.88 |
| 8 | W | 114 | 0.56 | 76 | 0.75 |
| 9 | M | 113 | 0.81 | 89 | 0.78 |
| 10 | M | 101 | 0.79 | 64 | 0.80 |
| 11 | M | 101 | 0.93 | 78 | 0.80 |
| 12 | W | 113 | 0.66 | 53 | 0.50 |
| 13 | M | 114 | 0.87 | 89 | 0.78 |
| 14 | W | 103 | 0.69 | 35 | 0.39 |
| 15 | M | 114 | 0.84 | 75 | 0.66 |
| 16 | W | 114 | 0.67 | 51 | 0.45 |
| **17** | **W** | **113** | **0.35** | **26** | **0.30** |
| 18 | W | 114 | 0.83 | 75 | 0.66 |
| 19 | W | 114 | 0.90 | 100 | 0.88 |
| 20 | W | 114 | 0.71 | 68 | 0.60 |
| 21 | W | 114 | 0.89 | 80 | 0.70 |
| 22 | W | 114 | 0.90 | 99 | 0.89 |
| 23 | W | 113 | 0.83 | 93 | 0.82 |
| 24 | W | 110 | 0.70 | 64 | 0.62 |
| 25 | M | 114 | 0.87 | 84 | 0.74 |
| 26 | M | 95 | 0.62 | 71 | 0.75 |
| 27 | M | 114 | 0.75 | 81 | 0.71 |
| 28 | M | 109 | 0.95 | 100 | 0.88 |
| 29 | W | 106 | 0.72 | 81 | 0.79 |
| 30 | W | 114 | 0.92 | 95 | 0.86 |
| 31 | M | 114 | 0.74 | 79 | 0.69 |
| 32 | W | 109 | 0.62 | 50 | 0.50 |

**Appendix 3 Table 5.** Model selection using 3 criteria: Akaike information criterion (AIC), Bayesian information criterion (BIC), and log-liklihood ratio testing (LRT).

| REGION | SubArea | Coordinates<br>(x, y, z) | (1+adjHvM\|Subject +<br>(1+adjHvM\|StimID) |  |  | (1+adjHvM\|Subject)+(1\|StimID) |  |  | (1\|Subject)+(1\|StimID) |  |  |
| --- | --- | --- | --- | --- | --- | --- | --- | --- | --- | --- | --- |
|  |  |  | AIC | BIC | LRT<br>pvalue | AIC | BIC | LRT<br>pvalue | AIC | BIC | LRT<br>pvalue |
| LEFT IPL | A40c, caudal area<br>40(PFm) | -56, -49, 38 | -4778 | -4724 | - | -4781 | -4739 | 0.555 | -4784 | -4754 | 0.600 |
| RIGHT<br>IPL | A40c, caudal area<br>40(PFm) | 57, -44, 38 | -4632 | -4578 | - | -4614 | -4571 | 0.000 | -4639 | -4608 | 1.000 |
| LEFT IPL | A39rv, rostroventral<br>area 39(PGa) | -47, -65, 26 | -4907 | -4852 | - | -4910 | -4867 | 0.674 | -4914 | -4883 | 0.819 |
| RIGHT<br>IPL | A39rv, rostroventral<br>area 39(PGa) | 53, -54, 25 | -4765 | -4710 | - | -4768 | -4726 | 0.909 | -4772 | -4742 | 0.969 |
| LEFT ITC | A20iv, intermediate<br>ventral area 20 | -45, -26, -27 | -4786 | -4731 | - | -4790 | -4747 | 1.000 | -4788 | -4758 | 0.079 |
| LEFT ITC | A20r, rostral area<br>20 | -43, -2, -41 | -4188 | -4134 | - | -4187 | -4145 | 0.117 | -4191 | -4161 | 0.858 |
| LEFT ITC | A20il, intermediate<br>lateral area 20 | -56, -16, -28 | -4808 | -4753 | - | -4806 | -4764 | 0.064 | -4809 | -4779 | 0.663 |
| RIGHT<br>ITC | A20iv, intermediate<br>ventral area 20 | 46, -14, -33 | -4526 | -4472 | - | -4529 | -4487 | 0.623 | -4533 | -4502 | 0.716 |
| RIGHT<br>ITC | A20r, rostral area<br>20 | 40, 0, -43 | -3940 | -3886 | - | -3942 | -3901 | 0.510 | -3944 | -3914 | 0.337 |
| RIGHT<br>ITC | A20il, intermediate<br>lateral area 20 | 55, -11, -32 | -4554 | -4500 | - | -4558 | -4516 | 0.982 | -4562 | -4532 | 0.974 |
| LEFT ITC | A37elv, extreme<br>lateroventral area37 | -51, -57, -15 | -4920 | -4865 | - | -4924 | -4881 | 1.000 | -4924 | -4894 | 0.214 |
| LEFT ITC | A37vl, ventrolateral<br>area 37 | -55, -60, -6 | -4946 | -4891 | - | -4949 | -4907 | 0.875 | -4953 | -4923 | 0.875 |
| LEFT ITC | A20cl, caudolateral<br>of area 20 | -59, -42, -16 | -5003 | -4949 | - | -5005 | -4962 | 0.297 | -5007 | -4977 | 0.476 |
| LEFT ITC | A20cv, caudoventral of<br>area 20 | -55, -31, -27 | -4867 | -4812 | - | -4871 | -4828 | 0.996 | -4873 | -4843 | 0.413 |
| RIGHT<br>ITC | A37elv, extreme<br>lateroventral area37 | 53, -52, -18 | -4933 | -4878 | - | -4935 | -4892 | 0.301 | -4936 | -4906 | 0.352 |
| RIGHT<br>ITC | A37vl, ventrolateral<br>area 37 | 54, -57, -8 | -4875 | -4821 | - | -4877 | -4835 | 0.366 | -4881 | -4850 | 0.865 |
| RIGHT<br>ITC | A20cl, caudolateral<br>of area 20 | 61, -40, -17 | -4617 | -4563 | - | -4619 | -4576 | 0.291 | -4623 | -4592 | 0.966 |
| RIGHT<br>ITC | A20cv, caudoventral of<br>area 20 | 54, -31, -26 | -4683 | -4628 | - | -4685 | -4642 | 0.293 | -4688 | -4657 | 0.679 |
| LEFT<br>LOC | mOccG, middle<br>occipital gyrus | -31, -89, 11 | -5123 | -5068 | - | -5127 | -5084 | 0.941 | -5128 | -5097 | 0.259 |
| LEFT<br>LOC | V5/MT+, area<br>V5/MT+ | -46, -74, 3 | -4820 | -4765 | - | -4824 | -4782 | 0.996 | -4827 | -4797 | 0.631 |
| LEFT<br>LOC | OPC, occipital polar<br>cortex | -18, -99, 2 | -5046 | -4991 | - | -5049 | -5006 | 0.595 | -5052 | -5022 | 0.818 |
| LEFT<br>LOC | iOccG, inferior<br>occipital gyrus | -30, -88, -12 | -4959 | -4904 | - | -4962 | -4919 | 0.516 | -4966 | -4935 | 0.929 |
| **RIGHT LOC** | mOccG, middle occipital gyrus | 34, -86, 11 | -5265 | -5210 | - | -5269 | -5227 | 1.000 | -5271 | -5241 | 0.440 |
| **RIGHT LOC** | V5/MT+, area V5/MT+ | 48, -70, -1 | -4879 | -4824 | - | -4881 | -4839 | 0.382 | -4883 | -4852 | 0.279 |
| **RIGHT LOC** | OPC, occipital polar cortex | 22, -97, 4 | -4995 | -4940 | - | -4989 | -4946 | 0.007 | -4991 | -4961 | 0.472 |
| **RIGHT LOC** | iOccG, inferior occipital gyrus | 32, -85, -12 | -5126 | -5071 | - | -5126 | -5083 | 0.144 | -5129 | -5098 | 0.618 |
| **% Selected** |  |  | **4%** | **0%** | **8%** | **0%** | **0%** | **0%** | **96%** | **100%** | **92%** |

**Appendix 3 Table 6.** Significant nuisance effects in memorability model.

| REGION | SubArea | Coordinates<br>(x, y, z) | Conceptual Memorability |  |  |  |  | Reaction Time |  |  |  |  |
| --- | --- | --- | --- | --- | --- | --- | --- | --- | --- | --- | --- | --- |
|  |  |  | Mean | SE | t Values | p | q | coeff | SE | t Values | p | q |
| **LEFT IPL** | A40c, caudal area 40(PFm) | -56, -49, 38 | 0.077 | 0.060 | 1.274 | 0.203 | 0.659 | 0.005 | 0.004 | 1.369 | 0.171 | 0.869 |
| **RIGHT IPL** | A40c, caudal area 40(PFm) | 57, -44, 38 | -0.053 | 0.057 | -0.931 | 0.352 | 0.773 | 0.007 | 0.004 | 1.731 | 0.084 | 0.869 |
| **LEFT IPL** | A39rv, rostroventral area 39(PGa) | -47, -65, 26 | -0.019 | 0.056 | -0.341 | 0.733 | 0.866 | 0.005 | 0.004 | 1.433 | 0.152 | 0.869 |
| **RIGHT IPL** | A39rv, rostroventral area 39(PGa) | 53, -54, 25 | -0.039 | 0.059 | -0.666 | 0.506 | 0.773 | 0.010 | 0.004 | 2.482 | 0.013 | 0.869 |
| **LEFT ITC** | A20iv, intermediate ventral area 20 | -45, -26, -27 | -0.052 | 0.059 | -0.876 | 0.381 | 0.773 | 0.004 | 0.004 | 1.101 | 0.271 | 0.869 |
| **LEFT ITC** | A20r, rostral area 20 | -43, -2, -41 | -0.051 | 0.065 | -0.780 | 0.436 | 0.773 | -0.004 | 0.004 | -0.846 | 0.397 | 0.869 |
| **LEFT ITC** | A20il, intermediate lateral area 20 | -56, -16, -28 | -0.053 | 0.060 | -0.893 | 0.372 | 0.773 | 0.004 | 0.004 | 0.889 | 0.374 | 0.869 |
| **RIGHT ITC** | A20iv, intermediate ventral area 20 | 46, -14, -33 | 0.017 | 0.061 | 0.278 | 0.781 | 0.867 | 0.002 | 0.004 | 0.556 | 0.578 | 0.869 |
| **RIGHT ITC** | A20r, rostral area 20 | 40, 0, -43 | 0.029 | 0.065 | 0.444 | 0.657 | 0.854 | 0.000 | 0.004 | 0.016 | 0.987 | 0.869 |
| **RIGHT ITC** | A20il, intermediate lateral area 20 | 55, -11, -32 | -0.053 | 0.059 | -0.887 | 0.375 | 0.773 | 0.002 | 0.004 | 0.439 | 0.661 | 0.869 |
| **LEFT ITC** | A37elv, extreme lateroventral area 37 | -51, -57, -15 | -0.044 | 0.059 | -0.750 | 0.453 | 0.773 | 0.004 | 0.004 | 1.050 | 0.294 | 0.869 |
| **LEFT ITC** | A37vl, ventrolateral area 37 | -55, -60, -6 | -0.043 | 0.062 | -0.682 | 0.495 | 0.773 | -0.003 | 0.004 | -0.627 | 0.531 | 0.869 |
| **LEFT ITC** | A20cl, caudolateral of area 20 | -59, -42, -16 | 0.009 | 0.057 | 0.165 | 0.869 | 0.869 | 0.002 | 0.004 | 0.529 | 0.597 | 0.869 |
| **LEFT**<br>**ITC** | A20cv,<br>caudoventral of<br>area 20 | -55, -31, -27 | 0.032 | 0.057 | 0.550 | 0.582 | 0.797 | 0.001 | 0.004 | 0.218 | 0.827 | 0.869 |
| **RIGHT**<br>**ITC** | A37elv,<br>extreme<br>lateroventral<br>area37 | 53, -52, -18 | 0.034 | 0.059 | 0.580 | 0.562 | 0.797 | 0.000 | 0.004 | 0.015 | 0.988 | 0.869 |
| **RIGHT**<br>**ITC** | A37vl,<br>ventrolateral<br>area 37 | 54, -57, -8 | -0.012 | 0.059 | -0.210 | 0.834 | 0.867 | -0.007 | 0.004 | -1.662 | 0.097 | 0.869 |
| **RIGHT**<br>**ITC** | A20cl,<br>caudolateral of<br>area 20 | 61, -40, -17 | 0.022 | 0.060 | 0.366 | 0.714 | 0.866 | -0.001 | 0.004 | -0.137 | 0.891 | 0.869 |
| **RIGHT**<br>**ITC** | A20cv,<br>caudoventral of<br>area 20 | 54, -31, -26 | 0.013 | 0.060 | 0.221 | 0.825 | 0.867 | -0.004 | 0.004 | -0.909 | 0.363 | 0.869 |
| **LEFT**<br>**LOC** | mOccG, middle<br>occipital gyrus | -31, -89, 11 | **-0.233** | **0.067** | **-3.473** | **0.001** | **0.005** | **0.015** | **0.005** | **3.199** | **0.001** | **0.033** |
| **LEFT**<br>**LOC** | V5/MT+, area<br>V5/MT+ | -46, -74, 3 | **-0.193** | **0.059** | **-3.241** | **0.001** | **0.006** | **0.008** | **0.004** | **2.093** | **0.036** | **0.033** |
| **LEFT**<br>**LOC** | OPC, occipital<br>polar cortex | -18, -99, 2 | **-0.259** | **0.074** | **-3.516** | **0.000** | **0.005** | **0.017** | **0.005** | **3.359** | **0.001** | **0.033** |
| **LEFT**<br>**LOC** | iOccG, inferior<br>occipital gyrus | -30, -88, -12 | -0.054 | 0.069 | -0.779 | 0.436 | 0.773 | 0.011 | 0.005 | 2.371 | 0.018 | 0.869 |
| **RIGHT**<br>**LOC** | mOccG, middle<br>occipital gyrus | 34, -86, 11 | **-0.224** | **0.068** | **-3.286** | **0.001** | **0.006** | **0.015** | **0.005** | **3.361** | **0.001** | **0.033** |
| **RIGHT**<br>**LOC** | V5/MT+, area<br>V5/MT+ | 48, -70, -1 | -0.089 | 0.064 | -1.388 | 0.165 | 0.614 | 0.006 | 0.004 | 1.311 | 0.190 | 0.869 |
| **RIGHT**<br>**LOC** | OPC, occipital<br>polar cortex | 22, -97, 4 | **-0.296** | **0.074** | **-3.987** | **0.000** | **0.002** | **0.014** | **0.005** | **2.867** | **0.004** | **0.033** |
| **RIGHT**<br>**LOC** | iOccG, inferior<br>occipital gyrus | 32, -85, -12 | -0.099 | 0.069 | -1.439 | 0.150 | 0.614 | 0.012 | 0.005 | 2.610 | 0.009 | 0.869 |

### Appendix 4. Validation in an Independent Object Naming Dataset

#### Representational Similarity Matrices

The dataset consisted of 131 object images drawn from six semantic categories: animals, fruits, vegetables, vehicles, tools, and musical instruments. For the model RSM, a categorical model was constructed based on the six semantic categories. Stimulus pairs belonging to the same semantic category were maximally similar (1), whereas stimulus pairs belonging to different categories were maximally dissimilar (0). Diagonal elements, as is typical for both cRSA and tRSA approaches, were excluded from all analyses. Neural RSMs were created for 10 bilateral ventral-stream ROIs demonstrated to mediate object representation (Clarke et al., 2014; Devereux et al., 2018; Davis et al., 2021) spanning early visual cortex through anterior temporal cortex: left/right V1, V2, V3, posterior ventral temporal cortex (pVTC), and perirhinal cortex (PRC). For each participant and ROI, representational similarity matrices (RSMs) were computed from trial-level multivoxel activation patterns across the 131 stimuli. Neural similarity structure was compared against a categorical model RSM reflecting semantic category membership.

#### Representational Similarity Analysis

For cRSA, lower-triangular elements of each participant’s neural RSM were vectorized and correlated with the corresponding lower-triangular elements of the category model RSM using Spearman correlation. This produced a single similarity estimate for each participant and ROI. To test whether neural similarity structure significantly reflected categorical organization, one-sample one-tailed t-tests were conducted across participants separately for each ROI, testing whether cRSA similarity estimates were greater than zero.

For tRSA, similarity structure was quantified at the stimulus level. For each participant, ROI, and stimulus, the corresponding row of the neural RSM was correlated with the matching row of the category model RSM, excluding self-comparisons. This produced trial-level similarity estimates for each stimulus within each ROI and participant. Trial-level effects were analyzed using mixed-effects models with random intercepts Participant and Item.

False discovery rate (FDR) correction was applied separately to cRSA and tRSA analyses across the 10 ROIs using the Benjamini–Hochberg procedure (q < .05).

## Results

Both cRSA and tRSA revealed reliable category-level similarity structure across ventral visual regions. However, Across ROIs, tRSA produced effect sizes and statistical significance that were comparable to, and in several regions slightly larger than, cRSA. The strongest effects were observed in bilateral pVTC and early visual regions. In right pVTC, both methods produced highly robust category-model fits (cRSA: t = 5.46, tRSA: t = 5.47). Similarly strong effects were observed in right V1 (cRSA: t = 5.18, tRSA: t = 5.35) and right V2 (cRSA: t = 5.65, tRSA: t = 5.32). Both methods detected significant positive effects following FDR correction.

Across ROIs, cRSA and tRSA estimates were highly correlated, indicating strong agreement between the two approaches in the spatial distribution of representational similarity effects. Overall these findings are consistent with our main findings that tRSA reproduces the canonical category-level similarity organization previously observed along the ventral visual stream while providing a more flexible inferential framework.

**Appendix 4 Figure 15.**
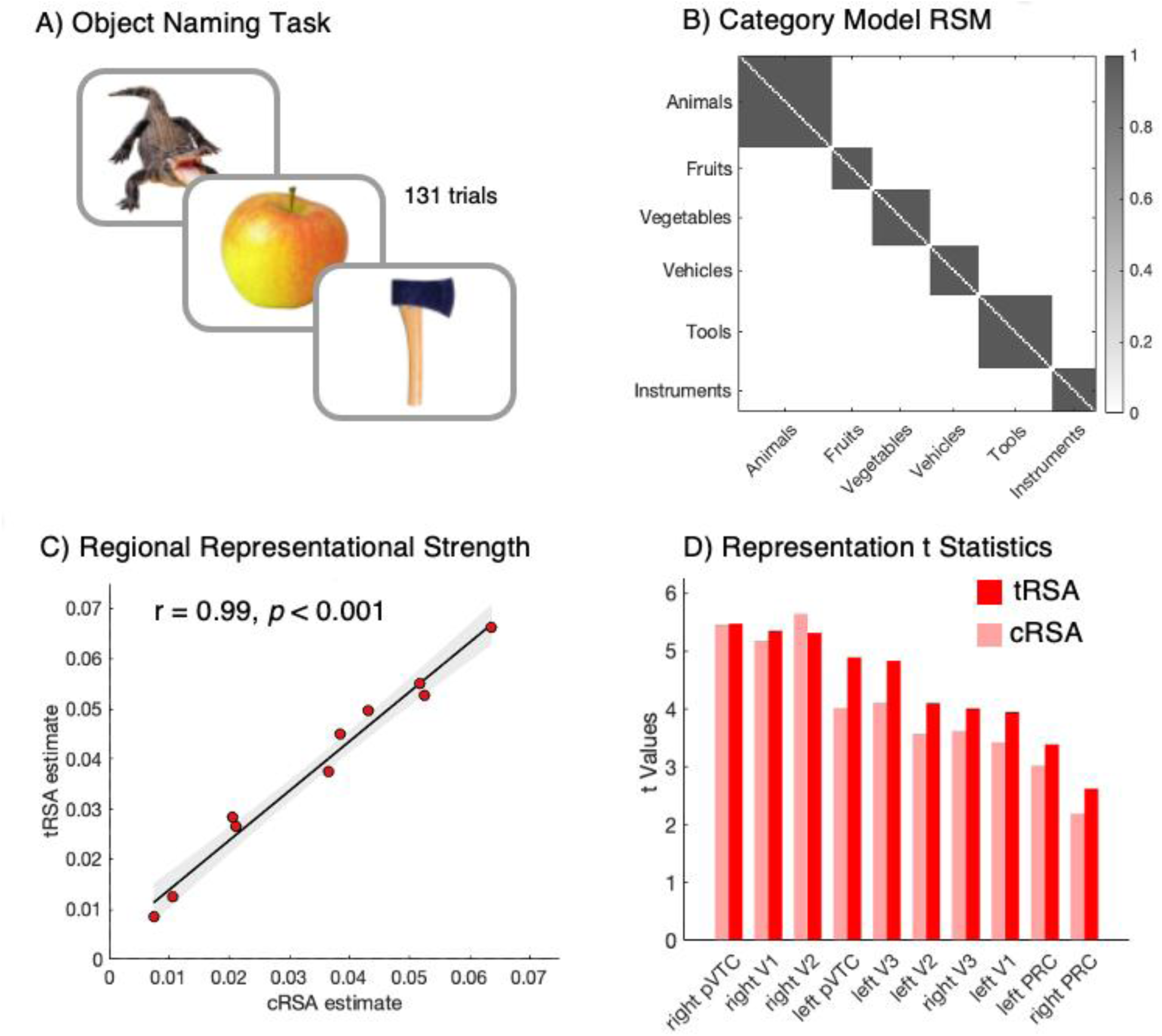
A) Object naming task from Clarke & Tyler, 2014, which used a distinct stimulus set from the studies considered in the main text. B) Category model RSM, emphasizing category membership. C) Correspondence between tRSA and cRSA representational strength across ROIs. D) Regional t-values obtained using tRSA (dark red) and cRSA (light red). All regions of interest show significant Neural-Model RSM correspondence when utilizing both methods.

**Appendix 4 Table 7.** cRSA and tRSA in an open Object Naming dataset.

| REGION | cRSA<br>coeff | cRSA<br>SE | cRSA t<br>Values | p | q | tRSA<br>coeff | tRSA<br>SE | tRSA t<br>Values | p | q |
| --- | --- | --- | --- | --- | --- | --- | --- | --- | --- | --- |
| RIGHT PVTC | 0.052 | 0.010 | 5.460 | < 0.001 | < 0.001 | 0.053 | 0.010 | 5.473 | < 0.001 | < 0.001 |
| RIGHT V1 | 0.064 | 0.012 | 5.184 | < 0.001 | < 0.001 | 0.066 | 0.012 | 5.351 | < 0.001 | < 0.001 |
| RIGHT V2 | 0.052 | 0.009 | 5.648 | < 0.001 | < 0.001 | 0.055 | 0.010 | 5.318 | < 0.001 | < 0.001 |
| LEFT PVTC | 0.038 | 0.010 | 4.021 | 0.001 | 0.001 | 0.045 | 0.009 | 4.892 | < 0.001 | < 0.001 |
| LEFT V3 | 0.021 | 0.005 | 4.108 | < 0.001 | 0.001 | 0.027 | 0.005 | 4.833 | < 0.001 | < 0.001 |
| LEFT V2 | 0.021 | 0.006 | 3.571 | 0.001 | 0.002 | 0.028 | 0.007 | 4.098 | < 0.001 | < 0.001 |
| **RIGHT V3** | 0.037 | 0.010 | 3.620 | 0.001 | 0.002 | 0.037 | 0.009 | 4.009 | < 0.001 | < 0.001 |
| **LEFT V1** | 0.043 | 0.013 | 3.424 | 0.002 | 0.002 | 0.050 | 0.013 | 3.948 | < 0.001 | < 0.001 |
| **LEFT PRC** | 0.011 | 0.004 | 3.024 | 0.004 | 0.005 | 0.012 | 0.004 | 3.389 | < 0.001 | < 0.001 |
| **RIGHT PRC** | 0.007 | 0.003 | 2.193 | 0.022 | 0.022 | 0.009 | 0.003 | 2.624 | 0.004 | 0.004 |

## Footnotes

1 *Representational geometry* refers to the pattern of similarities or dissimilarities among stimuli in a representational space, as captured by a representational similarity (or dissimilarity) matrix (Kriegeskorte & Kievit, 2013). This concept is independent of how the representational space is estimated and does not inherently require repeated stimulus presentations. Throughout this paper, we use the term representational strength to refer to the degree of correspondence between a neural representational geometry and a hypothesized model RSM (i.e., the first-level RSA correlation). We note that the term representational strength has been used differently in other contexts, and our definition is adopted here for clarity.

## References

Allen, E. J., St-Yves, G., Wu, Y., Breedlove, J. L., Prince, J. S., Dowdle, L. T., Nau, M., Caron, B., Pestilli, F., Charest, I., Hutchinson, J. B., Naselaris, T., & Kay, K. (2022). A massive 7T fMRI dataset to bridge cognitive neuroscience and artificial intelligence. Nature Neuroscience, 25(1), 116–126. 10.1038/s41593-021-00962-x

Aly, M., & Turk-Browne, N. B. (2016). Attention Stabilizes Representations in the Human Hippocampus. Cerebral Cortex, 26(2), 783–796. 10.1093/cercor/bhv041

Anzellotti, S., & Coutanche, M. N. (2018). Beyond Functional Connectivity: Investigating Networks of Multivariate Representations. Trends in Cognitive Sciences, 22(3), 258–269. 10.1016/j.tics.2017.12.002

Avants, B. B., Tustison, N. J., Song, G., Cook, P. A., Klein, A., & Gee, J. C. (2011). A reproducible evaluation of ANTs similarity metric performance in brain image registration. NeuroImage, 54(3), 2033–2044. 10.1016/j.neuroimage.2010.09.025

Baayen, R. H., Davidson, D. J., & Bates, D. M. (2008). Mixed-effects modeling with crossed random effects for subjects and items. *Journal of Memory and Language*, Special Issue: Emerging Data Analysis, 59(4), 390–412. 10.1016/j.jml.2007.12.005

Bainbridge, W. A. (2022). Memorability: Reconceptualizing memory as a visual attribute. In Visual memory (pp. 173–187). Routledge. 10.4324/9781003158134-11

Bainbridge, W. A., Dilks, D. D., & Oliva, A. (2017). Memorability: A stimulus-driven perceptual neural signature distinctive from memory. NeuroImage, 149, 141–152. 10.1016/j.neuroimage.2017.01.063

Bainbridge, W. A., & Rissman, J. (2018). Dissociating neural markers of stimulus memorability and subjective recognition during episodic retrieval. Scientific Reports, 8(1), 8679. 10.1038/s41598-018-26467-5

Baker, B., Lansdell, B., & Kording, K. P. (2022). Three aspects of representation in neuroscience. Trends in Cognitive Sciences, 26(11), 942–958. 10.1016/j.tics.2022.08.014

Barron, H. C., Garvert, M. M., & Behrens, T. E. J. (2016). Repetition suppression: A means to index neural representations using BOLD? Philosophical Transactions of the Royal Society B: Biological Sciences, 371(1705), 20150355. 10.1098/rstb.2015.0355

Bates, D., Mächler, M., Bolker, B., & Walker, S. (2015). Fitting Linear Mixed-Effects Models Using lme4. Journal of Statistical Software, 67(1), 1–48.

Bauer, A. J., & Just, M. A. (2019). Neural Representations of Concept Knowledge. In G. I. de Zubicaray & N. O. Schiller (Eds.), The Oxford Handbook of Neurolinguistics (p. 0). Oxford University Press. 10.1093/oxfordhb/9780190672027.013.21

Becker, M., Sommer, T., & Cabeza, R. (2024). Neural Mechanisms of Creative Problem Solving—From Representational Change to Memory Formation (p. 2023.06.13.544774). bioRxiv. 10.1101/2023.06.13.544774

Behzadi, Y., Restom, K., Liau, J., & Liu, T. T. (2007). A component based noise correction method (CompCor) for BOLD and perfusion based fMRI. NeuroImage, 37(1), 90–101. 10.1016/j.neuroimage.2007.04.042

Bobadilla-Suarez, S., Ahlheim, C., Mehrotra, A., Panos, A., & Love, B. C. (2020). Measures of Neural Similarity. Computational Brain & Behavior, 3(4), 369–383. 10.1007/s42113-019-00068-5

Brentano, F. (1874). Psychology From an Empirical Standpoint. Routledge.

Chang, N., Pyles, J. A., Marcus, A., Gupta, A., Tarr, M. J., & Aminoff, E. M. (2019). BOLD5000, a public fMRI dataset while viewing 5000 visual images. Scientific Data, 6(1), 49. 10.1038/s41597-019-0052-3

Charest, I., Kriegeskorte, N., & Kay, K. N. (2018). GLMdenoise improves multivariate pattern analysis of fMRI data. NeuroImage, 183, 606–616. 10.1016/j.neuroimage.2018.08.064

Chen, G., Padmala, S., Chen, Y., Taylor, P. A., Cox, R. W., & Pessoa, L. (2021). To pool or not to pool: Can we ignore cross-trial variability in FMRI? NeuroImage, 225, 117496. 10.1016/j.neuroimage.2020.117496

Cohen, J. (1983). The cost of dichotomization. Applied Psychological Measurement, 7(3), 249–253. 10.1177/014662168300700301

Cox, R. W., & Hyde, J. S. (1997). Software tools for analysis and visualization of fMRI data. NMR in Biomedicine, 10(4–5), 171–178. 10.1002/(SICI)1099-1492(199706/08)10:4/5%3C171::AID-NBM453%3E3.0.CO;2-L

Critchley, H. D., & Garfinkel, S. N. (2018). The influence of physiological signals on cognition. *Current Opinion in Behavioral Sciences*, Emotion-Cognition Interactions, 19, 13–18. 10.1016/j.cobeha.2017.08.014

Dale, A. M., Fischl, B., & Sereno, M. I. (1999). Cortical Surface-Based Analysis: I. Segmentation and Surface Reconstruction. NeuroImage, 9(2), 179–194. 10.1006/nimg.1998.0395

Davis, S. W., Geib, B. R., Wing, E. A., Wang, W.-C., Hovhannisyan, M., Monge, Z. A., & Cabeza, R. (2021). Visual and Semantic Representations Predict Subsequent Memory in Perceptual and Conceptual Memory Tests. Cerebral Cortex, 31(2), 974–992. 10.1093/cercor/bhaa269

Dedrick, R. F., Ferron, J. M., Hess, M. R., Hogarty, K. Y., Kromrey, J. D., Lang, T. R., Niles, J. D., & Lee, R. S. (2009). Multilevel Modeling: A Review of Methodological Issues and Applications. Review of Educational Research, 79(1), 69–102. 10.3102/0034654308325581

Devereux, B. J., Clarke, A., & Tyler, L. K. (2018). Integrated deep visual and semantic attractor neural networks predict fMRI pattern-information along the ventral object processing pathway. Scientific Reports, 8(1), Article 1. 10.1038/s41598-018-28865-1

Diedrichsen, J., Berlot, E., Mur, M., Schütt, H. H., Shahbazi, M., & Kriegeskorte, N. (2021). *Comparing representational geometries using whitened unbiased-distance-matrix similarity* (arXiv:2007.02789). arXiv. 10.48550/arXiv.2007.02789

Diedrichsen, J., Yokoi, A., & Arbuckle, S. A. (2018). Pattern component modeling: A flexible approach for understanding the representational structure of brain activity patterns. NeuroImage, 180(Pt A), 119–133. 10.1016/j.neuroimage.2017.08.051

Dimsdale-Zucker, H. R., & Ranganath, C. (2018). Representational Similarity Analyses: A Practical Guide for Functional MRI Applications. In D. Manahan-Vaughan (Ed.), Handbook of Behavioral Neuroscience (Vol. 28, pp. 509–525). Elsevier. 10.1016/B978-0-12-812028-6.00027-6

Esteban, O., Markiewicz, C. J., Blair, R. W., Moodie, C. A., Isik, A. I., Erramuzpe, A., Kent, J. D., Goncalves, M., DuPre, E., Snyder, M., Oya, H., Ghosh, S. S., Wright, J., Durnez, J., Poldrack, R. A., & Gorgolewski, K. J. (2019). fMRIPrep: A robust preprocessing pipeline for functional MRI. Nature Methods, 16(1), Article 1. 10.1038/s41592-018-0235-4

Esteban, O., Markiewicz, C. J., Goncalves, M., Provins, C., Salo, T., Kent, J. D., DuPre, E., Ciric, R., Pinsard, B., Blair, R. W., Poldrack, R. A., & Gorgolewski, K. J. (2023). fMRIPrep: A robust preprocessing pipeline for functional MRI (Version 23.0.1) [Computer software]. Zenodo. 10.5281/zenodo.11123262

Fan, L., Li, H., Zhuo, J., Zhang, Y., Wang, J., Chen, L., Yang, Z., Chu, C., Xie, S., Laird, A. R., Fox, P. T., Eickhoff, S. B., Yu, C., & Jiang, T. (2016). The Human Brainnetome Atlas: A New Brain Atlas Based on Connectional Architecture. Cerebral Cortex, 26(8), 3508– 3526. 10.1093/cercor/bhw157

Favela, L. H., & Machery, E. (2023). Investigating the concept of representation in the neural and psychological sciences. Frontiers in Psychology, 14. 10.3389/fpsyg.2023.1165622

Favila, S. E., Lee, H., & Kuhl, B. A. (2020). Transforming the Concept of Memory Reactivation. Trends in Neurosciences, 43(12), 939–950. 10.1016/j.tins.2020.09.006

Freeman, J. B., Stolier, R. M., Brooks, J. A., & Stillerman, B. S. (2018). The neural representational geometry of social perception. *Current Opinion in Psychology*, Social Neuroscience, 24, 83–91. 10.1016/j.copsyc.2018.10.003

Freund, M. C., Bugg, J. M., & Braver, T. S. (2021). A Representational Similarity Analysis of Cognitive Control during Color-Word Stroop. Journal of Neuroscience, 41(35), 7388– 7402. 10.1523/JNEUROSCI.2956-20.2021

Freund, M. C., Etzel, J. A., & Braver, T. S. (2021). Neural Coding of Cognitive Control: The Representational Similarity Analysis Approach. Trends in Cognitive Sciences, 25(7), 622–638. 10.1016/j.tics.2021.03.011

Friston, K. J., Ashburner, J. T., Kiebel, S. J., Nichols, T. E., & Penny, W. D. (Eds.). (2006). Statistical parametric mapping: The analysis of functional brain images (1st ed.). Academic Press.

Greve, D. N., & Fischl, B. (2009). Accurate and robust brain image alignment using boundary-based registration. NeuroImage, 48(1), 63–72. 10.1016/j.neuroimage.2009.06.060

Grill-Spector, K., Henson, R., & Martin, A. (2006). Repetition and the brain: Neural models of stimulus-specific effects. Trends in Cognitive Sciences, 10(1), 14–23. 10.1016/j.tics.2005.11.006

Hebart, M. N., Contier, O., Teichmann, L., Rockter, A. H., Zheng, C. Y., Kidder, A., Corriveau, A., Vaziri-Pashkam, M., & Baker, C. I. (2023). THINGS-data, a multimodal collection of large-scale datasets for investigating object representations in human brain and behavior. eLife, 12, e82580. 10.7554/eLife.82580

Hovhannisyan, M., Clarke, A., Geib, B. R., Cicchinelli, R., Monge, Z., Worth, T., Szymanski, A., Cabeza, R., & Davis, S. W. (2021). The visual and semantic features that predict object memory: Concept property norms for 1,000 object images. Memory & Cognition, 49(4), 712–731. 10.3758/s13421-020-01130-5

Howard, C. M., Huang, S., Hovhannisyan, M., Cabeza, R., & Davis, S. W. (2024). Differential Mnemonic Contributions of Cortical Representations during Encoding and Retrieval. Journal of Cognitive Neuroscience, 1–29. 10.1162/jocn_a_02227

Huang, S., Bogdan, P. C., Howard, C. M., Gillette, K., Deng, L., Welch, E., McAllister, M. L., Giovanello, K. S., Davis, S. W., & Cabeza, R. (2025). Cortico-hippocampal interactions underlie schema-supported memory encoding in older adults. Cerebral Cortex, 35(8). 10.1093/cercor/bhaf211

Huang, S., De Brigard, F., Cabeza, R., & Davis, S. W. (2024). Connectivity analyses for task-based fMRI. Physics of Life Reviews, 49, 139–156. 10.1016/j.plrev.2024.04.012

Huang, S., Faul, L., Parikh, N., LaBar, K. S., & De Brigard, F. (2024). Counterfactual thinking induces different neural patterns of memory modification in anxious individuals. Scientific Reports, 14(1), 10630. 10.1038/s41598-024-61545-x

Huang, S., Howard, C. M., Hovhannisyan, M., Ritchey, M., Cabeza, R., & Davis, S. W. (2024). Hippocampal Functions Modulate Transfer-Appropriate Cortical Representations Supporting Subsequent Memory. Journal of Neuroscience, 44(1). 10.1523/JNEUROSCI.1135-23.2023

Huang, T., Song, Y., & Liu, J. (2022). Real-world size of objects serves as an axis of object space. Communications Biology, 5(1), 1–12. 10.1038/s42003-022-03711-3

Jenkinson, M., Bannister, P., Brady, M., & Smith, S. (2002). Improved Optimization for the Robust and Accurate Linear Registration and Motion Correction of Brain Images. NeuroImage, 17(2), 825–841. 10.1006/nimg.2002.1132

Jozranjbar, B., Kristjánsson, Á., Starrfelt, R., Gerlach, C., & Sigurdardottir, H. M. (2023). Using representational similarity analysis to reveal category and process specificity in visual object recognition. Cortex, 166, 172–187. 10.1016/j.cortex.2023.05.012

Jozwik, K. M., Kietzmann, T. C., Cichy, R. M., Kriegeskorte, N., & Mur, M. (2023). Deep Neural Networks and Visuo-Semantic Models Explain Complementary Components of Human Ventral-Stream Representational Dynamics. Journal of Neuroscience, 43(10), 1731– 1741. 10.1523/JNEUROSCI.1424-22.2022

Karimi-Rouzbahani, H., Woolgar, A., Henson, R., & Nili, H. (2022). Caveats and Nuances of Model-Based and Model-Free Representational Connectivity Analysis. Frontiers in Neuroscience, 16. https://www.frontiersin.org/article/10.3389/fnins.2022.755988

King, M. L., Groen, I. I. A., Steel, A., Kravitz, D. J., & Baker, C. I. (2019). Similarity judgments and cortical visual responses reflect different properties of object and scene categories in naturalistic images. NeuroImage, 197, 368–382. 10.1016/j.neuroimage.2019.04.079

Klein, A., Ghosh, S. S., Bao, F. S., Giard, J., Häme, Y., Stavsky, E., Lee, N., Rossa, B., Reuter, M., Neto, E. C., & Keshavan, A. (2017). Mindboggling morphometry of human brains. PLOS Computational Biology, 13(2), e1005350. 10.1371/journal.pcbi.1005350

Kleiner, M., Brainard, D., & Pelli, D. (2007). What’s new in Psychtoolbox-3? Perception, 36(14), 1–16.

Kramer, M. A., Hebart, M. N., Baker, C. I., & Bainbridge, W. A. (2023). The features underlying the memorability of objects. Science Advances, 9(17), eadd2981. 10.1126/sciadv.add2981

Kriegeskorte, N., & Diedrichsen, J. (2019). Peeling the Onion of Brain Representations. Annual Review of Neuroscience, 42(1), 407–432. 10.1146/annurev-neuro-080317-061906

Kriegeskorte, N., & Kievit, R. A. (2013). Representational geometry: Integrating cognition, computation, and the brain. Trends in Cognitive Sciences, 17(8), 401–412. 10.1016/j.tics.2013.06.007

Kriegeskorte, N., Mur, M., & Bandettini, P. A. (2008). Representational similarity analysis— Connecting the branches of systems neuroscience. Frontiers in Systems Neuroscience, 2. 10.3389/neuro.06.004.2008

Kuhn, M., Vaughan, D., & Hvitfeldt, E. (2024). yardstick: Tidy Characterizations of Model Performance (Version 1.3.1) [Computer software]. https://github.com/tidymodels/yardstick

Kuznetsova, A., Brockhoff, P. B., & Christensen, R. H. B. (2017). lmerTest Package: Tests in Linear Mixed Effects Models. Journal of Statistical Software, 82(13). 10.18637/jss.v082.i13

Lahner, B., Dwivedi, K., Iamshchinina, P., Graumann, M., Lascelles, A., Roig, G., Gifford, A. T., Pan, B., Jin, S., Ratan Murty, N. A., Kay, K., Oliva, A., & Cichy, R. (2024). Modeling short visual events through the BOLD moments video fMRI dataset and metadata. Nature Communications, 15(1), 6241. 10.1038/s41467-024-50310-3

Lanczos, C. (1964). Evaluation of Noisy Data. Journal of the Society for Industrial and Applied Mathematics Series B Numerical Analysis, 1(1), 76–85. 10.1137/0701007

Long, N. M., & Kuhl, B. A. (2021). Cortical Representations of Visual Stimuli Shift Locations with Changes in Memory States. Current Biology. 10.1016/j.cub.2021.01.004

MacCallum, R. C., Zhang, S., Preacher, K. J., & Rucker, D. D. (2002). On the practice of dichotomization of quantitative variables. Psychological Methods, 7(1), 19–40. (2002-00925-002). 10.1037/1082-989X.7.1.19

Matuschek, H., Kliegl, R., Vasishth, S., Baayen, H., & Bates, D. (2017). Balancing Type I error and power in linear mixed models. Journal of Memory and Language, 94, 305–315. 10.1016/j.jml.2017.01.001

Mazurchuk, S., Conant, L. L., Tong, J.-Q., Binder, J. R., & Fernandino, L. (2023). Stimulus repetition and sample size considerations in item-level representational similarity analysis. Language, Cognition and Neuroscience, 0(0), 1–12. 10.1080/23273798.2023.2232903

Meteyard, L., & Davies, R. A. I. (2020). Best practice guidance for linear mixed-effects models in psychological science. Journal of Memory and Language, 112, 104092. 10.1016/j.jml.2020.104092

Morales-Torres, R., Wing, E. A., Deng, L., Davis, S. W., & Cabeza, R. (2024). Visual Recognition Memory of Scenes Is Driven by Categorical, Not Sensory, Visual Representations. Journal of Neuroscience, 44(21). 10.1523/JNEUROSCI.1479-23.2024

Morton, N. W., Zippi, E. L., Noh, S. M., & Preston, A. R. (2021). Semantic Knowledge of Famous People and Places Is Represented in Hippocampus and Distinct Cortical Networks. Journal of Neuroscience, 41(12), 2762–2779. 10.1523/JNEUROSCI.2034-19.2021

Mumford, J. A., Turner, B. O., Ashby, F. G., & Poldrack, R. A. (2012). Deconvolving BOLD activation in event-related designs for multivoxel pattern classification analyses. NeuroImage, 59(3), 2636–2643. 10.1016/j.neuroimage.2011.08.076

Naselaris, T., Kay, K. N., Nishimoto, S., & Gallant, J. L. (2011). Encoding and decoding in fMRI. NeuroImage, 56(2), 400–410. 10.1016/j.neuroimage.2010.07.073

Naspi, L., Hoffman, P., Devereux, B., Thejll-Madsen, T., Doumas, L. A. A., & Morcom, A. (2021). Multiple dimensions of semantic and perceptual similarity contribute to mnemonic discrimination for pictures. *Journal of Experimental Psychology: Learning*, Memory, and Cognition, 47(12), 1903–1923. (2021-79539-001). 10.1037/xlm0001032

Naspi, L., Stensholt, C., Karlsson, A. E., Monge, Z. A., & Cabeza, R. (2023). Effects of Aging on Successful Object Encoding: Enhanced Semantic Representations Compensate for Impaired Visual Representations. Journal of Neuroscience, 43(44), 7337–7350. 10.1523/JNEUROSCI.2265-22.2023

Nili, H., Wingfield, C., Walther, A., Su, L., Marslen-Wilson, W., & Kriegeskorte, N. (2014). A Toolbox for Representational Similarity Analysis. PLOS Computational Biology, 10(4), e1003553. 10.1371/journal.pcbi.1003553

Pacheco-Estefan, D., Fellner, M.-C., Kunz, L., Zhang, H., Reinacher, P., Roy, C., Brandt, A., Schulze-Bonhage, A., Yang, L., Wang, S., Liu, J., Xue, G., & Axmacher, N. (2024). Maintenance and transformation of representational formats during working memory prioritization. Nature Communications, 15(1), 8234. 10.1038/s41467-024-52541-w

Park, J., Cardwell, R., & Yu, H.-T. (2020). Specifying the Random Effect Structure in Linear Mixed Effect Models for Analyzing Psycholinguistic Data. Methodology, 16(2), Article 2. 10.5964/meth.2809

Power, J. D., Mitra, A., Laumann, T. O., Snyder, A. Z., Schlaggar, B. L., & Petersen, S. E. (2014). Methods to detect, characterize, and remove motion artifact in resting state fMRI. NeuroImage, 84, 320–341. 10.1016/j.neuroimage.2013.08.048

Prince, J. S., Charest, I., Kurzawski, J. W., Pyles, J. A., Tarr, M. J., & Kay, K. N. (2022). Improving the accuracy of single-trial fMRI response estimates using GLMsingle. eLife, 11, e77599. 10.7554/eLife.77599

R Core Team. (2024). R: A Language and Environment for Statistical Computing (Version 4.4.1) [Computer software]. R Foundation for Statistical Computing. https://www.R-project.org/

Reuter, M., Rosas, H. D., & Fischl, B. (2010). Highly accurate inverse consistent registration: A robust approach. NeuroImage, 53(4), 1181–1196. 10.1016/j.neuroimage.2010.07.020

Riesenhuber, M., & Poggio, T. (1999). Hierarchical models of object recognition in cortex. Nature Neuroscience, 2(11), 1019–1025. 10.1038/14819

Riesenhuber, M., & Poggio, T. (2002). Neural mechanisms of object recognition. Current Opinion in Neurobiology, 12(2), 162–168. 10.1016/S0959-4388(02)00304-5

Ritchey, M., Wing, E. A., LaBar, K. S., & Cabeza, R. (2013). Neural Similarity Between Encoding and Retrieval is Related to Memory Via Hippocampal Interactions. Cerebral Cortex, 23(12), 2818–2828. 10.1093/cercor/bhs258

Ritchie, J. B., Lee Masson, H., Bracci, S., & Op de Beeck, H. P. (2021). The unreliable influence of multivariate noise normalization on the reliability of neural dissimilarity. NeuroImage, 245, 118686. 10.1016/j.neuroimage.2021.118686

Rothlein, D., DeGutis, J., & Esterman, M. (2018). Attentional Fluctuations Influence the Neural Fidelity and Connectivity of Stimulus Representations. Journal of Cognitive Neuroscience, 30(9), 1209–1228. 10.1162/jocn_a_01306

Satterthwaite, F. E. (1946). An Approximate Distribution of Estimates of Variance Components. Biometrics Bulletin, 2(6), 110–114. 10.2307/3002019

Schütt, H. H., Kipnis, A. D., Diedrichsen, J., & Kriegeskorte, N. (2023). Statistical inference on representational geometries. eLife, 12, e82566. 10.7554/eLife.82566

Serre, T., Oliva, A., & Poggio, T. (2007). A feedforward architecture accounts for rapid categorization. Proceedings of the National Academy of Sciences, 104(15), 6424–6429. 10.1073/pnas.0700622104

Serre, T., Wolf, L., & Poggio, T. (2005). Object recognition with features inspired by visual cortex. 2005 IEEE Computer Society Conference on Computer Vision and Pattern Recognition (CVPR’05), 2, 994–1000 vol. 2. 10.1109/CVPR.2005.254

Shao, X., Li, A., Chen, C., Loftus, E. F., & Zhu, B. (2023). Cross-stage neural pattern similarity in the hippocampus predicts false memory derived from post-event inaccurate information. Nature Communications, 14(1), Article 1. 10.1038/s41467-023-38046-y

Silver, N. C., & Dunlap, W. P. (1987). Averaging correlation coefficients: Should Fisher’s z transformation be used? Journal of Applied Psychology, 72, 146–148. 10.1037/0021-9010.72.1.146

Slayton, M., Howard, C. M., Huang, S., Hovhannisyan, M., Cabeza, R., & Davis, S. W. (2025). Semantic Dimensions Support the Cortical Representation of Object Memorability. Journal of Cognitive Neuroscience, 1–27. 10.1162/jocn.a.60

Sucholutsky, I., Muttenthaler, L., Weller, A., Peng, A., Bobu, A., Kim, B., Love, B. C., Cueva, C. J., Grant, E., Groen, I., Achterberg, J., Tenenbaum, J. B., Collins, K. M., Hermann, K., Oktar, K., Greff, K., Hebart, M. N., Cloos, N., Kriegeskorte, N., … Griffiths, T. L. (2025). Getting aligned on representational alignment. Transactions on Machine Learning Research. https://openreview.net/forum?id=Hiq7lUh4Yn

Sufikarimi, H., & Mohammadi, K. (2020). Role of the secondary visual cortex in HMAX model for object recognition. Cognitive Systems Research, 64, 15–28. 10.1016/j.cogsys.2020.07.001

Tustison, N. J., Avants, B. B., Cook, P. A., Zheng, Y., Egan, A., Yushkevich, P. A., & Gee, J. C. (2010). N4ITK: Improved N3 Bias Correction. IEEE Transactions on Medical Imaging, 29(6), 1310–1320. IEEE Transactions on Medical Imaging. 10.1109/TMI.2010.2046908

Tyler, L. K., Chiu, S., Zhuang, J., Randall, B., Devereux, B. J., Wright, P., Clarke, A., & Taylor, K. I. (2013). Objects and Categories: Feature Statistics and Object Processing in the Ventral Stream. Journal of Cognitive Neuroscience, 25(10), 1723–1735. 10.1162/jocn_a_00419

van der Wel, P., & van Steenbergen, H. (2018). Pupil dilation as an index of effort in cognitive control tasks: A review. Psychonomic Bulletin & Review, 25(6), 2005–2015. 10.3758/s13423-018-1432-y

Venables, W. N., Ripley, B. D., & Venables, W. N. (2002). *Modern applied statistics with S* (4th ed). Springer.

Vilarroya, O. (2017). Neural Representation. A Survey-Based Analysis of the Notion. Frontiers in Psychology, 8. 10.3389/fpsyg.2017.01458

Walsh, C. R., & Rissman, J. (2023). Behavioral representational similarity analysis reveals how episodic learning is influenced by and reshapes semantic memory. Nature Communications, 14(1), Article 1. 10.1038/s41467-023-42770-w

Walther, A., Nili, H., Ejaz, N., Alink, A., Kriegeskorte, N., & Diedrichsen, J. (2016). Reliability of dissimilarity measures for multi-voxel pattern analysis. NeuroImage, 137, 188–200. 10.1016/j.neuroimage.2015.12.012

Weaverdyck, M. E., Lieberman, M. D., & Parkinson, C. (2020). Tools of the Trade Multivoxel pattern analysis in fMRI: A practical introduction for social and affective neuroscientists. Social Cognitive and Affective Neuroscience, 15(4), 487–509. 10.1093/scan/nsaa057

Westfall, J., Nichols, T. E., & Yarkoni, T. (2017). Fixing the stimulus-as-fixed-effect fallacy in task fMRI. Wellcome Open Research, 1, 23. 10.12688/wellcomeopenres.10298.2

Wing, E. A., Geib, B. R., Wang, W.-C., Monge, Z., Davis, S. W., & Cabeza, R. (2020). Cortical Overlap and Cortical-Hippocampal Interactions Predict Subsequent True and False Memory. Journal of Neuroscience, 40(9), 1920–1930. 10.1523/JNEUROSCI.1766-19.2020

Winter, B., & Grice, M. (2021). Independence and generalizability in linguistics. Linguistics, 59(5), 1251–1277. 10.1515/ling-2019-0049

Xue, G. (2018). The Neural Representations Underlying Human Episodic Memory. Trends in Cognitive Sciences, 22(6), 544–561. 10.1016/j.tics.2018.03.004

Yarkoni, T. (2022). The generalizability crisis. Behavioral and Brain Sciences, 45, e1. 10.1017/S0140525X20001685

Yu, C., Huang, S., Howard, C. M., Hovhannisyan, M., Clarke, A., Cabeza, R., & Davis, S. W. (2024). Subsequent Memory Effects in Cortical Pattern Similarity Differ by Semantic Class. Journal of Cognitive Neuroscience, 1–12. 10.1162/jocn_a_02238

Zhang, Y., Brady, M., & Smith, S. (2001). Segmentation of brain MR images through a hidden Markov random field model and the expectation-maximization algorithm. IEEE Transactions on Medical Imaging, 20(1), 45–57. 10.1109/42.906424

